# Knockdown of LC3 increases mitochondria-to-micronucleus transition

**DOI:** 10.1101/2020.11.06.372144

**Authors:** Bolin Hou, Erwei Li, Haiwen Huang, Huaiyi Yang, Zhijun Xi, Xuejun Jiang

**Author notes:** These authors contributed equally to this work (Bolin Hou, Erwei Li). Correspondence to: Xuejun Jiang and Zhijun Xi.

## Abstract

Nuclear-localized mitochondria were discovered over sixty years ago^1^; however, the function of these organelles in the initiation of nuclear formation and development remains unknown. Here, we showed that mitochondria fragmented into dense particles to initiate and develop a nucleus, and multiple nuclei were separately and simultaneously formed by fragmented mitochondria in a single cell. The combination of nuclei individually constructed by the mitochondrial assembly of dense particles for nuclear transition partitioned the cytoplasm to form an intranuclear inclusion (INC), whose formation was not related to herniation or invagination of the cytoplasm. During nuclear conversion of itself and neighbouring organelles, the mitochondrion was incorporated into the nucleus to become a nuclear mitochondrion. Knockdown of microtubule-associated protein light chain 3 (LC3), a key autophagic protein, increased free micronuclei by delaying nuclear fusion and enhancing the mitochondria-to-micronuclei transition.

---

Before the term nuclear mitochondria was introduced^2^, cytoplasmic organelles such as mitochondria, endoplasmic reticulum (ER)^3, 4^ and Golgi complex were identified in INCs^5, 6^, which can be either large or small and either opened or closed^3, 7^. Thereafter, nuclear mitochondria were found in primary cells^8, 9^, human tissues^10, 11^ and the cells with ruptured nuclear envelop^12^. Actually, the INC was discovered over a century ago^13^, while the exact mechanism of its formation remains unclear and elusive until now.

Micronuclei are small membrane-enveloped structures separate from the primary nucleus of the cell that can contain chromosomal fragments as well as whole chromosomes^14^. Usually, micronuclei are believed to be generated in response to genome damage by various causes^15, 16^; they are widely accepted diagnostic markers for genomic instability and are often used to screen chemicals for genotoxicity^17^. Loss of autophagy has been linked to increased nuclear instability and leads to the disruption of mitochondrial quality^18–20^.

By transmission electron microscopy (TEM), we observed that one or more micronuclei formed separately in a single K562 cell, which already contained a large or primary nucleus (PN), whereas partial or completely fragmented mitochondria existed between nuclei (Fig. 1a and Extended data Fig. 1). Through the assembly of dense particles, mitochondria fragmented into particles for nuclear development, concomitantly attaching a micronucleus (MN) to the PN (Fig. 1b and Extended data Fig. 2). In addition to an MN, the mitochondrial aggregation of dense particles achieved nuclear transition and fusion to link the individually built large nuclei (Fig. 1c and Extended data Fig. 3), while partial nuclear merging induced by mitochondrial fragmentation partitioned the cytoplasm to form an INC, which was either opened or closed (Fig. 1d and Extended data Fig. 4). Notably, the mitochondrial assembly of dense particles into the nucleolus (Nu) was observed (Fig. 1d), whereas INC formation was merely due to incomplete mitochondria-to-nucleus transition but not related to herniation or invagination of the cytoplasm.

**Fig. 1.**
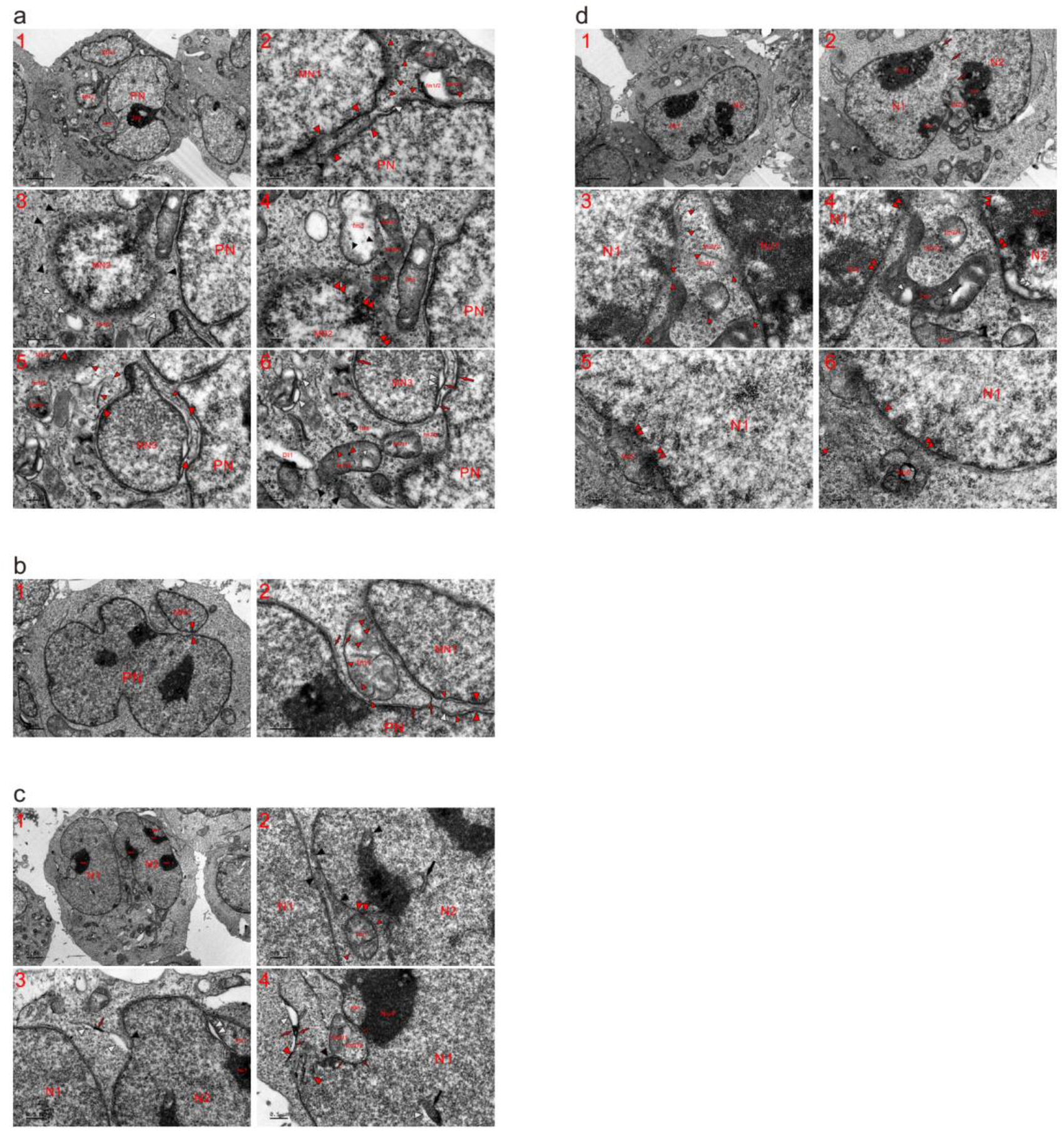
Mitochondria completely fragmented into dense particles to achieve nuclear transition and fusion. Transmission electron microscopy (TEM) was performed on four K562 cells following 2 h of incubation in new medium. (**a**) In addition to a large nucleus (PN), three micronuclei (MN1-MN2) were being built and attached to the PN by fragmented mitochondria. (**b**) MN1 was fusing with the PN (opposite large red arrowheads) via the further aggregation of dense particles (large white and small red arrowheads), whose assembly enlarged both nuclei (PN and MN1), leading the less fragmented mitochondrion (fm1) to localize between the PN and MN1. (**c**) Two individually formed large nuclei (N1 and N2) were merged by fragmented mitochondria (black arrowheads), and mitochondrial assembly of dense particles or incomplete mitochondria-to-nucleus transition was displayed at the edge of Nu2. fm1 diffused into N2 (double red arrowheads) and the surroundings in the form of dense particles (small red arrowheads). (**d**) The nuclei (N1 and N2) joined together to form a large nucleus and partial nuclear fusion partitioned the cytoplasm to form INC1, in which less fragmented mitochondria (such as fm1-fm3) were dispersing into dense particles (small red arrowheads), while fm2/2 and fm3/2 were diffused into the particles. Condensed fm1 assembled dense particles (small red arrowheads) into N1 and nucleoli (Nu1 and Nu2) (double red arrowheads), and their aggregation formed electron-transparent spots in the organelle (white arrowheads).

In either the nucleus or the partitioned cytoplasm (or an INC), the organelles assembled dense particles for nuclear development or growth (Fig. 2a and Extended data Fig. 5). We found that the incomplete nuclear conversion of one or more mitochondria led to the formation of a small INC at the nuclear edge and within the nucleus (Fig. 2b and Extended data Fig. 6). In the nucleus and at its edge, mitochondria exhibited both internal and external aggregation of dense particles (Fig. 2c and Extended data Fig. 7), whose assembly at the peripheries (external assembly) dispersed mitochondria into the nucleus and caused parts of the organelles to become electron transparent (Fig. 2d and Extended data Fig. 8). At a relatively early time point (4 h), mitochondria were usually electron-opaque and directly fragmented into a young nucleus (YN)^21^ in the form of dense particles (Extended data Fig. 9a); in contrast, external assembly markedly increased to aggregate the particles into the YN for nuclear enlargement at the 12 h time point, accompanied by the accumulation of vesicles in the cytoplasm (Extended data Fig. 9b).

**Fig. 2.**
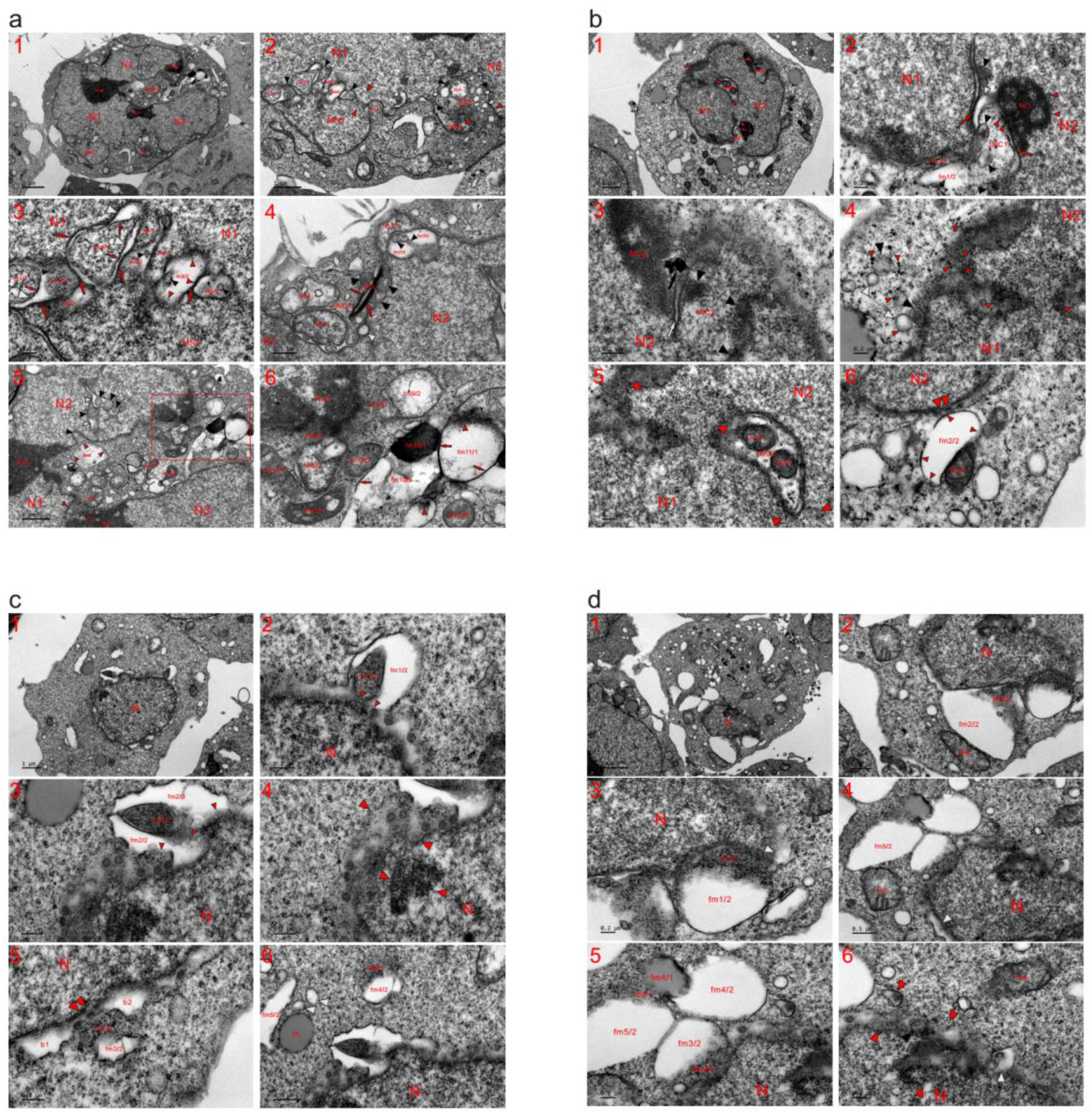
Mitochondrial externally and internally assembled dense particles for nuclear development to build a large nucleus. TEM was performed on four K562 cells at the 2 (**a**) and 12 (**b-d**) h time points, and the micrographs revealed that mitochondria performed internal and external assembly of dense particles to build and develop a large nucleus. (**a**) Combination of separately built nuclei (N1-N3) compartmentalized the cytoplasm to form INC1, and the individually formed MN1 was merged with N1 by fragmented mitochondria (fm1 and m1-m3; opposite red, black and white arrowheads). MN2 was built by the mitochondrial assembly of dense particles, and the formation of the micronucleus together with N3 partitioned the cytoplasm to shape INC2, which was filled with fragmented mitochondria (such as fm2, black and white arrowheads). Mitochondrial fragmentation led to the formation of SDBs (black arrowheads) and SDIBs (white arrowheads), some of which look like vesicles. At the opening of INC2, SDBs and SDIBs were becoming dense particles for nuclear development to seal the INC. (**b**) Mitochondria-to-nucleus transition (opposite red arrowheads) recently occurred to accomplish partial fusion between individually constructed nuclei (N1 and N2); following the nuclear development of mitochondria, the area of the cytoplasm was consequently decreased and partitioned by the nuclei to form intranuclear inclusions (INC1-INC3). (**c**) A nucleus (N) was constructed by fragmented mitochondria, which surrounded and assembled dense particles into the nucleus. Mitochondria (fm1 and fm2) assembled the particles both internally (fm1/1 and fm2/1) and externally (red arrows and small red arrowheads) to disperse into the nucleus (small red arrowheads), causing the organelles to become electron transparent (fm1/2, fm2/2 and fm2/3). (**d**) A small or young nucleus (YN) appeared in this cell, and the mitochondria assembled dense particles (such as fm1/1-fm3/1) into the nucleus and concurrently became electron-transparent (such as fm1/2-fm3/2). The aggregation of dense particles separated fm4 into lucent fm4/2 and dense fm4/1, which looked like a lipid droplet and fused with fm5/1. At the nuclear edge, the mitochondria assembled the particles to diffuse into the YN (opposite red, black and white arrowhead).

In cells with no recognizable nucleus, the organelles completely fragmented into dense particles, which dispersed among each other (Extended data Fig. 10a) or assembled to form groups (Extended data Fig. 10b and c). We found that mitochondria assembled dense particles to initiate the formation of the nascent nucleus (NN) (Fig. 3a and Extended data Fig. 10d), which occasionally looked like a dispersed mitochondrion (Fig. 3a). The NN later developed the appearance of a YN by the continuous congregation of dense particles of the organelles; as mentioned above, the mitochondrial assembly of the particles appeared within the nucleus as well as at its edge (Extended data Fig. 11). It was not a rare phenomenon for mitochondria to form an NN in a cell that already possessed a PN (or a large nucleus) (Fig. 3b and Extended data Fig. 12). In contrast to what happened at the 2 h time point, it was much easier to observe that mitochondria-derived dense particles formed virus-like granules (VLGs, 70-120 nm in diameter) around an NN at both the 8 and 12 h time points (Extended data Fig. 12a-c). Through the assembly of dense particles, a single mitochondrion developed a nuclear appearance and appeared as a small MN (Fig. 3c and Extended data Fig. 13), whereas mitochondrial aggregation of the particles initially formed an electron-opaque body, which tended to evolve into an MN via further assembly of the enclosed dense particles (Fig. 3d and Extended data Fig. 14). Along with the external assembly of mitochondrial particles for nuclear growth, the electron-transparent structures were connected to form a lunar halo around an MN or a large nucleus (Fig. 3c and d). In addition to K562 cells, the involvement of mitochondria in the initiation of nuclear formation and development was also observed in all adherent cell lines (HepG2, HeLa and HEK293T) that had been examined (Extended data Figs. 15-17).

**Fig. 3.**
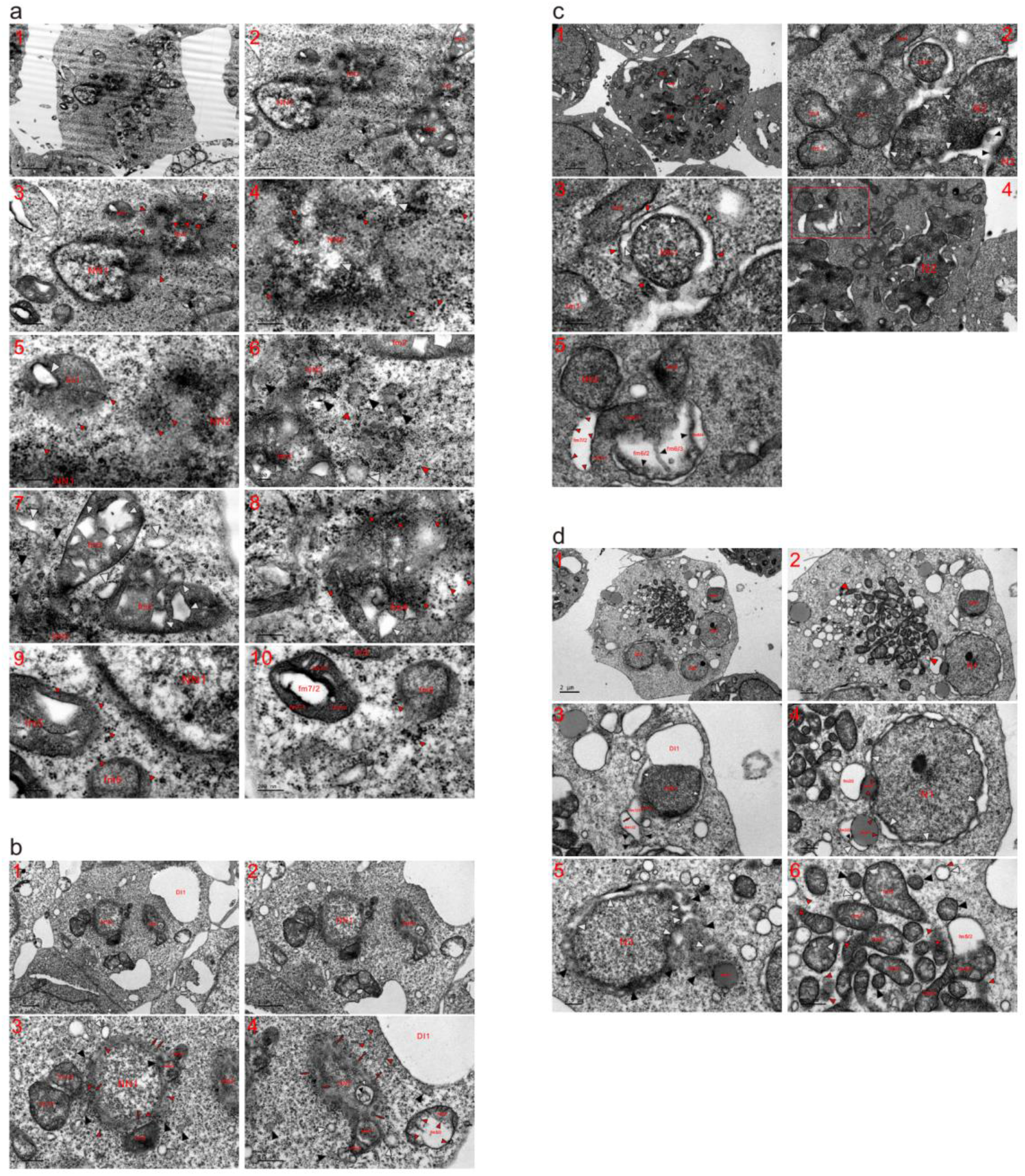
Initiation of nuclear formation began with mitochondrial fragmentation into dense particles. TEM was performed on four K562 cells at the 2 (**a**), 8 (**b**) and 12 (**c** and **d**) h time points, and the micrographs showed that mitochondria fragmented into dense particles to initiate the formation of a nascent nucleus (NN). (**a**) Three nascent nuclei (NN1-NN3) appeared in this cell, and NN2 was formed by the mitochondrial assembly of dense particles (opposite red arrowheads), whose aggregation fused NN2 with NN1. Mitochondria completely fragmented into dense particles (small red arrowheads) to enlarge NN2, concurrently creating electron-transparent spots in the nascent nucleus (white arrowheads). Partially fragmented fm1 was dispersed into dense particles (small red arrowheads), which entered both NN1 and NN2, and the external aggregation to the periphery produced an electron-lucent body in the mitochondrion (white arrowheads). (**b**) In addition to a YN, a nascent nucleus was built by the mitochondrial assembly of dense particles, and partially (such as fm1-fm3) or completely (white arrows, and black, white and small red arrowheads) fragmented mitochondria surrounded the YN to promote nuclear growth. The internal aggregation of dense particles (red arrows) separated mitochondria (fm2/1 and fm2/2; fm3/1 and fm3/2). At the nuclear edges, virus-like granules (VLGs) (white arrows) were transiently derived from mitochondrial fragmentations. (**c**) Mitochondria (or mitochondrion)-to-nucleus transition and thereafter nuclear fusion continuously and repeatedly occurred to build two large nuclei in the cell; mitochondrial aggregations of dense particles (black arrowheads) for nuclear formation created electron-lucent structures at the nuclear edges (white arrowheads) and caused mitochondria to disperse (such as fm1-fm4). Between the internal (MN1) and external (small red arrowheads) congregation, a lunar halo structure appeared, in which dispersion or aggregation of the particles was observed (white arrowheads). (**d**) Three nuclei of similar size (N1-N3) separately and simultaneously formed in the cell, and adjacent to N1, a group of condensed mitochondria appeared (opposite red arrowheads). Mitochondrial aggregations of dense particles formed a nascent nucleus (NN1) and led to the formation of electron-transparent structures (DI1 and white arrowheads).

Autophagy plays a critical role in both maintaining nuclear stability and controlling mitochondrial quality^22–24^. To examine whether autophagy plays a regulatory role in the mitochondria-to-nucleus transition, we knocked down LC3 in K562 cells and found that LC3 loss markedly reduced the amount of cytochrome C oxidase subunit IV (COX-IV) in the soluble nuclear fraction (Extended Data Fig. 18). The TEM results demonstrated that LC3 deprivation obviously hindered nuclear fusion, preventing individually formed nuclei from joining together (Extended data Fig. 19). Compared to the control cells, LC3 silencing led to the accumulation of free micronuclei (Fig. 4a and Extended data Fig. 20). In the control cells and following nuclear development of the neighbouring counterparts, a mitochondrion became embedded between nuclei and exhibited similar density to that of the nucleus, while mitochondria between the widely separated nuclei were usually heavily or completely fragmented and lacked energy in LC3-depleted cells (Fig. 4b and Extended data Fig. 21). Furthermore, LC3 knockdown appeared to enhance the mitochondria-to-micronucleus transition in LC3-deprived cells (Fig. 4c and Extended data Fig. 22). While nascent nuclei were connectedly and simultaneously formed by fragmented mitochondria in control cells, three or more nuclei separately and concurrently formed in LC3-silenced cells (Fig. 4d and Extended data Fig. 23).

**Fig. 4.**
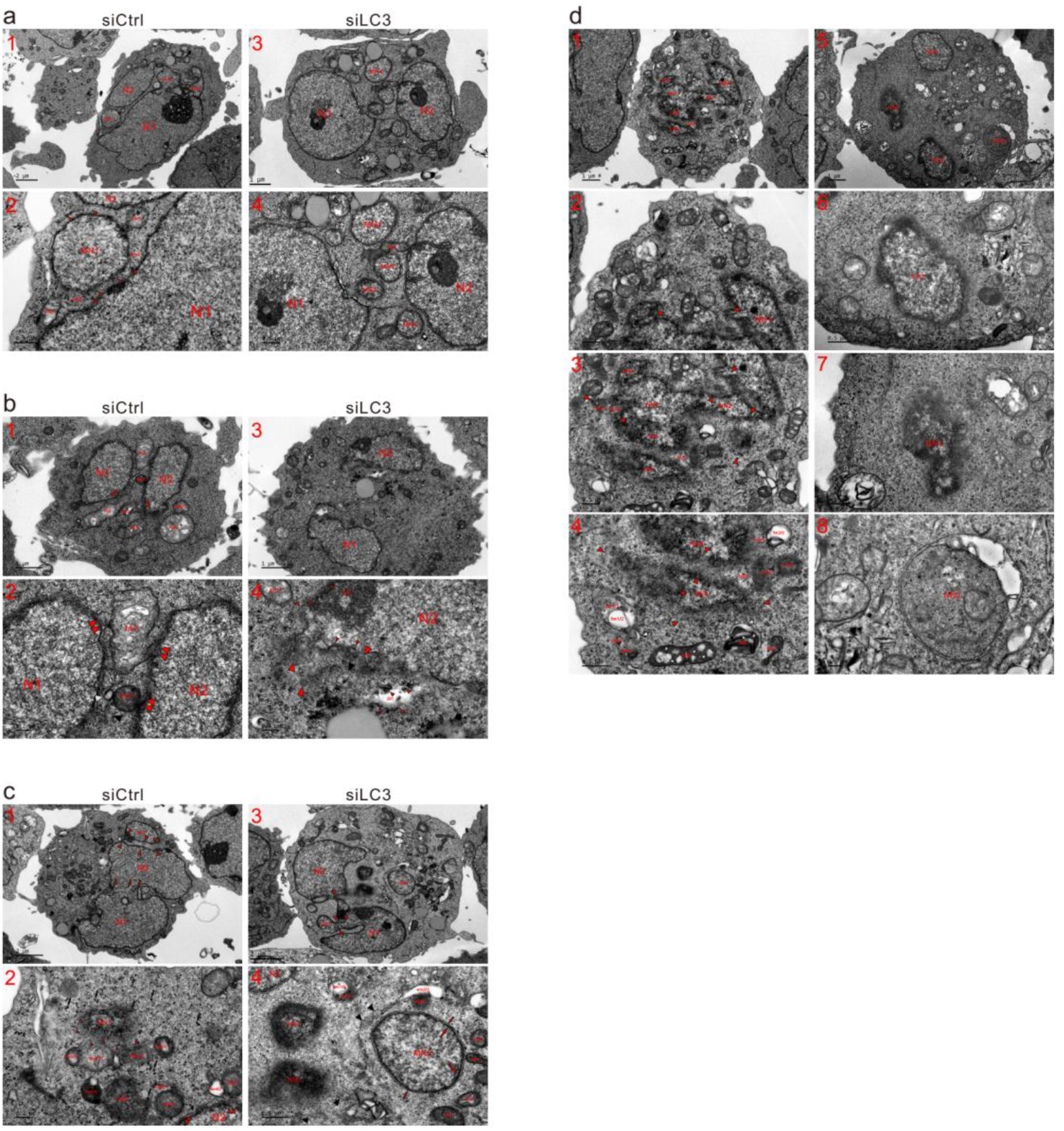
Knockdown of LC3 increased free micronuclei by hindering nuclear fusion and enhancing mitochondria-to-micronucleus transition. K562 cells were transfected with either small interfering RNA (siRNA) against LC3 (siLC3) or control siRNA (siCtrl) for 48 h, and the cells were gathered and cultured in fresh medium for 2 h before TEM observation. (**a**) In this control cell, MN1 was fused with the large nuclei (N1 and N2) by partially (fm1 and fm2) or completely (such as fm3 and fm4; small red arrowheads) fragmented mitochondria, while both MN2 and MN3 were already attached to N1 (**a1** and **a2**). (**a3** and **a4**) Compared to the large nuclei in control cells, N1 and N2 in this LC3-depleted cell were widely separated, and four micronuclei (MN1-MN4) existed between the large nuclei, while MN5 was relatively far away from N1. fm1 dispersed into N1 and micronuclei, and both fm2 and fm3 changed to a similar density to the nuclei. (**b**) Along with the neighbouring counterparts that had achieved nuclear transition to enlarge both N1 and N2, a mitochondrion (fm1) became embedded between the nuclei and developed a nuclear appearance (**b1** and **b2**). (**b3** and **b4**) Mitochondrial fragmentation was enhanced, and nuclei (N1 and N2) were widely separated in this LC3-silenced cell. (**c**) Mitochondria-to-nucleus transition led to partial nuclear fusion between two large nuclei (N1 and N2; three abreast red arrows), and MN1 was attached to N2 by the mitochondrial assembly of dense particles (opposite red arrowheads), whose aggregation (small red arrowheads) formed a nascent nucleus (NN1). (**c1** and **c2**). (**c3** and **c4**) In this LC3-depleted cell, the mitochondrial aggregation of dense particles attached MN1 to N1 and joined together two large nuclei (N1 and N2) (opposite red arrowheads), while MN2 was widely separated from the large nuclei. (**d**) In this control cell, seven nascent nuclei (NN1-NN7) appeared, which were connected and surrounded by partially or completely fragmented mitochondria (opposite red arrowheads), and NN7 looked like a mitochondrion and linked two nascent nuclei (NN5 and NN6) (**d1-d4**). (**d5-d8**) In the LC3-deprived cell, four nuclei (YN1, YN2, NN1 and NN2) were separately formed and/or enlarged.

The data presented here showed that mitochondria actively participated in the initiation of nuclear formation and consequent nuclear growth. The Individually formed nuclei fused by the mitochondrial aggregation of dense particles to construct a large nucleus in a single cell, and a combination of separately built nuclei partitioned the cytoplasm to create an INC, whose formation was merely due to incomplete mitochondria-to-nucleus transition but not related to herniation or invagination of the cytoplasm. Whether the organelle localized in an INC or appeared as a nuclear mitochondrion was strictly of cytoplasmic origin, and its discovery in the nucleus was merely due to incomplete nuclear conversion. Autophagy appeared to function to mediate the nuclear development of the organelles, and disruption of the autophagic process enhanced the mitochondria-to-micronucleus transition and hindered nuclear fusion.

## Methods

### Antibodies and siRNA

Antibodies against PARP-1 (9542) were purchased from Cell Signaling Technology (Beverly, MA, U.S.A.). Polyclonal antibodies against LC3 (L7543) were purchased form Sigma-Aldrich (St. Louis, MO, U.S.A.). Antibodies against LaminB1 (12987-1-AP) and COXIV (11242-1-AP) were purchased from Proteintech (Wuhan, China). Antibodies against actin (TA-09) were obtained from ZhongShanJinQiao Biocompany (Beijing, China). Antibodies against COX-IV (COX-4) (PA5-17511) were purchased from Invitrogen (Wyman Street, MA, U.S.A.).

The siRNA specific for human LC3 was purchased from Santa Cruz Biotechnology (sc-43390) and Dharmacon (L-012846-00-0020), along with the control siRNA (sc-37007).

### Cell culture

K562 (a myeloid leukemia-derived human cell line), HepG2, HeLa and HEK293T cells were cultured in DMEM (HyClone; SH20022.01B) containing 1% antibiotics and 10% fetal bovine serum (GIBCO; 16000)^25^. The culture durations of K562 cells ranged from 2 to 12 h, and the incubation times of HepG2, HeLa and HEK293T cells ranged from 2 to 6 h.

### Electron microscopy

The cells were harvested by centrifugation with (HepG2, HeLa and HEK293T cells) or without (K562 cells) trypsinization, washed three times with cold PBS and fixed with ice-cold glutaraldehyde (3% in 0.1 M cacodylate buffer, pH 7.4) for 30 minutes. After washing in PBS, the cells were postfixed in 1% OsO4 at room temperature (RT) for 1 h and dehydrated stepwise with ethanol. The dehydrated pellets were rinsed with propylene oxide at RT for 30 min and embedded in Spurr resin; 0.1 mm thin sections were stained with uranyl acetate/lead citrate (Fluka) and viewed under a JEM-1400 electron microscope (JEOL, Japan)^26^.

### Small RNA interference (siRNA)

K562 cells were grown in their respective media without antibiotics and transfected with siRNA against LC3 or control siRNA using lipofectamine RNAiMAX (Invitrogen) according to the manufacturer’s instructions. After 48 h transfection, cells were split and cultured with new medium for 2 h prior to electron microscopy observation and immunoblotting analysis^27^.

### Immunoblotting

Whole cell lysates were prepared via lysis using Triton X-100/glycerol buffer containing 50 mM Tris-HCl, 4 mM EDTA, 2 mM EGTA, and 1 mM dithiothreitol (pH 7.4), supplemented with 1% Triton X-100 and protease inhibitors, separated on an SDS-PAGE gel (13% or 8%, according to the molecular weights of the proteins of interest), and transferred to a PVDF membrane. Immunoblotting was performed using appropriate primary antibodies and suitable horseradish peroxidase-conjugated secondary antibodies, followed by detection with enhanced chemiluminescence (Pierce Chemical). A chemiluminescence gel imaging system (Tanon-5200, Tanon, Shanghai, China) was used to collect the chemiluminescence signals^28^.

### Subcellular fractionation

After transfection with siRNA of LC3 or control siRAN for 48 h, K562 cells were split and cultured with new medium for 2 h, Cells were collected, pelleted and washed three times with cold PBS. Cells (10%) were resuspended in Triton X-100/glycerol buffer and labeled whole cell lysate (WCL); the others were resuspended in 500 μl homogenization buffer A (10 mM HEPES-KOH [pH 7.9], 10 mM KCl, 1.5 mM MgCl2, 0.5 mM PMSF and 0.5 mM dithiothreitol) containing 0.5% NP-40, and then the homogenate was centrifuged at 3000 rpm for 5 min at 4°C after static incubation on ice for 15 min. After washing twice with buffer A without NP-40, the pellet was resuspended in 200 μl buffer C (20 mM Hepes-KOH [pH 7.9], 600 mM KCl, 1.5 mM MgCl2, 0.2 mM and 25% glycerol). Following rotation in a cold room (4 °C) for 15 min, the homogenate was equally divided in half. Half of the homogenate continued to be centrifuged at 13000 rpm for 15 min at 4 °C; the supernatant was collected as the soluble nuclear fraction (S-Nu), and the insoluble precipitate (Ins-Nu) was collected by directly adding 100 μl × loading buffer. The other half was subjected to proteinase (PK) treatment before centrifugation to separate S-Nu and Ins-Nu. WCL and nuclear fractions were electrophoresed on SDS-PAGE for immunoblotting analysis^27^.

### Proteinase K treatment

The nuclear fractions were treated with 200 ng/mL PK in buffer C without protease inhibitors at RT for 30 min. The reaction was stopped by the addition of 1 mM PMSF^29^.

## Acknowledgements

This work was supported by grants from the Scientific Development Fund of Institute of Microbiology, Chinese Academy of Sciences (Y954FJ1016) and the National Natural Science Foundation of China (81872771).

## Author contributions

Z.X. and X.J. designed the study and wrote the manuscript. B.H. and E.L. conducted the research and performed the Electron microscopy observation. Also B.H. and E.L. prepared the cell samples for immunoblot assays and subcellular fractionation. H.H. prepared Electron microscopy samples. Y.H. helped to polish the manuscript and analyzed results. All authors read and approved the final manuscript.

## Competing Interest

The authors declare that they have no conflict of interest.

## Additional information

## Extended Data and materials availability

All the data are available in the manuscript or the supplementary materials. Materials are available from X.J. on request.

## Notes

### Competing Interest Statement

The authors have declared no competing interest.

