## Supplemental Information for "Knockdown of LC3 increases mitochondria-to-micronucleus transition"

**This PDF file includes:**

**Detailed figure legends 1 to 4**

**Extended data Figs 1 to 23**

**Detailed figure legends**

**Fig. 1. Mitochondria completely fragmented into dense particles to achieve nuclear transition and fusion.** Transmission electron microscopy (TEM) was performed on four K562 cells following 2 h of incubation in new medium. (**a**) In addition to a large nucleus (PN), three micronuclei (MN1-MN2) were being built and attached to the PN by fragmented mitochondria. Between nuclei (MN1 and PN) and at the nuclear edge, partially (such as fm1 and fm2) and completely (black, white and opposite red arrowheads) fragmented mitochondria turned into dense particles (small red arrows) for nuclear growth to combine MN1 and PN. The aggregation of dense particles separated fm1 (fm1/1 and fm1/2), caused fm1/2 to become electron-transparent, and dispersed condensed fm2 into dense particles (**a1** and **a2**). (**a3** and **a4**) Mitochondria (fm3-fm5) assembled dense particles into MN2 (double red arrowheads) and heavily or completely fragmented to surround the micronucleus (such as fm6/2, black and white arrowheads). The external assembly of the particles to the periphery (small black arrowheads) allowed fm3 to become electron-lucent, and their aggregation separated (fm4/1-fm4/3) and dispersed dense fm4, while fm5 diffused into the PN (double red arrowheads), fm4 and the surroundings. (**a5** and **a6**) MN3 was being attached to the PN by the mitochondrial assembly of dense particles (fm7/1, fm7/2 and opposite red arrowheads). Fragmented mitochondria at the edge of MN3 and between nuclei (MN2 and MN3) (such as fm7-fm9; opposite white arrowheads) became particles (small red arrowheads). Dense fm7/1 aggregated the particles to both nuclei (PN and MN3) (double red arrowheads), and dilute fm7/2 dispersed into dense particles, whose aggregation formed dark strands (large red arrows) or threads (small red arrows) and led to the formation of electron-lucent structure within MN3 (double white arrowheads). Along with the aggregation of dense particles (small red arrowheads), electron-opaque fm10 was being dispersed and created a transparent spot (white arrowhead) within the organelle. Mitochondrial assembly of the particles formed electron-lucent DI1, dense bodies (DE1-DE4) or small dense bodies (SDBs) (black arrowheads) and small dilute bodies (SDIBs) (white arrowheads), all of which ultimately dispersed into dense particles. fm: fragmented mitochondrion; Nu: nucleolus; PN: primary nucleus; MN: micronucleus. (**b**) MN1 was fusing with the PN (opposite large red arrowheads) via the further aggregation of dense particles (large white and small red arrowheads), whose assembly enlarged both nuclei (PN and MN1), leading the less fragmented mitochondrion (fm1) to localize between the PN and MN1. Through congregation of the particles (small red arrowheads), fm1 became fused with nuclei (double red arrowheads) and transiently blackened its LMM (red arrow). The aggregation and linearization of mitochondria-derived dense particles intermediately formed dark threads, and both the dark threads and the LMM disappeared with further assembly of the particles. (**c**) Two individually formed large nuclei (N1 and N2) were merged by fragmented mitochondria (black arrowheads), and mitochondrial assembly of dense particles or incomplete mitochondria-to-nucleus transition was displayed at the edge of Nu2. fm1 diffused into N2 (double red arrowheads) and the surroundings in the form of dense particles (small red arrowheads) (**c1** and **c2**). (**c3**) The aggregation of dense particles (red arrow and black arrowhead) dispersed the organelles, forming electron-lucent phases or structures (white arrowheads) in the cytoplasm, which were partitioned by N1 and N2. Mitochondria-derived particles assembled to expand the sizes of both nuclei (N2 and MN1) and to form Nu1, causing MN1 to become an attached micronucleus. The remaining part of a fragmented mitochondrion formed a nuclear tubule (white arrow) and an endoplasmic reticulum (ER)-like structure (ERL) (double white arrowheads), representing incomplete nuclear transition of the organelle. (**c4**) At the edge of N1, the aggregation of dense particles separated fm2 (dense fm2/1 and dilute fm2/2), and their further assembly in fm2/2 formed dark threads (small red arrowheads), whose formation caused fm2 to look like a small and shallow INC at the nuclear edge. Within N1 and at its edge, the mitochondrial assembly of particles for nuclear development was observed (black arrow and opposite red, black and white arrowheads), and their aggregation enlarged Nu4, concurrently forming a nuclear body (NB1). Adjacent to N1 and in the cytoplasm, the mitochondrial congregation of dense particles or vestiges of mitochondrial fragmentations was demonstrated (large and small red arrows and black and white arrowheads). (**d**) The nuclei (N1 and N2) should have been separately formed and joined together to form a large nucleus, as trace of nuclear fusion were observed (three abreast red arrows). Partial nuclear fusion partitioned the cytoplasm to form INC1, in which less fragmented mitochondria (such as fm1-fm3) were dispersing into dense particles (small red arrowheads), while fm2/2 and fm3/2 were diffused into the particles. Condensed fm1 assembled dense particles (small red arrowheads) into N1 and nucleoli (Nu1 and Nu2) (double red arrowheads), and their aggregation formed electron-transparent spots in the organelle (white arrowheads); dense fm4 dispersed into fm1 and the surroundings, and the external mitochondrial assembly of the particles to the periphery incorporated the organelle in N2, leading part of a mitochondrion to become a dilute nuclear body (double white arrowheads) (**d1**-**d4**). (**d5** and **d6**) At the edge of N1, the mitochondrial morphology of both fm5 and fm6 was recognized, while the mitochondria (fm5, fm6 and opposite red arrowheads) assembled dense particles to disperse into N1 (double red arrowheads) for nuclear growth.

**Fig. 2. Mitochondrial externally and internally assembled dense particles for nuclear development to build a large nucleus.** TEM was performed on four K562 cells at the 2 (**a**) and 12 (**b-d**) h time points, and the micrographs revealed that mitochondria performed internal and external assembly of dense particles to build and develop a large nucleus. (**a**) Combination of separately built nuclei (N1-N3) compartmentalized the cytoplasm to form INC1, and the individually formed MN1 was merged with N1 by fragmented mitochondria (fm1 and m1-m3; opposite red, black and white arrowheads). MN2 was built by the mitochondrial assembly of dense particles, and the formation of the micronucleus together with N3 partitioned the cytoplasm to shape INC2, which was filled with fragmented mitochondria (such as fm2, black and white arrowheads). Mitochondrial fragmentation led to the formation of SDBs (black arrowheads) and SDIBs (white arrowheads), some of which look like vesicles. At the opening of INC2, SDBs and SDIBs were becoming dense particles for nuclear development to seal the INC (**a1** and **a2**). (**a3**) Along with the nuclear transition of their neighbouring counterparts to expand both MN1 and N1, mitochondria were included between the nuclei, and the internal aggregation of dense particles caused parts of the organelles to adopt a nuclear appearance (such as m1/1-m4/1). Through further assembly of the particles, the remaining parts achieved nuclear transition (m1/2 and m4/2-m6/2). Their external aggregation (small red arrowheads) separated m3 (m3/1 and m3/2), with m3/2 becoming electron-lucent through the external and internal (small black arrowhead) assembly of dense particles. Between the internal and external (large red arrows) aggregation, electron-transparent structures or intervals (large white arrowheads) appeared, then disappeared following the further assembly of dense particles in the aggregates (m1/2 and m4/2-m6/2; large red arrows and small black arrowheads). Small red arrows: dark threads, which formed by the congregation and linearization of the particles contained in these internal aggregations (m1/1 and m2/1). (**a4**) The aggregation of dense particles (fm7/1) incorporated the mitochondrion itself into N2, concurrently separating fm7 and causing part of fm7 to become electron transparent structure, which was further divided (fm7/2 and fm7/3) by internal assembly (black arrowheads). Mitochondrial congregations of particles formed a vesicle within N2 (white arrowhead) and appeared at the nuclear edge (fm5; large and small red arrows; black and white arrowheads). fm3 assembled dense particles to adopt a nuclear appearance, and their assembly diluted and dispersed fm4. (**a5** and **a6**) Within N2, incomplete mitochondria-to-nucleus transition was observed (black and white arrowheads). Within INC1, fm6 occurred mainly in the external assembly of the particles (small red arrowheads) to become electron-lucent, and the mitochondrial aggregation of dense particles was demonstrated at the edge of Nu1, which neighboured the dispersed fm7. Mitochondria assembled the particles to diffuse into Nu2 (fm8/1 and fm9/1), concomitantly rendering fm9/2 lucent; fm8/2 developed a nuclear appearance, and their aggregation condensed the organelles (fm10/1 and fm12-fm17/1), causing parts of mitochondria to become electron-transparent (fm10/2 and fm11/2) and transiently blackening the LMMs (red arrows). (**a6**) A high-magnification image of the inset in (**a5**). (**b**) Mitochondria-to-nucleus transition (opposite red arrowheads) recently occurred to accomplish partial fusion between individually constructed nuclei (N1 and N2); following the nuclear development of mitochondria, the area of the cytoplasm was consequently decreased and partitioned by the nuclei to form intranuclear inclusions (INC1-INC3) (**b1**). (**b2**) Mitochondrial aggregations of dense particles separated fm1 (electron-opaque fm1/1 and electron-lucent fm1/2), enlarged Nu2 (small red arrowheads), formed SDBs (black arrowheads), and created dark strands, while electron-transparent intervals (white arrowheads) was formed and a mitochondrion was diluted to look like a small INC (INC1/m1) at the nuclear edge. Incomplete nuclear conversion was observed between the nuclei (white arrow and black arrowhead), and two strands (red arrows), which were derived from mitochondrial assembly of the particles, combined to form a nuclear tubule (white arrow). (**b3**) In the low-magnification image (**b1**), mitochondrial congregation of dense particles (black and white arrows; black arrowheads) caused a small portion of N2 to look like an INC (INC2), in which the mitochondria almost completed nuclear transition. Nu2 appeared to be formed by the mitochondrial assembly of particles. (**b4** and **b5**) Both SDBs (black arrowheads) and SDIBs (white arrowheads) turned into dense particles, whose further assembly for nuclear development was observed within N2 and at its edge (small red arrowheads). Newly formed nuclear pieces (opposite red arrowheads) caused the partitioned cytoplasm to become a closed INC (INC3), in which condensed mitochondria (m2/1 and m3/1) were dispersed into the nucleus (double red arrowheads) as well as the surroundings. With the dispersion of dense aggregates (m2/1 and m2/2; red arrows, black and small red arrowheads) into the electron-lucent or electron-dilute area of the INC for nuclear transition, INC2 ultimately disappeared. (**b6**) The aggregation of dense particles (fm2/1 and small red arrowheads) separated fm2 (electron-dense fm2/1 and -lucent fm2/2), simultaneously causing transparent fm2/2 to fuse with N2 via their external assembly to the periphery (small red arrowheads). (**c**) A nucleus (N) was constructed by fragmented mitochondria, which surrounded and assembled dense particles into the nucleus. Mitochondria (fm1 and fm2) assembled the particles both internally (fm1/1 and fm2/1) and externally (red arrows and small red arrowheads) to disperse into the nucleus (small red arrowheads), causing the organelles to become electron transparent (fm1/2, fm2/2 and fm2/3). At the edges of fm1/1 and fm2/1, the further assembly of dense particles formed dark threads (red arrows). At the nuclear edge and within the nucleus, the mitochondrial aggregation of particles for nuclear development was observed (opposite red arrowheads) (**c1**-**c4**). (**c5** and **c6**) Mitochondria assembled dense particles into the nucleus as well as the surroundings to form nuclear bubbles (b1 and b2), and their aggregation dispersed electron-opaque fm3/1 into the nucleus (double red arrowheads) and bubbles, leading to nuclear growth. In the cytoplasm, electron-opaque fm4/1 directly dispersed into dense particles, and fm4/2 lucent because of the external aggregation of the particles. LD: a lipid droplet, which could derive from one or more mitochondria (fm5/2 and white arrowheads) and turned into particles (small red arrowheads). (**d**) A small or young nucleus (YN) appeared in this cell, and the mitochondria assembled dense particles (such as fm1/1-fm3/1) into the nucleus and concurrently became electron-transparent (such as fm1/2-fm3/2). The aggregation of dense particles separated fm4 into lucent fm4/2 and dense fm4/1, which looked like a lipid droplet and fused with fm5/1. At the nuclear edge, the mitochondria assembled the particles to diffuse into the YN (opposite red, black and white arrowhead), and condensed mitochondria fragmented into dense particles in the cytoplasm (such as fm6-fm8) (**d1**-**d6**).

**Fig. 3. Initiation of nuclear formation began with mitochondrial fragmentation into dense particles.** TEM was performed on four K562 cells at the 2 (**a**), 8 (**b**) and 12 (**c** and **d**) h time points, and the micrographs showed that mitochondria fragmented into dense particles to initiate the formation of a nascent nucleus (NN). (**a**) Three nascent nuclei (NN1-NN3) appeared in this cell, and NN2 was formed by the mitochondrial assembly of dense particles (opposite red arrowheads), whose aggregation fused NN2 with NN1 (**a1-a3**). (**a4** and **a5**) Mitochondria completely fragmented into dense particles (small red arrowheads) to enlarge NN2, concurrently creating electron-transparent spots in the nascent nucleus (white arrowheads). Partially fragmented fm1 was dispersed into dense particles (small red arrowheads), which entered both NN1 and NN2, and the external aggregation to the periphery produced an electron-lucent body in the mitochondrion (white arrowheads). (**a6-a8**) NN3 looked like a fragmented mitochondrion, which was surrounded and enlarged by partially (fm2-fm4) and heavily or completely (black and opposite red arrowheads) fragmented mitochondria via dispersion into it in the form of dense particles. The aggregation of the particles in mitochondria formed electron-transparent spots (small white arrowheads), which appeared in the cytoplasm following dispersion of the organelles (large white arrowheads) and ultimately became particles. Heavily fragmented fm4 was dispersing into dense particles, which appeared within the mitochondrion and at its edge (small red arrowheads). (**a9** and **a10**) Condensed mitochondria (fm5 and fm6) further assembled to diffuse into NN1 as well as the surroundings in the form of dense particles (small red arrowheads), and a large lucent fm7/2 formed following the aggregation of the particles in the mitochondrion (fm7/1 and red arrows). (**b**) In addition to a YN, a nascent nucleus was built by the mitochondrial assembly of dense particles, and partially (such as fm1-fm3) or completely (white arrows, and black, white and small red arrowheads) fragmented mitochondria surrounded the YN to promote nuclear growth. The internal aggregation of dense particles (red arrows) separated mitochondria (fm2/1 and fm2/2; fm3/1 and fm3/2). At the nuclear edges, virus-like granules (VLGs) (white arrows) were transiently derived from mitochondrial fragmentations. Assembling the particles caused fm4 (fm4/1 and fm4/2) to become incorporated into the NN and become part of the nucleus, in which incomplete mitochondrion-to-nucleus transition was observed (black and white arrowheads). The aggregation of dense particles to the periphery (small red arrowheads), that is, external assembly, separated fm5 (electron-opaque fm5/1 and electron-lucent fm5/2) and concomitantly fragmented or dispersed the organelle, and the external assembly formed a large, electron-transparent DI1, causing the particles to enter the NN (small red arrowheads); moreover, both SDBs (black arrowheads) and SDIBs (white arrowheads) ultimately diffused into the particles, leading to nuclear development (**b1-b4**). (**c**) Mitochondria (or mitochondrion)-to-nucleus transition and thereafter nuclear fusion continuously and repeatedly occurred to build two large nuclei in the cell; mitochondrial aggregations of dense particles (black arrowheads) for nuclear formation created electron-lucent structures at the nuclear edges (white arrowheads) and caused mitochondria to disperse (such as fm1-fm4). Between the internal (MN1) and external (small red arrowheads) congregation, a lunar halo structure appeared, in which dispersion or aggregation of the particles was observed (white arrowheads) (**c1-c3**). (**c4** and **c5**) Both MN2 and fm5 formed mitochondria assembling dense particles for nuclear development, and MN2 tended to adopt the appearance of MN1 as the nuclear transition proceeded. Near MN2 and fm5, the mitochondrial aggregation of dense particles was observed (fm6/1 and fm6/2-fm6/4; fm7/1 and fm7/2), and their internal congregations (black arrowheads) further divided the electron-lucent part of fm6 (fm6/2-fm6/4). Through external assembly (small red arrowheads), fm7/2 became electron transparent. (**c5**) A high-magnification image of the inset in (**c4**). (**d**) Three nuclei of similar size (N1-N3) separately and simultaneously formed in the cell, and adjacent to N1, a group of condensed mitochondria appeared (opposite red arrowheads) (**d1** and **d2**). (**d3**) Mitochondrial aggregations of dense particles formed a nascent nucleus (NN1) and led to the formation of electron-transparent structures (DI1 and white arrowheads). A mitochondrion assembled the particles into NN1 (fm1/1) to become electron transparent, and the internal aggregation of dense particles (red arrow) further divided the electron-lucent region (fm1/2 and fm1/3). (**d4**) Near N1, mitochondria simultaneously underwent external congregation of particles to the peripheries to form a lunar halo structure (white arrowheads). The aggregation of dense particles separated the organelles (fm2/1 and fm2/2; fm3/1, fm3/2, small black and white arrowheads), caused parts of mitochondria to become lucent (fm2/2, fm3/2 and white arrowhead), and led to the formation of lipid droplets (fm2/1 and fm3/1), which aggregated dense particles to N1 (small red arrowheads). (**d5**) The aggregation of dense particles (black arrowheads and fm4/1) and the formation of electron-transparent spots or structures occurred at the edge of N3 as well as in the cytoplasm. (**d6)** The congregation of the particles separated fm5 (into fm5/1 and fm5/2), condensed mitochondria (such as fm5/1 and fm6-fm9), and caused the formation of SDBs or small mitochondria-like bodies (SMLBs) (black arrowheads) and SDIBs (white arrowheads). SDBs, SDIBs and large mitochondria were turned into dense particles (small red arrowheads).

**Fig. 4. Knockdown of LC3 increased free micronuclei by hindering nuclear fusion and enhancing mitochondria-to-micronucleus transition.** K562 cells were transfected with either small interfering RNA (siRNA) against LC3 (siLC3) or control siRNA (siCtrl) for 48 h, and the cells were gathered and cultured in fresh medium for 2 h before TEM observation. (**a**) In this control cell, MN1 was fused with the large nuclei (N1 and N2) by partially (fm1 and fm2) or completely (such as fm3 and fm4; small red arrowheads) fragmented mitochondria, while both MN2 and MN3 were already attached to N1 (**a1** and **a2**). (**a3** and **a4**) Compared to the large nuclei in control cells, N1 and N2 in this LC3-depleted cell were widely separated, and four micronuclei (MN1-MN4) existed between the large nuclei, while MN5 was relatively far away from N1. fm1 dispersed into N1 and micronuclei, and both fm2 and fm3 changed to a similar density to the nuclei. (**b**) Along with the neighbouring counterparts that had achieved nuclear transition to enlarge both N1 and N2, a mitochondrion (fm1) became embedded between the nuclei, dispersed to fuse with them (double red arrowheads), and developed a nuclear appearance. Less fragmented mitochondria (such as fm2-fm5, fm6/1 and fm7/1) diffused into nuclei (double red arrowheads) and/or into each other, while the mitochondrial assembly of dense particles transiently formed the ERL (white arrow), an SDB (black arrowhead) and an SDIB (white arrowhead), and all of them eventually became particles (**b1** and **b2**). (**b3** and **b4**) Compared to the control cell, mitochondrial fragmentation was enhanced, and nuclei (N1 and N2) were widely separated in this LC3-silenced cell. At the edge of N2, mitochondrial aggregation of dense particles (small red arrowheads) was observed (fm1, DI1, white arrows and black arrowhead). (**c**) Mitochondria-to-nucleus transition led to partial nuclear fusion between two large nuclei (N1 and N2; three abreast red arrows), and MN1 was attached to N2 by the mitochondrial assembly of dense particles (opposite red arrowheads), whose aggregation (small red arrowheads) formed a nascent nucleus (NN1). Between N2 and NN1, mitochondria assembled the particles for nuclear development (opposite white arrowheads). The aggregation of dense particles separated mitochondria (such as fm1/1-fm5/1 and fm1/2-fm4/2) and dispersed fm4 (dense fm4/1 and lucent fm4/2) into N2 (double red arrowheads) (**c1** and **c2**). (**c3** and **c4**) In this LC3-depleted cell, the mitochondrial aggregation of dense particles attached MN1 to N1 and joined together two large nuclei (N1 and N2) (opposite red arrowheads), while MN2 was widely separated from the large nuclei. Not far from MN2, two nascent nuclei were constructed by the mitochondrial assembly of dense particles, whose aggregation formed dark strands (large red arrows) and threads (small red arrows), both of which combined to look like the tubule-nuclear envelope (NE) of MN2. The aggregation of the particles separated (fm1/1 and fm1/2; fm2/1 and fm2/2) and condensed mitochondria (fm1/1, fm2/1 and fm3-fm6), causing parts of the organelles to become electron-lucent (fm1/2 and fm2/2). Based on mitochondrial fragments (such as fm7), SDBs (black arrowheads) and SDIBs (white arrowheads) were derived. (**d**) In this control cell, seven nascent nuclei (NN1-NN7) appeared, which were connected and surrounded by partially or completely fragmented mitochondria (opposite red arrowheads), and NN7 looked like a mitochondrion and linked two nascent nuclei (NN5 and NN6) (**d1-d4**). (**d5-d8**) In the LC3-deprived cell, four nuclei (YN1, YN2, NN1 and NN2) were separately formed and/or enlarged (**d5**). (**d6**) YN2 was enlarged by the mitochondrial assembly dense particles (black arrowheads), and their aggregation led to the formation of lucent or dilute structures at the nuclear edge (white arrowheads). Less fragmented mitochondria (fm1, fm2, fm3/1, fm3/2 and DE1) were becoming particles. (**d7**) NN1 was at the initial stage of nuclear formation and was enlarged by the mitochondrial assembly of dense particles (opposite red arrowheads). (**d8**) The mitochondria assembled dense particles to fragment concurrently, forming a group whose formation promoted the nuclear development (NN2) of enclosed organelles. The external assembly of dense particles caused part of a mitochondrion to become electron transparent, and the internal aggregation (small black and white arrowheads) further divided the lucent part (fm1/1, fm1/2 and large white arrowhead). Within NN2, traces of mitochondrial fission were displayed (fm2/1, black and white arrowheads), while most mitochondria had completely fragmented into dense particles.

**Extended data Figs**

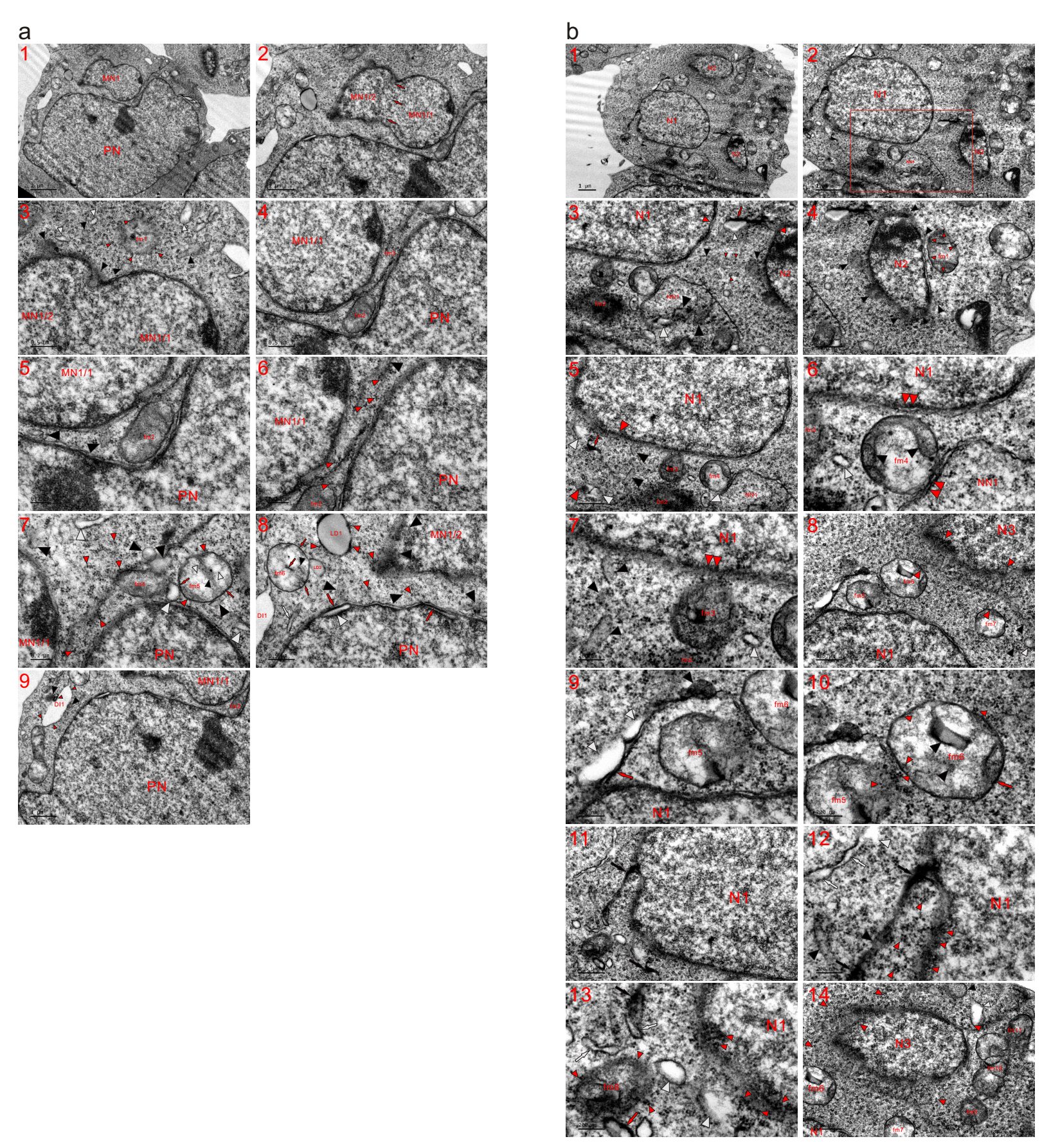

**Extended data Fig. 1. Partially and completely fragmented mitochondria existed between nuclei.** Transmission electron microscopy (TEM) was performed on two K562 cells at the 2 h time point, and the micrographs revealed that more than one nucleus separately formed in a single cell. (**a**) In addition to a PN in the cell, one micronucleus (MN1) was observed and appeared to have been formed by the combination of two smaller micronuclei (MN1/1 and MN1/2) via nuclear fusion (three abreast red arrows). At the edge of MN1, fm1 was dispersing into dense particles (small red arrowheads). During mitochondrial fragmentation, small dense bodies (SDBs) (black arrowheads) or small dilute bodies (SDIBs) (white arrowheads) were intermediately formed, and both of them eventually became particles (**a1-a3**). (**a4-a7**) Between PN and MN1, partially (fm2) or completely (such as fm3 and black arrowheads) fragmented mitochondria diffused into dense particles (small red arrowheads) for nuclear development, leading to the combination of nuclei (PN and MN1) or to nuclear merging. At nuclear edges, most mitochondria had completely fragmented and dispersed into dense particles (small red arrowheads), and only less fragmented ones (fm4 and fm5) and vestigial mitochondrial fragmentations (black and white arrowheads) were observed. fm4, SDBs and SDIBs diffused into the particles, and both the internal aggregation of dense particles (small black arrowhead) and their external assembly to the periphery (red arrows and arrowheads) occurred in and around fm5, concurrently creating electron-transparent spots or areas in the organelle (small white arrowheads). (**a8** and **a9**) Within nuclei and at nuclear edges, mitochondria assembled dense particles (small red arrowheads) for nuclear development (red arrows, black and white arrowheads) with complete fragmentation. The mitochondrial aggregation of the particles led to the formation of dark strands (red arrows) and an electron-lucent structure (white arrowhead) in the PN. Their external assembly temporarily blackened the limiting membrane (LMM) of fm6 (small red arrows), in which aggregations of dense particles also occurred (red arrow and black arrowhead). Based on mitochondrial fragmentation, an endoplasmic reticulum-like (ERL) structure (white arrow) was derived. Mitochondrial assembly of the particles caused the formation of lipid droplets (LD1 and LD2), which consequently became dense particles (small red arrowheads). Adjacent to fm6, their external aggregation (black and small red arrowheads) formed a large electron-transparent body (DI1); fm: fragmented mitochondrion. (**b**) In addition to large N1, two small nuclei (N2 and N3) appeared in the cell. Neighbouring N1, a nascent nucleus (NN1) was formed by the mitochondrial assembly of dense particles (large black and white arrowheads). Between N1 and N2, the organelles were heavily (red arrow, black and white arrowheads) or completely (small red arrowheads) fragmented. Near N2, mitochondria assembled dense particles to fragment for the nuclear growth (fm1 and black arrowheads), and the less fragmented fm1 was dispersing into N2 as well as the surroundings in the form of dense particles (small red arrowheads) (**b1-b4**). (**b3**) A high-magnification image of the inset in (**b2**). (**b5-b7**) At the edge of N1, mitochondria assembled dense particles, leading to fragmentation (black, white and opposite red arrowheads), and fm2 dispersed into the particles, while less fragmented mitochondria (fm3 and fm4) diffused into the nuclei (double red arrowheads) and the surroundings. fm4 mainly occurred in the external assembly of the particles (black arrowheads), and both SDBs (black arrowheads) and SDIBs (white arrowheads) eventually became dense particles. (**b8-b13**) Between N1 and N3, partially (fm5-fm7) or heavily (red arrow, black and white arrowheads) fragmented mitochondria were present, and the completely fragmented mitochondria had lost their mitochondrial morphology (opposite red arrowheads). fm5 was dispersed into dense particles (small red arrowheads), and fm6 exhibited both internal (black arrowheads) and external (red arrow and small red arrowheads) aggregations of the particles. At the edge of N1 and before complete fragmentation into dense particles (small red arrowheads), the mitochondrial assembly of the particles transiently formed ink masses (black arrows), a dark strand (red arrow), ERLs (white arrows), SDBs (black arrowheads) and SDIBs (white arrowheads). The assembly of the particles led fm8 to lose its LMM and let it disperse into dense particles (small red arrowheads). (**b14**) Immediately beside N3, mitochondria completely fragmented into particles for growth of the nucleus, and the organelles that were located somewhat far away were usually less fragmented (such as fm6, fm7 and fm9-11; black and white arrowheads).

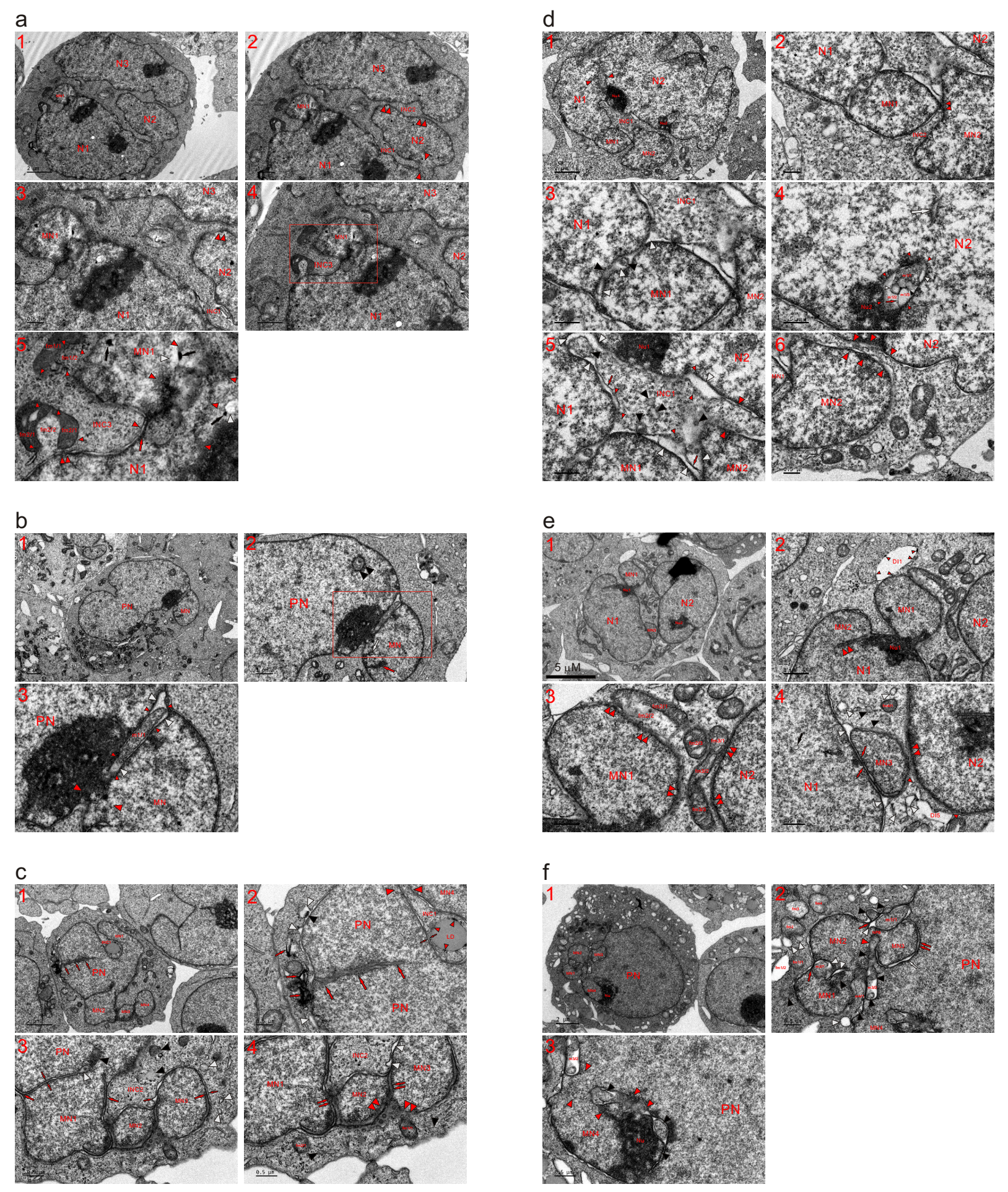

**Extended data Fig. 2. Attachment of an MN and partial nuclear fusion partitioned the cytoplasm to form an intranuclear inclusion (INC).** TEM was performed on six K562 cells at the 2 (**a** and **b**), 4 (**c**), 6 (**d** and **e**) and 12 (**f**) h time points, and the micrographs demonstrated that mitochondria fragmented to link individually constructed nuclei, concurrently partitioning the cytoplasm to form an INC. (**a**) Nuclei had been separately formed (N1-N3 and MN1), and partial nuclear fusion (opposite red arrowheads) between nuclei (N1 and N2) partitioned the cytoplasm to form an opened INC (INC1). A closed INC (INC2) was formed by partial merging between N2 and N3 (double red arrowheads) (**a1-a3**). (**a4** and **a5**) MN1 was attached to N1 by the mitochondrial assembly of dense particles for nuclear development (opposite red arrowheads), and its attachment compartmentalized its cytoplasm to form INC3 at the nuclear edge. The mitochondrial assembly of dense particles or traces of mitochondrial fragmentations were observed in the nuclei (black arrows, red arrow and white arrowheads). The aggregation of particles separated the organelles into dense (fm1/1 and fm2/1) and dilute (fm1/2 and fm2/2) parts, and dense particles in fm1/2 (small red arrowheads) reassembled for nuclear development to enlarge MN1; moreover condensed mitochondria (fm1/1 and fm2/1) were dispersed into N1 (double red arrowheads) and the surroundings in the form of dense particles (small red arrowheads). (**a5**) High-magnification images of the insets in (**a4**). (**b**) An MN was attached to the PN by the mitochondrial assembly of dense particles (m1/1 and opposite red arrowheads), whose internal aggregation formed m1/1; between their internal (m1/1) and external (small red arrowheads) congregations, electron-lucent or electron-dilute structures appeared, which could disappear to accomplish complete merging between PN and MN following further assembly of the particles in m1/1 as well as in the external aggregates. In either nucleus, incomplete mitochondrion-to-nucleus transition was observed (double black arrowheads and red and black arrows) (**b1**-**b3**). (**b3**) A high-magnification image of the inset in (**b2**). (**c**) Even in the PN, vestiges of nuclear fusion or the mitochondrial assembly of dense particles were observed (three abreast red arrows); traces of merging between MN1 and PN were discerned, and the attachment of the micronucleus partitioned the cytoplasm to form INC1, which was sealing by the nuclear development of mitochondria-derived particles (opposite red arrowheads). Within INC1, a lipid droplet (LD) diffused into dense particles (small red arrowheads), whose aggregation transiently formed tubule-NE fragments (opposite red arrows), falsely suggesting that the formation of the INC was due to herniation or invagination of the cytoplasm. At the edge of the PN, mitochondrial aggregations of dense particles for fragmentation are displayed (red arrows, black and white arrowheads) (**c1** and **c2**). (**c3** and **c4**) Nuclear fusions occurred among micronuclei (MN1-MN3) as well as between MN1 and PN by the mitochondrial aggregation of dense particles (double red arrows), whose assembly temporarily formed nuclear tubules (opposite red arrow), dark strands (large red arrows) or threads (small red arrows), SDBs (black arrowheads) and SDIBs (white arrowheads). Linkages of micronuclei (MN1-MN3) compartmentalized the cytoplasm to form INC2, and at the edges of the micronuclei, mitochondria assembled particles for addition into them (such as fm1/1, fm2/1 and double red arrowheads). (**d**) Incomplete mitochondria-to-nucleus transition (opposite red arrowheads) demonstrated between large nuclei (N1 and N2); partial fusion and attachments of micronuclei (MN1 and MN2) partitioned cytoplasm to shape INC1. MN1 was fused with N1 by the mitochondrial assembly of dense particles (black and white arrowheads), and fractional merging between micronuclei created INC2, in which fragmented mitochondria or a mitochondrion almost completely achieved nuclear transition (**d1-d3**). (**d4**) Within N2, a mitochondrion (m1) occurred both internally (m1/1, small black and white arrowheads) and externally (small red arrowheads) to assemble dense particles, whose aggregation further divided its electron-lucent part (m1/2 and m1/3). Not far from m1, an ERL (white arrowheads) appeared, representing incomplete mitochondrion-to-nucleus transition and mitochondrial congregation of the particles, which assembled and compacted to form Nu2. (**d5** and **d6**) Within INC2, the mitochondrial assembly of dense particles (small red arrowheads) for nuclear development was demonstrated (red arrows, and black and white arrowheads); after the mitochondria-to-nucleus transition expanded both N2 and MN2 (opposite red arrowheads), INC1 was consequently sealed to become a closed one. (**e**) The assembly and compaction of mitochondria-derived dense particles formed a nucleolus (Nu1) to join MN1 with N1, and an unfinished mitochondria-to-nucleus transition is shown between N1 and MN2 (double red arrowheads). At the edge of MN1, the external assembly of dense particles (small red arrowheads) caused a mitochondrion to become electron-lucent (DI1) (**e1** and **e2**). (**e3**) The organelles assembled dense particles into nuclei (double red arrowheads), and their aggregation led to the formation of condensed mitochondria (fm2/1 and fm2/3; fm3/1-fm3/3), which were dispersed into the particles to enlarge MN1 and N2. (**e4**) Within N1, there existed a vesicle (black arrow) representing incomplete the nuclear transition of a mitochondrion. An MN was attached to both N1 and N2 by the mitochondrial assembly of dense particles (red arrows and double red arrowheads), whose aggregations caused mitochondria to fragment concurrently and consequently to form SDBs (fm4/1 and black arrowheads) and SDIBs (white arrowheads). At the edge of N2, a mitochondrion assembled the particles into the nucleus to become electron transparent (DI5). (**f**) During their nuclear development, mitochondria (m1/1 and white arrowhead; m2/1 and white arrowhead; m3/2 and black arrowheads) included themselves in nuclei via the assembly of dense particles (red arrows), whose aggregation (black and white arrowheads) occurred in the neighbouring m2/1 within MN1. Adjacent to m1/1, mitochondrion-derived particles achieved nuclear transition (black arrowheads); m4 (m4/1, black and white arrowheads) assembled particles for nuclear development to combine MN1 and MN4. The aggregation of dense particles formed an NPB, which linked MN2 and MN3. Either the nuclear tubule (white arrow) or the gap between MN2 and MN3 disappeared through further assembly of the particles (double red arrows and red and black arrowheads). In the cytoplasm, a mitochondrion aggregated dense particles to form electron-dense fm1/1 and electron-lucent fm1/2, and the other mitochondria (fm2-fm4 and white arrowheads) were becoming particles (**f1** and **f2**). (**f3**) Between nuclei (MN4 and PN), partial fusion (opposite red arrowheads) was accomplished, and mitochondria assembled dense particles to complete nuclear merging between MN4 and PN by forming Nu and nuclear development (black and red arrowheads).

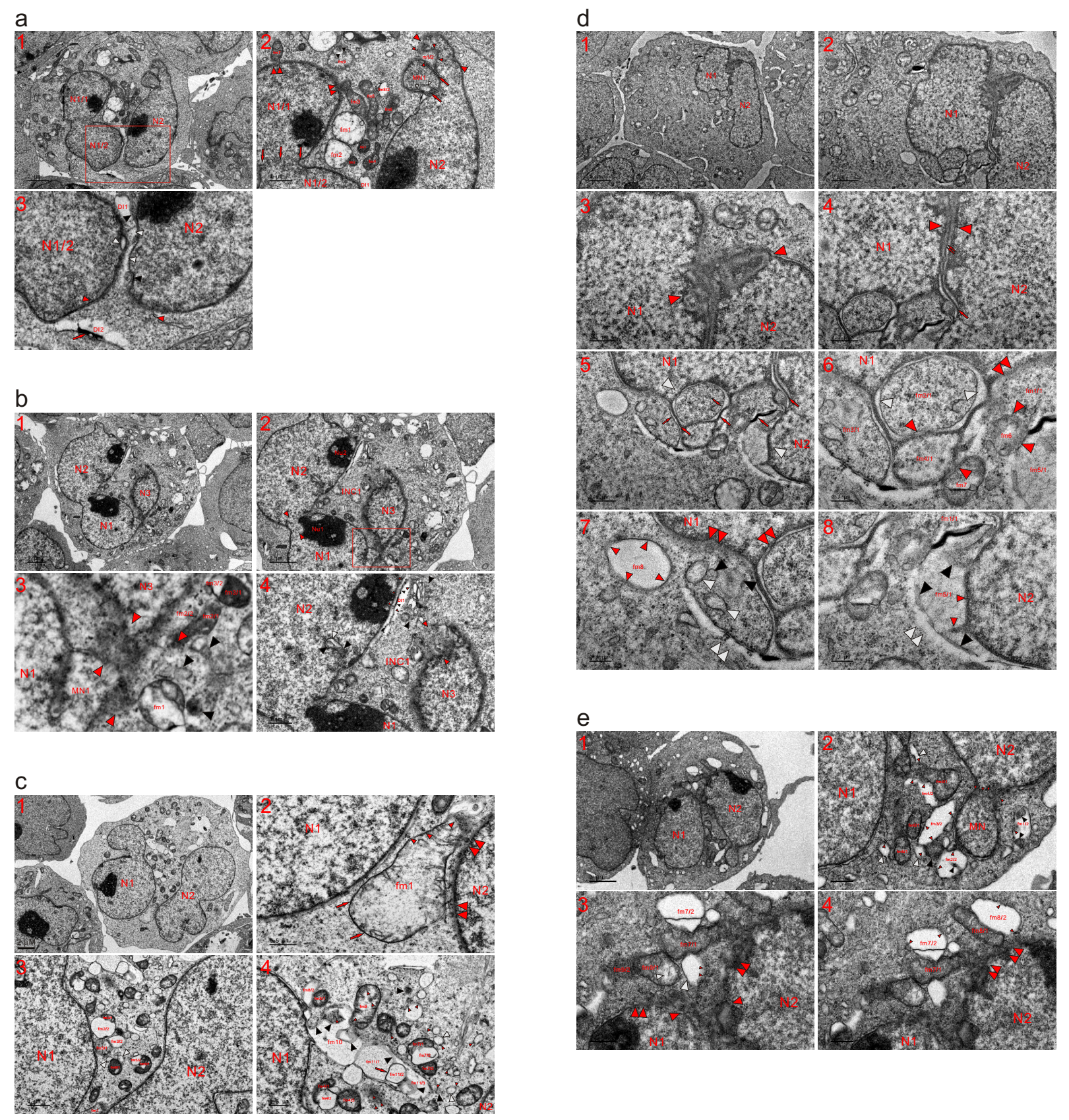

**Extended data Fig. 3. Mitochondria assemble dense particles for nuclear development to link large nuclei**. TEM was performed on five K562 cells at the 2 (**a** and **b**), 8 (**c** and **d**) and 12 (**e**) h time points, and micrographs showed that mitochondria fragmented into dense particles for nuclear transition to combine individually formed large nuclei. (**a**) The nuclear development of mitochondria to enlarge both nuclei (N1/1 and N1/2) consequently fused them (three abreast red arrows), resulting in a large N1. MN1 expanded and attached to N2 by the mitochondrial assembly of dense particles (m1, red arrows and white arrowheads). Adjacent to MN1, the organelle became dilute (m1/2) via the external assembly of the particles (small red arrowheads), whose reassembly enlarged N2 (opposite red arrowheads). Partial nuclear fusion between two large nuclei (N1 and N2) was accomplished by the mitochondrial assembly of dense particles (DI1, black and white arrowheads; DI2, red arrow and opposite red arrowheads). Within the partitioned cytoplasm, the externally assembled dense particles caused the organelles (fm1 and fm2) to become electron-transparent concurrently, forming dense bodies (DE1 and DE2); condensed mitochondria (such as fm3-fm7) dispersed into the nuclei (double red arrowheads) as well as into each other, and the electron-lucent fm8/2 was diffusing. Both the internal aggregates (small black arrowheads) and whole fm9 were becoming particles simultaneously, achieving complete fragmentation (**a1-a3**). (**a3**) A high-magnification image of the inset in (**a1**). (**b**) Partial nuclear fusion was achieved between the nuclei (N1 and N2) (opposite red arrowheads) and mitochondrial assembly of dense particles (fm1; fm2/1 and fm2/2; fm3/1 and fm3/2; black and opposite red arrowheads) for nuclear development to expand nuclei (N1, N3 and MN1), concurrently accomplishing fractional merging between N1 and N3, and their linkage combining N2 partitioned the cytoplasm to form INC1. At the opening of INC1, mitochondrial fragmentations (opposite red and black arrowheads) occurred for nuclear development, and several mitochondria jointly assembled the particles to shape a structure of Golgi complex-like (opposite black arrowheads). Internally (small black arrowheads) and externally (small red arrowheads) mitochondrial assembly of dense particles concomitantly created electron-transparent DI1 at the edge of N2, in which incomplete mitochondrion-to-nucleus transition (black and white arrowheads) occurred (**b1-b4**). (**b3**) A high-magnification image of the inset in (**b2**). (**c**) Between N1 and N2, a large mitochondrion (fm1) assembled dense particles (small red arrowheads) to fuse with N2 (double red arrowheads) and transiently blackened its LMM (red arrows) (**c1** and **c2**). (**c3** and **c4**) Between the nuclei, the aggregation of dense particles separated organelles, leading to the formation of condensed mitochondria (such as fm2/1-fm8/1, fm4/3 and fm7/3) and letting parts of the organelles become electron-lucent (fm2/2-fm8/2 and white arrowheads). fm9 mainly exhibited external assembly of the particles (small red arrowheads), and their aggregation in the organelles (red arrow and black arrowheads) formed large electron-transparent or electron-dilute bodies (fm10 and fm11/1-fm11/3), and either SDBs (black arrowheads) or SDIBs (white arrowheads) ultimately dispersed into dense particles (small red arrowheads). (**d**) Two large nuclei (N1 and N2) were fused by mitochondrial aggregations of dense particles (opposite red arrowheads), whose further assembly dispersed (double red arrows) a dark strand (red arrow) (**d1-d4**). (**d5-d8**) The external aggregation of dense particles to fringes from neighbouring mitochondria formed dark strands (red arrows), and their further assembly within the internal aggregates (such as fm2/1 and fm3/1) intermediately formed dark threads at the edges of these aggregates (small red arrows). Between the external and internal (fm2/1) aggregations of the particles, electron-lucent intervals (white arrowheads) were created. The aggregation of dense particles caused mitochondria to disperse into the nucleus (such as fm1/1-fm3/1 and double red arrowheads) as well as into each other (red arrowheads), and mitochondria mainly conducted the external assembly of the particles to be diluted (such as fm4/1; fm5/1, black and small red arrowheads). Within fm3/1, both SDBs (black arrowheads) and SDIBs (white arrowheads) were dispersing into dense particles; electron-lucent structures became elongated (double white arrowheads) via their connection, and electron-opaque mitochondria (fm6 and fm7) diffused via further aggregation of dense particles. Between N1 and N2, mitochondria (fm1/1-fm5/1 and fm6-fm8) jointly assembled the particles (opposite white arrowheads) to promote their own nuclear development, and fm8 became electron-dilute through the assembly of dense particles to its periphery (small red arrowheads). (**e**) Mitochondrial assembly of dense particles (small red arrowheads) expanded nuclear sizes (N2 and MN), concurrently leading the MN to become an attached micronucleus. The aggregation of the particles (black and red arrowheads) condensed mitochondria (fm4/1-fm6/1 and large black arrowheads) and created electron-transparent bodies (fm1/2-fm4/2 and large white arrowheads) (**e1** and **e2**). (**e3** and **e4**) N1 and N2 joined together by the mitochondrial assembly of dense particles into both nuclei (double red arrowheads). The aggregations of the particles separated mitochondria (dense fm7/1 and fm8/1; lucent fm7/2, fm8/2 and white arrowhead) and dispersed the condensed parts into N2 (double red arrowheads); through the assembly of dense particles to peripheries (small red arrowheads) (the external assembly), the electron-transparent parts (fm7/2, fm8/2 and white arrowhead) ultimately dispersed into the cytoplasm; their aggregation created lucent spots in electron-opaque fm9/1 and dispersed fm9/2 into dense particles.

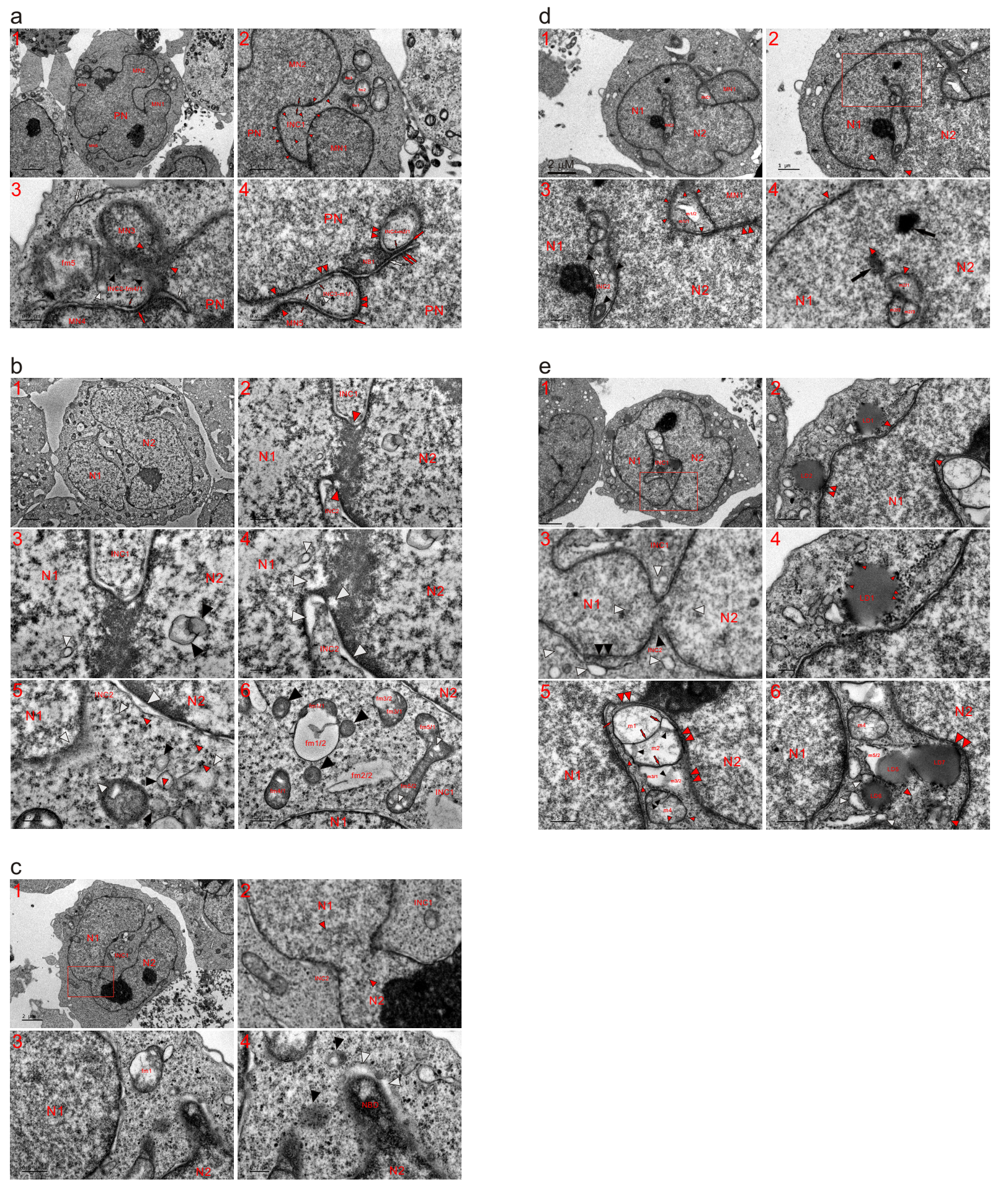

**Extended data Fig. 4. The formation of any INC was merely due to partitioning of the cytoplasm and incomplete mitochondria-to-nucleus transition**. TEM was performed on five K562 cells at the 4 (**a**), 8 (**b** and **c**) and 12 (**d** and **e**) h time points, and the attachment of an MN or partial nuclear fusion partitioned the cytoplasm to form an INC, which could be either small or large and was either opened or closed. (**a**) Attachments of MN1 and MN2 partitioned the cytoplasm to form INC1. Within INC1 and at its opening, mitochondria completely fragmented into dense particles (small red arrowheads) for nuclear development to enlarge the nuclei (PN, MN1 and MN2); the aggregation of the particles transiently formed the tubule-nuclear envelope (NE) fragment (opposite red arrows), which led to a false impression about forming the INC via the herniation or invagination of cytoplasm. Adjacent to the nuclei, mitochondria (such as fm1-fm3) assembled dense particles to fragment concurrently, providing the particles for nuclear expansions (MN1 and MN2), which consequently led to the sealing of INC1 (**a1** and **a2**). (**a3**) MN3, which looked like a condensed mitochondrion that being assembled dense particles, was merging with the PN by mitochondrial aggregation of the particles (opposite red arrowheads); their assembly (large and small red arrows) led the fragmented organelle to look like an INC (INC2-fm4/1) between MN3 and MN4. Within INC2, both SDBs (black arrowheads) and SDIB (white arrowhead) were dispersing into dense particles, whose assembly caused electron-opaque fm5 to disperse into micronuclei (MN3 and MN4) and INC2-fm4/1. White arrows: mitochondrial aggregations of dense particles transiently formed an ERL in the cytoplasm and one nuclear tubule in PN. (**a4**) Incomplete mitochondria-to-nucleus transition displayed in PN (NB1, INC3 and INC4-m2/1). The external assembly of dense particles from neighbouring mitochondria formed a thick strand (large red arrow), and two dark strands (red arrow and double red arrows) combined to shape the nuclear tubule (white arrow). Mitochondrial assembly of dense particles for nuclear development transiently caused the fragmented organelles to look like intranuclear inclusions (INC3 and INC4), and their further aggregation and linearization within the internal aggregates (m1/1 and m2/1) formed dark threads (small red arrows), which combined a dark strand (large red arrow) to shape the tubule-NE fragment (opposite small red arrows). Both dark strands and threads dispersed or disappeared (double red arrowheads) following aggregation of the particles for nuclear development as well as the sealing of INC3 (opposite red arrowhead). White arrowhead: an SDIB dispersed within INC3; NB1: a nuclear body, which derived from the mitochondrial aggregation of dense particles and was diffusing. (**b**) Mitochondrial aggregations of dense particles expanded nuclear sizes (N1 and N2), concomitantly combining two nuclei (opposite red arrowheads) and partial nuclear fusion compartmentalized cytoplasm to form INC1 and INC2. Incomplete mitochondria-to-nucleus transitions (black and small arrowheads) and traces of the mitochondrial assembly of dense particles (large white arrowheads) were observed in nuclei or at the nuclear edge, where a completely fragmented mitochondrion had lost its morphology (opposite white arrowheads). From those heavily fragmented, SDBs (black arrowheads) and SDIBs (small white arrowheads) were derived, both of which eventually turned into particles in the cytoplasm (small red arrowheads) (**b1-b5**). (**b6**) At the opening of INC1, the aggregation of dense particles ultimately caused mitochondria (fm1/1, fm2/1, fm4/1 and black arrowheads; fm1/1-fm3/2) to disperse into the particles and separated dense fm5 (fm5/1 and fm5/2), in which particle assembly created electron-lucent spots (small white arrowheads). (**c**) Mitochondrial aggregations of dense particles widened the linkage between individually built N1 and N2 (opposite red arrowheads), and the combination of nuclei consequently partitioned the cytoplasm to form INC1 and INC2 (**c1** and **c2**). (**c2**) A high-magnification image of the inset in (**c1**). (**c3** and **c4**) Aggregation of dense particles formed a nuclear bud (NBD), and their assembly led to the creation of electron-lucent structures at the edge of the NBD (white arrowheads), whose formation together with the nuclear development of fragmented mitochondria (such as fm1 and black arrowheads) promoted the sealing of INC1. (**d**) The mitochondrial aggregation of dense particles (small red arrowheads) to enlarge nuclei (N1 and MN2) while attaching MN1 to N1 (opposite white and double red arrowheads). Attachment of the micronucleus partitioned the cytoplasm to form closed INC1, in which aggregation of the particles formed electron-opaque m1/1 and electron-transparent m1/2. Partial nuclear fusions (N1 and N2) at two sites (opposite red arrowheads) compartmentalized the cytoplasm to form a closed INC (INC2), whose size had been reduced with nuclear expansion on four sides around it by the mitochondrial assembly of dense particles, whose aggregation was observed within INC2 and in the nuclei (black arrows, black and white arrowheads), representing incomplete mitochondria-to-nucleus transition. Immediately alongside INC2 and in the nucleus, the congregation of the particles separated the organelle (m2/1-m2/3), which became a nuclear mitochondrion due to its neighbouring counterparts having accomplished nuclear conversion (**d1-d4**). (**d4**) A high-magnification image of the inset in (**d2**). (**e**) Compared to the fusion (opposite red arrowheads) at the upper part of the nuclei (N1 and N2), the nuclear fusion at the lower site appeared to be recently completed (opposite white arrowheads) by the mitochondrial aggregation of dense particles (black and white arrowheads), and nuclear fusions at two sites partitioned the cytoplasm to form a closed INC1 and the small INC2. At the edge of N1, the aggregation of dense particles caused part of a mitochondrion assembling into the nucleus (double black arrowheads) and dispersed SDIBs. At the nuclear edge, lipid droplets (LD1 and LD2) further assembled into larger granules (small red arrowheads) and fused with N1 (double red arrowheads) to promote nuclear growth (**e1-e4**). (**e3**) A high-magnification image of the inset in (**e1**). (**e5** and **e6**) Within INC1, mitochondria (m1-m3) assembled dense particles (red arrows, small black and red arrowheads) to become electron transparent and to disperse into the nucleus (double red arrowheads), and the internal assembly of the particles (black arrowhead) divided m3 (m3/1 and m3/2). The congregation of the particles formed SDBs (black arrowheads) in m4, which diffused into dense particles (small red arrowheads) whose assembly separated m5 (m5/2 and LD5). These lipid droplets (LD5-LD7) could be derived from mitochondrial aggregation of the particles (such as m5/2 and white arrowheads), and LD7 further assembled dense particles to fuse with N2 (double red arrowheads). Adjacent to LD7, a completely fragmented mitochondrion dispersed into the particles (opposite red arrowheads), which in turn reassembled and rearranged for nuclear development.

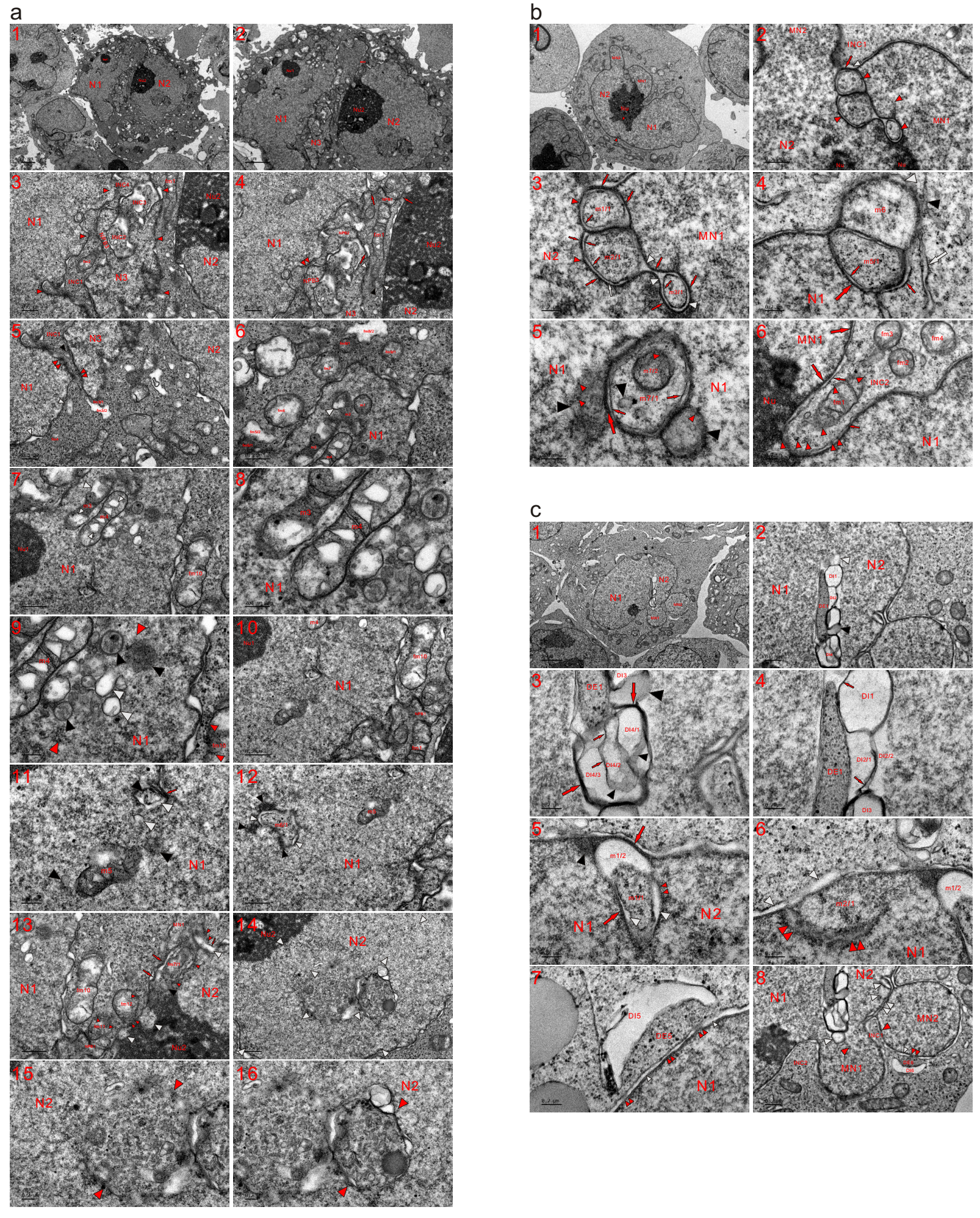

**Extended data Fig. 5. The formation of nuclear mitochondria occurred merely due to incomplete mitochondria-to-nucleus transition**. TEM was performed on three K562 cells at the 4 (**a** and **b**) and 8 (**c**) h time points, and micrographs revealed that the organelles became nuclear mitochondria along with their neighbouring counterparts having developed into the nucleus. (**a**) In addition to separately constructed N1 and N2, N3 was formed by the mitochondrial assembly of dense particles (opposite red arrowheads), whose aggregation first constructed nucleoplasmic bridges (NPBs) to link nuclei (NPB1-NPB3, red arrows), and NPB3 was fusing with N1 (double red arrowheads). Elongated fm1 dispersed into NPB1 and fragmented into the other organelle (black arrowhead) to link N3 and N2 (opposite white arrowheads). Nuclear fusion (between NPB3 and N1) partitioned the cytoplasm to create INC1, which was being sealed by the mitochondrial assembly of dense particles (black and double red arrowheads). Within INC1, less fragmented fm2 dispersed into dense particles, and fm3 assembled the particles into N3 (fm3/1) to become electron-lucent (fm3/2); dense fm4 diffused into N1 as well as the surroundings in the form of dense particles, whose external assembly caused the formation of nuclear bubbles (white arrowheads). Incomplete mitochondria-to-nucleus transition formed INCs (INC2-INC4) (**a1-a5**). (**a6-a9**) Within N1 and at its edge (such as fm5/1, fm5/2, fm6, fm7, fm8/1, fm8/2 and fm9/1), the mitochondrial aggregation of dense particles was observed, and several nuclear mitochondria appeared (m1-m4). Within the mitochondria (m3 and m4), the assembling particles created electron-transparent spots (small white arrowheads), which appeared in N1 following dispersion of the organelles (m2, m3 and large white arrowheads). Neighbouring m4, fragmented mitochondria formed a group (opposite red arrowheads), in which transiently built SDBs (black arrowheads) and SDIBs (white arrowheads) appeared, and some SDIBs looked like a vesicle. At the edge of N1, fm10 diffused into it and the surroundings in the form of dense particles (small red arrowheads). (**a10-a12**) A nuclear mitochondrion (m5) was dispersed in N1, and heavily or completely fragmented mitochondria had lost mitochondrial morphology (red arrow, black and white arrowheads). The appearance of m5 in the nucleus was merely due to its counterparts having undergone nuclear transition. Not far from m5, the aggregation of dense particles (black arrowheads) caused a mitochondrion (m6) to look like a closed INC, and between the internal (m6/1) and external (black arrowheads) congregation of m6, lucent intervals (white arrowheads) appeared. (**a13**) Aggregation of the particles for nuclear development (MN1, large and small arrows, and small red arrowheads) caused the incorporation of the organelles into N2 (m7/1, black and white arrowheads), and the mitochondrial assembly of dense particles formed dilute areas within Nu2 (white arrowheads). Both mitochondria (fm11 and fm12) dispersed into Nu2 and the surroundings in the form of dense particles (small red arrowheads). (**a14-a16**) Nuclear formation or four-sided growth (opposite white arrowheads) by the mitochondrial assembly of dense particles caused two groups of incompletely fragmented mitochondria to appear in N2 (opposite red arrowheads). (**b**) The mitochondrial assembly of dense particles to form a large Nu and to accomplish nuclear transition (opposite red arrowheads) concurrently merged individually constructed nuclei (N1 and N2). The attachment of micronuclei (MN1 and MN2) created small INCs (INC1 and INC2) at the nuclear edge, and the formation of Nu led to partial fusion between nuclei (N2 and MN1). Along with the nuclear development of their neighbouring counterparts to enlarge both N2 and MN1, the organelles undergoing nuclear transition became nuclear-localized mitochondria (red arrowheads), which underwent the internal (such as m1/1-m1/3) and external (large red arrows and small red arrowheads) aggregation of dense particles. Further assembly of the particles from the internal aggregates (m1/1-m1/3) to the peripheries formed dark threads (small red arrows), which combined dark strands to look like as a nuclear tubule (opposite small red and large white arrows). Between the internal and external aggregation of dense particles, electron-transparent lumens appeared (white arrowheads), and the external congregation of the particles (large red arrow and white arrowhead) caused m1 to incorporate itself into the nucleus while separating from INC1 (**b1-b3**). (**b4** and **b5**) During the nuclear transition of the organelles (m5 and m6), the mitochondria were incorporated into N1 with or without the help of other fragmented mitochondria (white and small red arrows; black and white arrowheads); m5 exhibited both internal (m5/1 and small red arrow) and external (large red arrow) aggregation of dense particles, and m6 displayed similar density to that of the nucleus and looked similar to a small MN. Within N1, incomplete mitochondria-to-nucleus transition was observed (m7/1, m7/2 and black arrowheads), and both m7/2 and SDBs (black arrowheads) were dispersing via further assembly of the particles (small red arrowheads), whose congregation or/and linearization intermediately formed dark strands (large red arrow) or threads (small red arrows). Nuclear transition of the neighbouring counterparts caused less fragmented mitochondria (m7 and black arrowheads) to become nuclear-localized. (**b6**) Mitochondrial assembly of dense particles to enlarge the nuclei (MN1 and N2) and to form Nu led to the attachment of MN1 to N2, consequently partitioning the cytoplasm with N1 to create INC2, in which completely fragmented mitochondria became dense particles (small red arrowheads), which entered Nu; the less fragmented mitochondria (fm1-fm4) were dispersing into the particles, whose further assembly diffused (small red arrowheads) into both dark strands (large red arrows) and threads (small red arrows) for nuclear development. (**c**) Nuclear fusion (N1, N2, MN1 and MN2) was conducted by fragmented mitochondria to form a large nucleus; between N1 and N2, the mitochondrial aggregation of dense particles was observed (DE1, DI1-DI3, black and white arrowheads). The assembly of particles formed dense bodies (large DE1 and large black arrowheads) and caused mitochondria to become electron-lucent or electron-dilute (DI1-DI3 and white arrowhead); the internal congregation of dense particles (small red arrows and small black arrowheads) further divided electron-transparent parts of the organelles (DI2/1 and DI2/2; DI4/1-DI4/3). Large red arrows: dark strands, which formed by the compacting of mitochondria-derived dense particles (**c1-c4**). (**c5** and **c6**) Between nuclei (N1 and N2) and at the nuclear edge, the mitochondrial assembly of dense particles was observed (electron-opaque m1/1 and -lucent m1/2), and their external assembly in m1 (large red arrow and black arrowhead) incorporated the organelle itself as a nuclear-localized mitochondrion or a nuclear mitochondrion; electron-lucent intervals (white arrowheads) formed between the internal (m1/1) and external (red arrow and small red arrowheads) congregations of the particles. Adjacent to m1, a large part of a mitochondrion (m2/1) almost completed nuclear transition accompanied by the three-sided assembly of dense particles (double red and white arrowheads). (**c7** and **c8**) At the edges of both N1 and MN2, the mitochondrial assembly of dense particles caused large electron-opaque bodies (DE5 and DE6) to fuse with nuclei (double red arrowheads) and allow electron-transparent lumens (small white arrowheads) to disappear, concurrently leading to the creation of large electron-lucent structures (DI5 and DI6). Between micronuclei (MN1 and MN2), mitochondria-derived dense particles assembled to close the gap (INC1) between MN1 and MN2 (opposite red arrowheads). Through enlarging nuclei (N1, N2, MN1 and MN2) by mitochondria-to-nucleus transition, both MN1 and MN2 became attached micronuclei, between which mitochondrial assembly of the particles was observed (single and double large white arrowheads), indicating the incomplete nuclear transition of the organelles.

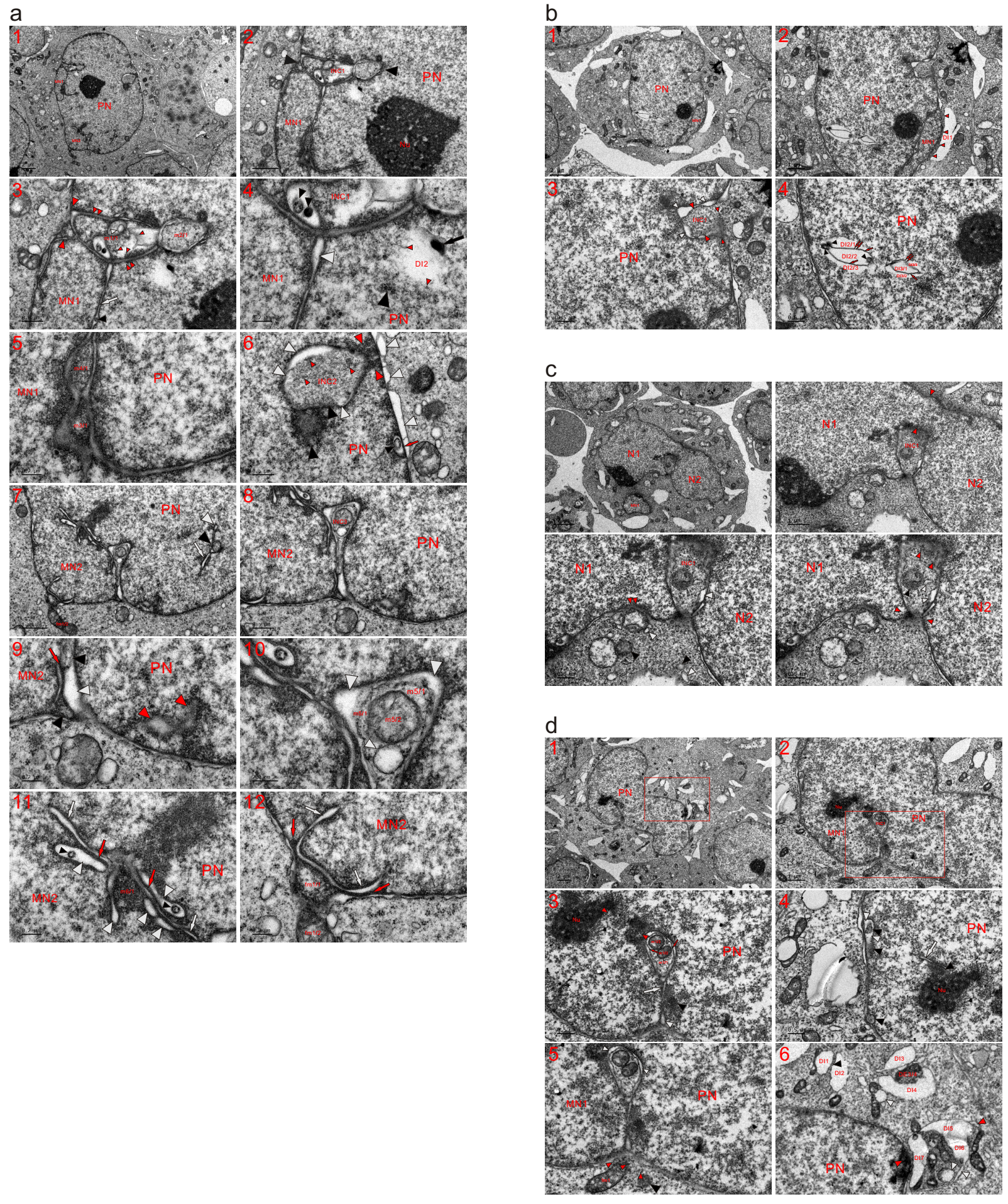

**Extended data Fig. 6. The mitochondria-to-nucleus transition enlarged the nucleus and sealed the INC.** TEM was performed on four K562 cells at the 4 (**a**) and 6 (**b-d**) h time points, and the micrographs demonstrated that incomplete nuclear conversion of the organelles created an INC, which was sealed following mitochondria-to-nucleus transition to enlarge nuclear sizes. (**a**) Incomplete mitochondria-to-nucleus transition (m1/1 and m2/1; black, white and small red arrowheads; double red arrowheads) transiently looked like an INC (INC1 and opposite black arrowheads), which was sealed by the mitochondrial aggregation of dense particles (opposite large red arrowheads). Adjacent to the INC and Nu, the aggregation of particles (black arrow and arrowhead and small red arrowheads) dispersed the electron transparency of the organelle (DI1). The mitochondria-to-nucleus transition enlarged both MN1 and PN, leading MN1 to merge with PN. Between the nuclei (MN1 and PN), incomplete nuclear conversion of the organelles was observed (m3/1, m4/1, white arrow, black and white arrowheads) (**a1-a5**). (**a6**) Mitochondrial aggregation of dense particles from both inside and outside the PN sealed INC2 (opposite red arrowheads), in which traces of mitochondrial fragmentation or the mitochondrial assembly of the particles were observed (black and white arrowheads), although the organelle had almost completely fragmented into dense particles (small red arrowheads). Immediately next to INC2, an SDB was dispersed (black arrowhead). At the nuclear edge and within the PN, the internal (red arrow and black arrowhead) and external (white arrowheads) aggregation of dense particles in mitochondria was observed, and the external assembly of dense particles to the periphery caused parts of the mitochondria to become electron-lucent, looking like nuclear bubbles (white arrowheads). (**a7-a11**) Similar to MN1, mitochondria assembled dense particles to expand both PN and MN2, causing MN2 to fuse with PN. Incomplete nuclear transition of the organelle created INC3, which was closed by the mitochondrial congregation of dense particles (red arrow, black and white arrowheads). Within INC3, the internal assembly of dense particles in a mitochondrion formed dense m5/1, m5/2 and a vesicle (small white arrowhead); between the internal and external congregation of the particles, electron-transparent structures or intervals (large white arrowheads) appeared. Between MN2 and PN, the mitochondrial aggregation of dense particles transiently formed nuclear tubules (white arrowheads), dark strands (red arrows), electron-lucent bodies or structures (white arrowheads), SDBs (small black arrowheads) and Nu-like m6/1, and through further assembly, the dark strands and SDBs dispersed into these electron-transparent structures or lumens to complete nuclear merging. Not far from INC3 and in the PN, the mitochondrial assembly of dense particles was observed (white arrow, and black, white and red arrowheads). (**a12**) At the edge of MN2, the internal assembly of dense particles caused part of a mitochondrion to adopt a nuclear appearance (fm1/1) and separated the organelle (fm1/1 and fm1/2). Within MN2, mitochondrial assembly of the particles intermediately formed dark strands (red arrows) and nuclear tubules (white arrows), both of which were derived from a mitochondrion or from neighbouring mitochondria. (**b**) MN1 fused with PN, and one or more mitochondria assembled dense particles to MN1 (red arrowheads) as well as the surroundings to form a large electron-transparent body (DI1). Mitochondrial aggregation of the particles (opposite large red arrowheads) for nuclear development transiently appeared as a small and shallow INC (INC1) at the nuclear edge. Between the internal and external congregation, electron-lucent structures (white arrowheads) were formed. At the opening of INC1, the assembly of mitochondria-derived dense particles from both sides of the INC was observed (small red arrowheads). Within the PN, the assembly of dense particles caused mitochondria to form large electron-transparent structures (DI2 and DI3), which were further divided (DI2/1-DI2/3 and DI3/1-DI3/3) by the internal congregation of the particles (red arrows and black arrowheads), whose further assembly dispersed dark threads (red arrows) (**b1-b4**). (**c**) Mitochondria-to-nucleus transition expanded the nuclear sizes of both N1 and N2, concomitantly leading to partial nuclear fusion (opposite red arrowheads), which in turn partitioned the cytoplasm to form INC1. Within the INC and at the nuclear edge, mitochondria fragmented to disperse into nuclei in the form of dense particles (single and double red arrowheads). In both the cytoplasm and partitioned cytoplasm (INC1), both SDBs and SDIBs were becoming particles, whose assembly caused the closure of INC1 (opposite red arrowheads) (**c1-c4**). (**d**) Mitochondrial assembly of dense particles for nuclear development (Nu, opposite red arrowheads, white arrow, black and white arrowheads) enlarged both nuclei (PN and MN1), and incomplete mitochondria-to-nucleus transition created INC1, which was almost sealed by the aggregation of the particles (white arrow, black and white arrowheads). In the cytoplasm, mitochondria fragmented (fm1 and black arrowhead) to disperse into nuclei in the form of dense particles (small red arrowheads). Within INC1, the assembly of dense particles in the internal aggregate of a mitochondrion (m1/1) formed dense m1/2 and m1/3, and further assembly of the particles formed dark threads at the edge of m1/1 (small red arrows); between the internal and external congregation of dense particles, electron-lucent or electron-dilute intervals (white arrowheads) were displayed (**d1-d5**). (**d5**) A high-magnification image of the inset in (**d2**). (**d6**) A high-magnification image of the inset in (**d1**), revealing that mitochondrial aggregations of dense particles led to the formation of electron-transparent or electron-dilute bodies (DI1-DI7 and white arrowheads); dispersion of the aggregate (black arrowhead) between DI1 and DI2 caused them to fuse and adopt the appearance of DI5 and DI6, while DI3 and DI5 seemed to connect into a circle to enclose condensed DE3/4. At the nuclear edge, the organelles assembled dense particles to form a group (opposite red arrowheads), whose formation accelerated the nuclear development of the enclosed mitochondria.

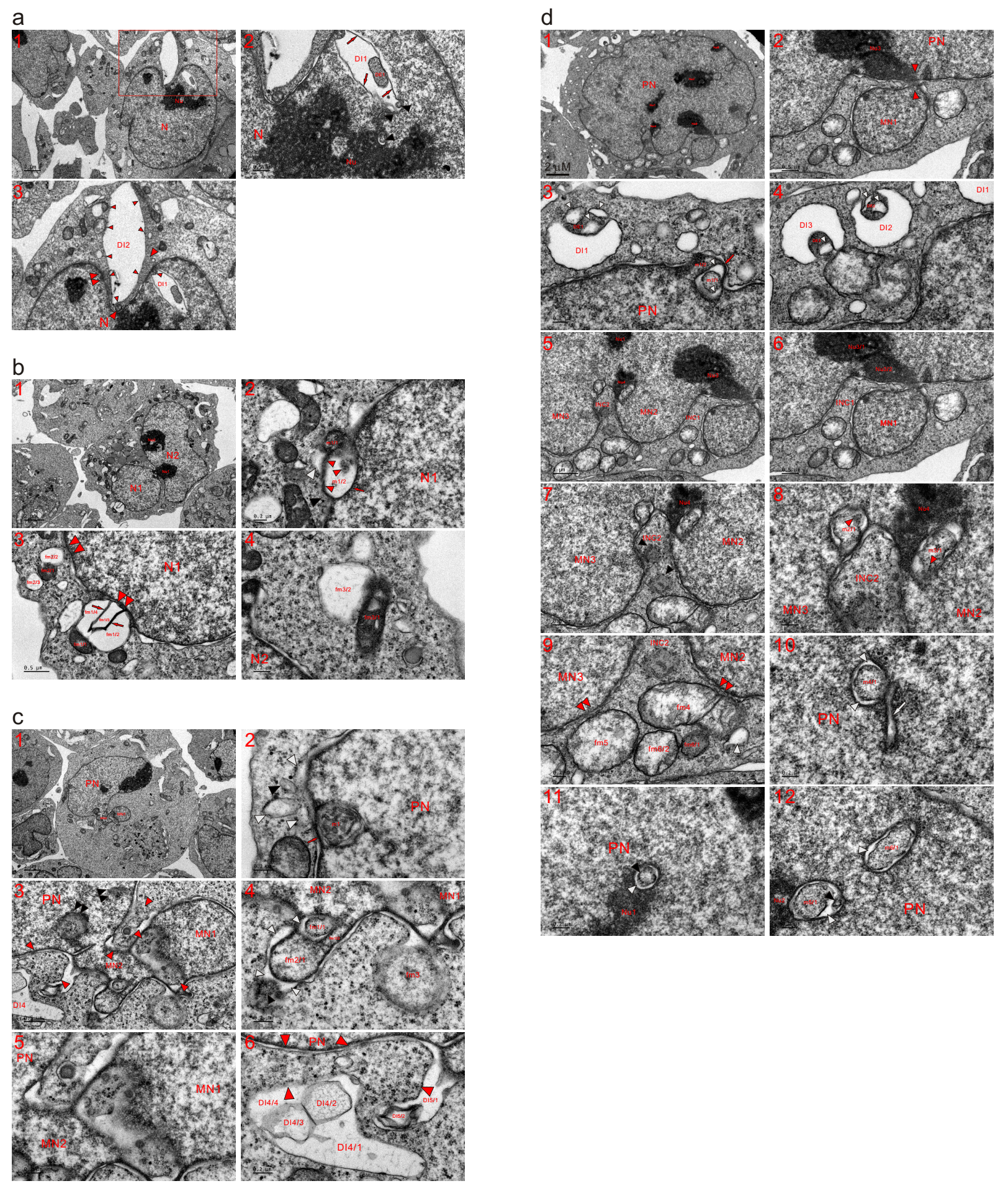

**Extended data Fig. 7. Internal and external aggregations of mitochondria occurred within nuclei and at the nuclear edge.** TEM was performed on four K562 cells at the 6 (**a**), 8 (**b** and **c**) and 12 (**d**) h time points, and the micrographs showed that mitochondria fragmented for nuclear development via the internal and/or external assembly of dense particles. (**a**) Within the nucleus and adjacent to Nu, mitochondrial aggregations of dense particles (DE1, red arrows and arrowhead) led to the formation of a large electron-transparent body (DI1). Through assembly of the particles at the periphery (small red arrowheads) for nuclear growth (double large and opposite red arrowheads), a larger lucent body was formed (DI2) (**a1-a3**). (**a3**) A high-magnification image of the inset in (**a1**). (**b**) Mitochondrial-derived dense particles assembled and compacted to form Nu1, which consequently caused nuclei (N1 and N2) to achieve partial fusion. Through assembly of the particles (m1/1, red arrow, and black and red arrowheads), a mitochondrion was included in N1, causing part of it to become electron-lucent (m1/2 and white arrowhead) (**b1** and **b2**). (**b3** and **b4**) The aggregation of dense particles separated mitochondria (fm1/1 and fm1/2-fm1/3; fm2/1-fm2/3) and caused the organelles to fuse with N1 (double red arrowheads) via the external assembly of dense particles; their internal congregations (red arrows) further divided the electron-transparent part of fm1 (fm1/2-fm1/3). In the cytoplasm and adjacent to N2, fm3 was separated into dense fm3/1 and lucent fm3/2 through the aggregation of dense particles. (**c**) In addition to PN, two micronuclei (MN1 and MN2) appeared in the cell. At the edge of PN, a mitochondrion (m1) assembled dense particles (red arrow) and was incorporated into the nucleus, and the internal aggregation of the particles caused m1 to look like a multivesicular body (MVB). At the nuclear edge, both an SDB (black arrowhead) and SDIBs (white arrowheads) were dispersed into the particles for nuclear development (**c1** and **c2**). (**c3-c5**) Between the nuclei and at the edge of PN, the mitochondrial assembly of dense particles promoted nuclear development (opposite red arrowheads); the organelles (fm1/1 and fm1/2; fm2/1) assembled the particles to become part of MN2, leading to the formation of lucent structures (white arrowheads), and an SDB (black arrowhead) diffused into the lucent structure. The aggregation of dense particles caused fm3 to lose its limiting membrane (LMM), adopt a nuclear appearance and fuse with MN1. Within PN, the mitochondrial assembly of dense particles or incomplete nuclear conversion of mitochondria was observed (double black arrowheads). (**c6**) The mitochondrial assembly of dense particles (opposite red arrowheads) into PN promoted its growth while forming large electron-transparent or electron-dilute bodies (DI4/1-DI4/4; DI5/1 and DI5/2), and DI4 (DI4/1-DI4/4) could be derived from more than one mitochondrion. The external assembly of dense particles at the mitochondrial periphery linked DI5 with DI4, and the internal aggregation of the particles formed DI4/2, DI4/3 and DI5/2. (**d**) Mitochondrial assembly of dense particles to expand both MN1 and PN narrowed the gap between nuclei and caused them to achieve partial fusion (opposite red arrowheads) (**d1** and **d2**). (**d3** and **d4**) At the nuclear edge, mitochondrial aggregations of the particles created electron-lucent bodies (DI1-DI3), and in the dense regions (DE1-DE3), the further assembly of dense particles was observed (small white arrowheads). A mitochondrion assembled particles to incorporate itself into PN (m1/1, m1/2, red arrow and white arrowheads), and through the further assembly of dense particles, m1/1 and a dark strand (red arrow) dispersed into lucent intervals, promoting nuclear development. (**d5-d9**) The attachment of MN1 partitioned the cytoplasm to form INC1; both micronuclei (MN2 and MN3) should have been individually constructed and merged with PN, similar to MN1. The attachments of MN2 and MN3 consequently compartmentalized the cytoplasm to create the opened INC2, in which mitochondria-derived SDBs (black arrowheads) were dispersing into dense particles. Within nuclei and neighbouring INC2, the internal aggregates of mitochondria (m2/1 and m3/1) were diffusing (red arrowheads). Immediately next to Nu3/1, mitochondrial assembly of the particles formed an ink body (Nu3/2), which underwent further assembly to enlarge Nu3/1. At the opening of INC2, mitochondria (fm4, fm5, fm6/1, fm6/2 and white arrowhead) assembled particles to disperse into the nuclei (double red arrowheads) as well as the surroundings, leading to the sealing or disappearance of the INC. (**d10-d12**) At several sites within the PN, incomplete mitochondrion-to-nucleus transition was observed (m4/1-m6/1, white arrow and black and white arrowheads), and mitochondria-derived dense particles assembled and compacted to become nucleoli (Nu1 and Nu2).

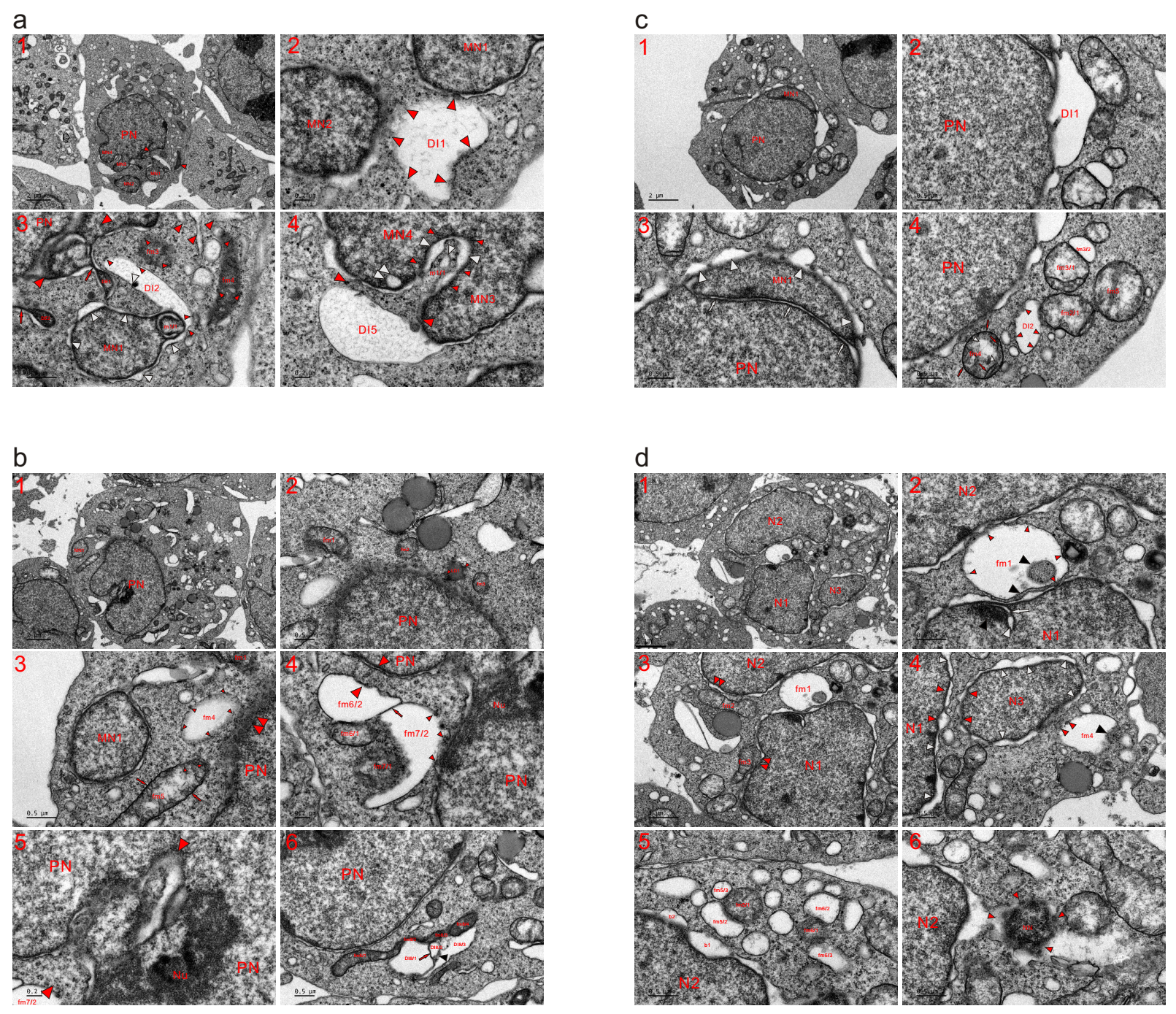

**Extended data Fig. 8. Mitochondria externally assemble dense particles to achieve nuclear transition and growth.** TEM was performed on four K562 cells at the 8 (**a**) and 12 (**b-d**) h time points, and the micrographs revealed that the external aggregation of dense particles to the periphery caused mitochondria to become electron-transparent. (**a**) In addition to a large PN, two free micronuclei (MN1 and MN2) and two attached ones (MN3 and MN4) were labelled. At the edges of the micronuclei (MN1 and MN2), mitochondrial assembly of the particles at the periphery (red arrowheads) caused a large mitochondrion to become electron-transparent (DI1) (**a1** and **a2**). (**a3**) At the edge of the PN, the mitochondrial aggregation of dense particles developed into a nuclear appearance (opposite red arrowheads) and formed dense bodies (DE1 and DE2) and dark strands (red arrows). A mitochondrion assembled particles to incorporate itself into MN1 (m1/1), around which nuclear bubbles or lucent structures formed (white arrowheads) during the mitochondrial aggregation of dense particles to enlarge the micronucleus. With the assembly of particles into MN1 (opposite white arrowheads) and the surroundings (small red arrowheads), a large electron-transparent body formed (DI2). Condensed mitochondria (fm3 and fm4) were being dispersed into dense particles (small red arrowheads) and formed a circle (opposite red arrowheads in **a1**) through connecting with nuclei (MN1 and PN) and fragmented mitochondria (DE1, red arrow and arrowheads), and the formation of the circle promoted the nuclear development of the enclosed mitochondria. (**a4**) Mitochondrial assembly of dense particles (opposite red arrowheads) enlarged both micronuclei (MN3 and MN4), produced a large lucent DI5, formed an INC (m1/1 and white arrowheads) and created a nuclear vesicle in MN4 (double white arrowheads). Between the internal (m1/1, small white arrowhead) and external aggregation of dense particles (small red arrowheads), electron-transparent intervals or structures were observed (large white arrowheads). (**b**) Adjacent to the PN, condensed mitochondria (such as fm1-fm3) developed a density similar to that of the nucleus, and a lipid droplet (LD1) derived from the mitochondrial aggregation of the particles dispersed into PN as well as the surroundings in the form of dense particles (small red arrowheads). fm4 became diluted and fused with the PN (double red arrowheads) via the external aggregation of particles to the periphery (small red arrowheads). Through further assembly, condensed fm5 blackened its LMM and was dispersing into dense particles (small red arrowheads), whose aggregation for nuclear development eventually joined the individually formed MN1 with PN (**b1**-**b3**). (**b4**) Mitochondria assembled dense particles into the nucleus (small red and opposite red arrowheads) and formed dense bodies or condensed mitochondria (fm6/1 and fm7/1), causing the organelles to become electron transparent (fm6/2 and fm7/2). These organelles were connected (red arrow), while the opaque parts (fm6/1 and fm7/1) dispersed into each other and re-entered lucent DIs (fm6/2 and fm7/2) via further particle assembly. DI: dilute body or phase. (**b5**) Near Nu, mitochondria fragmented into dense particles for nuclear development (opposite red arrowheads). (**b6**) At the edge of the PN, the organelles jointly and externally assembled dense particles to form elongated and condensed mitochondria (fm8/1-fm8/4), concurrently forming a large lucent body, which was further divided by internal aggregation (DI8/1-DI8/3). (**c**) Mitochondrial assembly of dense particles for nuclear growth formed a large electron-transparent body at the edge of the PN, enlarged both nuclei (PN and MN1) and combined them; their aggregation intermediately formed nuclear tubules (white arrows) and lucent structures or bubbles at the edge of MN1 (white arrowheads). In the cytoplasm, the external aggregation at the periphery (small red arrowheads) caused a mitochondrion to become electron transparent (DI2). Their assembly separated fm3 (fm3/1 and fm3/2), blackened the LMM (red arrows) of fm 4, formed tubular cristae (red arrows) and created electron-lucent spots (white arrowheads) in the organelle, while assembly of dense particles in mitochondria transiently caused the organelle (such as fm2/1, fm3/1, fm4 and fm5) to appear as occurrence of cristolysis. (**d**) Three nuclei were separately constructed, and the mitochondrial aggregation of dense particles for nuclear growth transiently formed lunar halo structures around them. Between N1 and N2, the internal (black arrowheads) and external (small red arrowheads) mitochondrial assembly of dense particles caused fm1 to become lucent and fuse with nuclei. Within N1, incomplete mitochondrion-to-nucleus transition or mitochondrial aggregation of the particles was observed (white arrow, black and white arrowheads). At the nuclear edges, both mitochondria (fm2 and fm3) developed a similar density to that of the nucleus and were being dispersed into nuclei (double red arrowheads) as well as the surroundings (**d1-d3**). (**d4**) Mitochondrial assembly of dense particles for nuclear expansion (both N1 and N3) led to the joining together of two nuclei (opposite red arrowheads). Around the nuclei, the external aggregation of dense particles formed electron-lucent structures, which appeared as lunar halos (white arrowheads). At the edge of N3, the mitochondrion (fm4) became electron-transparent by assembling dense particles into the nucleus (small red arrowheads) and the surroundings (black arrowheads). (**d5**) In the cytoplasm, mitochondrial aggregations of dense particles separated the organelles into dense (fm5/1 and fm6/1) and lucent (fm5/2 and fm5/3; fm6/2 and fm6/3) parts, which became vesicles (fm5/2, fm5/3 and fm6/2) and/or nuclear bubbles (b1 and b2). (**d6**) Adjacent to N2, mitochondria-derived dense particles (small red arrowheads) assembled to form a nascent nucleus (NN).

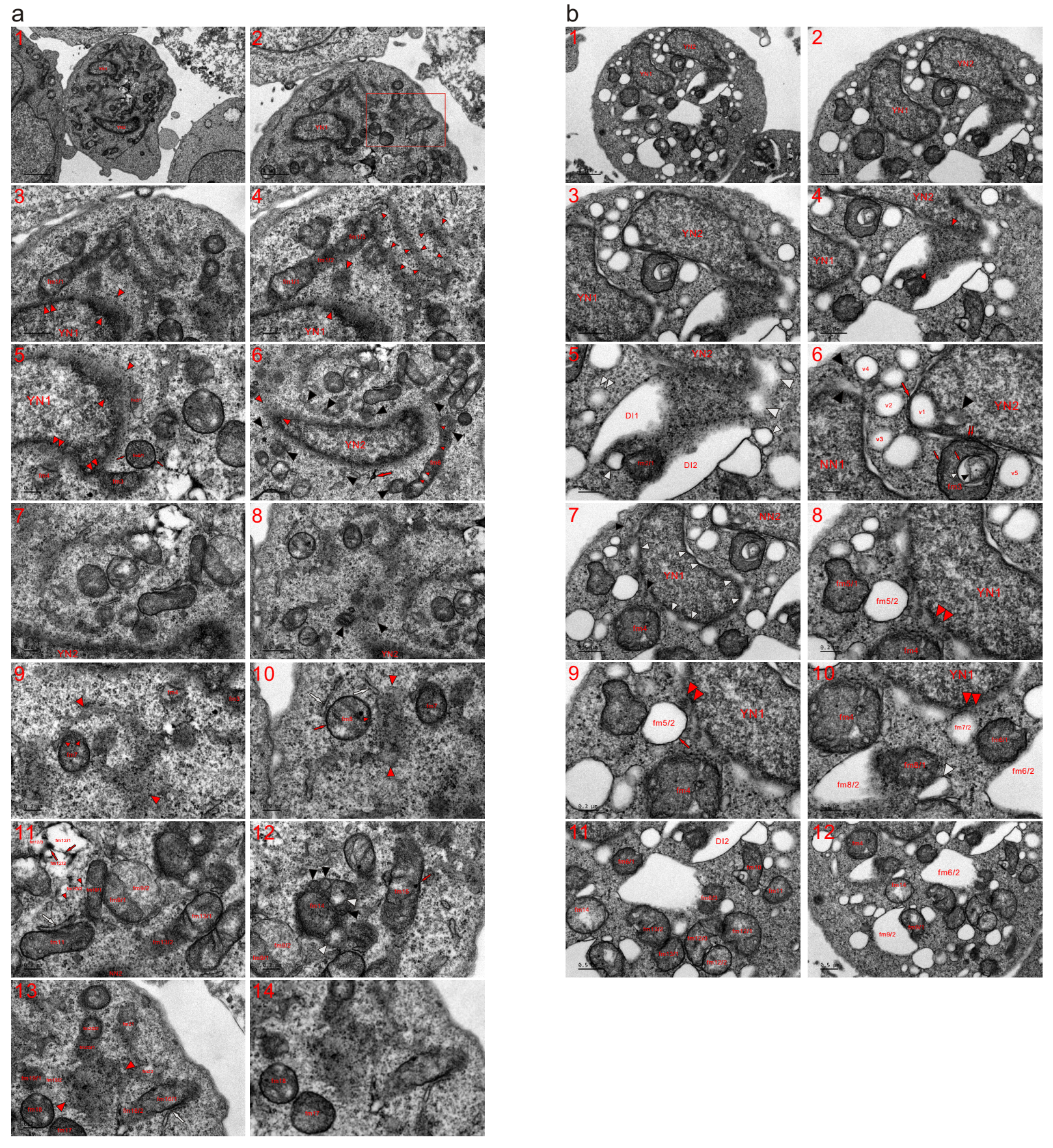

**Extended data Fig. 9. Mitochondria fragmented into dense particles to enlarge a young nucleus (YN).** TEM was performed on two K562 cells at the 4 and 12 h time points, and the micrographs revealed that mitochondria either directly dispersed into an NN in the form of dense particles or assembled the particles into the NN for nuclear growth, depending on culturing duration. (**a**) Two small or young nuclei (YN1 and YN2) appeared in the cell, and fragmented mitochondria surrounded them and dispersed into dense particles for nuclear growth. At the edge of YN1, completely fragmented mitochondria diffused directly into the nucleus in the form of dense particles (opposite red arrowheads). fm1/1 externally assembled the particles into YN1 (double red arrowheads) as well as into the surroundings, and both fm1/1 and fm1/3 were being dispersed into dense particles. Part of fm2 was aggregated into the nascent nucleus, fm2/1 was dispersed, and electron-dense fm3 and fm4 assembled dense particles into YN1 (double red arrowheads). The aggregation of the particles condensed the organelle (fm5/1), which blackened its LMM (red arrows) via the assembly of dense particles at the periphery (the external assembly), and the external assembly (small red arrowheads) formed relatively dilute areas in the cytoplasm (**a1-a5**). (**a6-a8**) Heavily or completely fragmented mitochondria (opposite red and black arrowheads, red arrow) surrounded and enlarged YN2 by providing dense particles, whose aggregation (small red arrowheads) caused the less fragmented fm6 to disperse into dense particles. (**a9** and **a10**) In the cytoplasm, most mitochondria had completely fragmented into dense particles (opposite red arrowheads), and less fragmented or condensed mitochondria (fm4, fm7 and fm8) were dispersed into the particles, whose external aggregation temporally blackened the LMM of fm8 (red arrows). Based on mitochondrial fragmentation and before dispersion into dense particles, endoplasmic reticulum-like structures (ERLs) were derived (white arrows). (**a11** and **a12**) The aggregation of dense particles separated fm9 into dense fm9/1 and electron-opaque fm9/2; the opaque part of fm10 (fm10/2) evolved into particles (small red arrowheads) while becoming more diluted. An ERL (white arrow) was separated from condensed fm11, and their assembly dispersed electron-transparent fm12, which was further divided (fm12/1-fm12/3) by the internal aggregations of dense particles (red arrows), whose congregation caused part of fm13 (fm13/2) to diffuse into NN2. In fm14, the assembly of dense particles formed SDBs (black arrowheads), which caused external aggregation to produce SDIBs (white arrowheads). Condensed fm15 lost the LMM (red arrow) via further assembly of the particles and diffused into dense particles. (**a13** and **a14**) Among the less fragmented organelles (such as fm16-fm22), a mass or an aggregate of completely fragmented mitochondria (opposite red arrowheads) appeared, which could initiate nuclear formation through further assembly of its dense particles and was enlarged by mitochondrial fragmentation or mitochondrial assembly of the particles (such as fm16/2, fm19/2, fm20/1 and fm22). At the edge of fm16, an ERL (white arrow) was formed via particle assembly. (**a14**) A high-magnification image of the inset in (**a2**). (**b**) Compared to the cell in (**a**), many electron-transparent structures or vesicles appeared in the cytoplasm of this cell. Mitochondria mainly underwent the external assembly of dense particles to expand YN1 and YN2. At the edge of YN2, mitochondria externally assembled the particles into and fused the nucleus (opposite red arrowheads) concurrently forming condensed fm2/1 and electron-lucent bodies or structures (DI1, DI2 and white arrowheads). Based on mitochondrial fragmentation, the continuous and external assembly of dense particles formed vesicles (small white arrowheads), which could derive from an SDB (double white arrowheads) (**b1-b5**). (**b6**) Mitochondrial assembly of dense particles occurred at the nuclear edge as well as within YN2 (black arrowheads), and their external aggregations led to the formation of vesicles (such as v2-v5), which fused with either (v3-v5) or both (v2) nuclei. The formation of a dark strand (large red arrow) by the external assembly of dense particles in the organelle included the mitochondrion itself in YN2, and part of the organelle transiently became a vesicle (v1). The aggregation of dense particles (small red arrow and small black arrowhead) formed lucent spots (small white arrowhead) in the condensed fm3 and caused the organelle to fuse with YN2 (double red arrows) via external particle assembly, which intermediately blackened the LMM (small red arrow). (**b7-b10**) Around YN1, the mitochondrial assembly of dense particles into the nucleus was observed (black and white arrowheads). Condensed fm4 was dispersed into the nucleus (double red arrowheads) and the surroundings. The aggregation of dense particles separated and condensed mitochondria (fm5/1, fm6/2 and fm6/3) concurrently caused parts of the organelles to become electron-transparent (fm5/2 and fm6/2); their external assembly transiently blackened LMM of fm5/2 (red arrow) and allowed the transparent bodies to fuse with YN1 (double red arrowheads). Through the further assembly of dense particles, dense fm 6/1 consequently provided the particles for fm7/2 and YN1. Near fm4, dense fm8/1 was formed by the external assembly of dense particles (fm8/2 and white arrowhead); mitochondrial assembly of dense particles formed condensed fm9/1 and electron-transparent fm9/2. Further aggregation of the particles caused these condensed mitochondria (fm10-fm14) to fragment into dense particles after first separating (fm12/1-fm12/3; fm13/1 and fm13/2), while particles in these condensed mitochondria could return back to the electron-lucent counterparts for nuclear development.

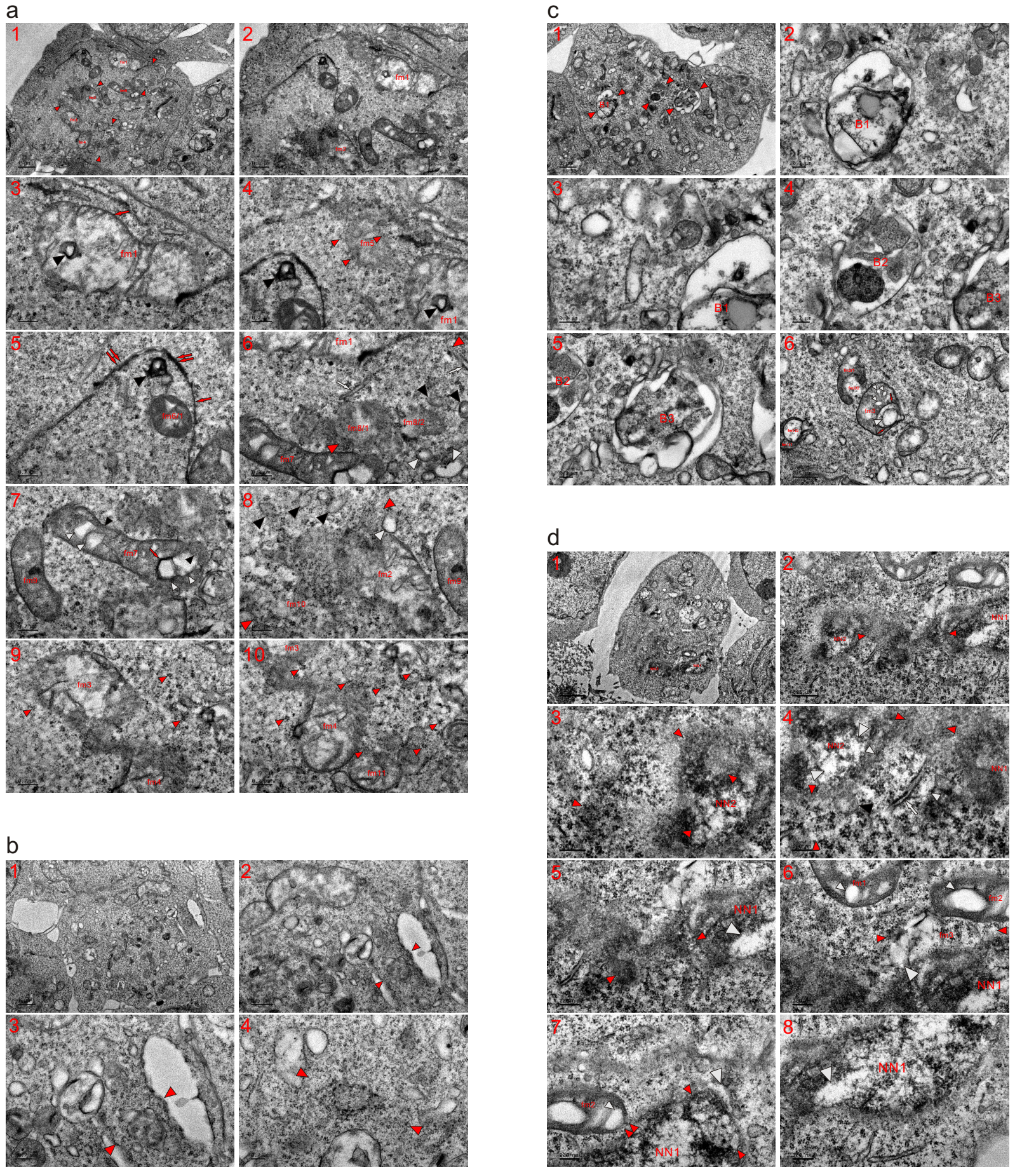

**Extended data Fig. 10. Mitochondria fragmented into dense particles to initiate nuclear formation.** TEM was performed on four K562 cells at the 2 h time point, and the micrographs showed that mitochondria fragmented into dense particles to initiate nuclear formation. (**a**) No recognizable nucleus was observed in this cell, and completely fragmented mitochondria had dispersed into dense particles. Three groups of fragmented mitochondria were indicated (opposite red arrowheads). The aggregation of dense particles diluted fm1, which was partially fragmented, almost lost its LMM (red arrow) and was dispersed into dense particles; with the dispersion of fm1, an internal aggregate of the particles appeared in the cytoplasm (red arrowheads). Adjacent to fm1, fm5 diffused into particles (small red arrowheads) whose aggregation formed dark threads or strands (red arrows), which further assembled and then disbanded (double red arrows). Part of fm6 was completely fragmented, and the condensed fm6/1 was dispersed (**a1-a5**). (**a6-a8**) Adjacent to the less fragmented organelles (such as fm1 and fm7), heavily or completely fragmented mitochondria (fm8/1, fm8/2, white arrows, black and white arrowheads) formed a group (opposite red arrowheads). Based on mitochondrial fragmentation, ERLs (white arrows), SDBs (black arrowheads) and SDIBs (white arrowheads) were intermediately derived. In dense fm7, the aggregation of dense particles (red arrow and small black arrowheads) led to the formation of lucent spots (small white arrowheads). Near fm9, which underwent partial fission, the organelles (fm2, fm10, black and white arrowheads) fragmented into each other (opposite red arrowheads). (**a9** and **a10**) In the cytoplasm, partially (such as fm3, fm4 and fm11) and completely fragmented mitochondria ultimately became dense particles (small red arrowheads), whose reassembling and rearranging could initiate nuclear formation. (**b**) No identifiable nucleus existed in this cell; two sites in the cytoplasm, where initiation of nuclear formation was likely to occur, are indicated (opposite red arrowheads) (**b1-b4**). (**c**) No recognizable nucleus was observed in the cell, and three bodies of fragmented mitochondria, which displayed the shape of a micronucleus, formed in the cytoplasm (B1-B3 and opposite red arrowheads). Both inside and outside these bodies, the mitochondrial assembly of dense particles caused most organelles to lose their characteristic morphology (**c1-c5**). (**c6**) The aggregation of dense particles led dense fm2/1 to surround lucent fm2/2, which entered the cytoplasm (fm2/2) following dispersion of the organelle, and through further assembly, the dense part of fm2 diffused (black and white arrowheads). Within fm3, the internal assembly of the particles formed dark threads (red arrows), an SDIB (large white arrowhead) and vesicular cristae (small white arrowheads). (**d**) Two nascent nuclei appeared in this cell (NN1 and NN2), and fragmented mitochondria existed between the nuclei (opposite red arrowheads) (**d1** and **d2**). (**d3** and **d4**) Around NN2, mitochondria completely fragmented into dense particles for the nuclear growth (black and opposite red arrowheads), and a mitochondrion assembled dense particles to adopt a nuclear appearance (opposite white arrowheads). Within NN2, traces of mitochondrial assembly of the particles were observed (large white arrowheads). White arrow: an ERL. (**d5-d8**) At the edge of NN1, mitochondrial aggregation of dense particles to become part of the nucleus (opposite red arrowheads) and external assembly of the particles occurred at the nuclear edge (white arrowheads) as well as within NN1 (double white arrowheads). fm1 was being dispersed into dense particles, and fm2 assembled dense particles to fuse with NN1 (double red arrowheads) and a neighbouring mitochondrion (fm3 and double red arrowheads), which was diluted, heavily fragmented and became part of the nascent nucleus (opposite red arrowheads); aggregation of the particles led to the formation of electron-transparent spots (small white arrowheads) in either fm1 or fm2.

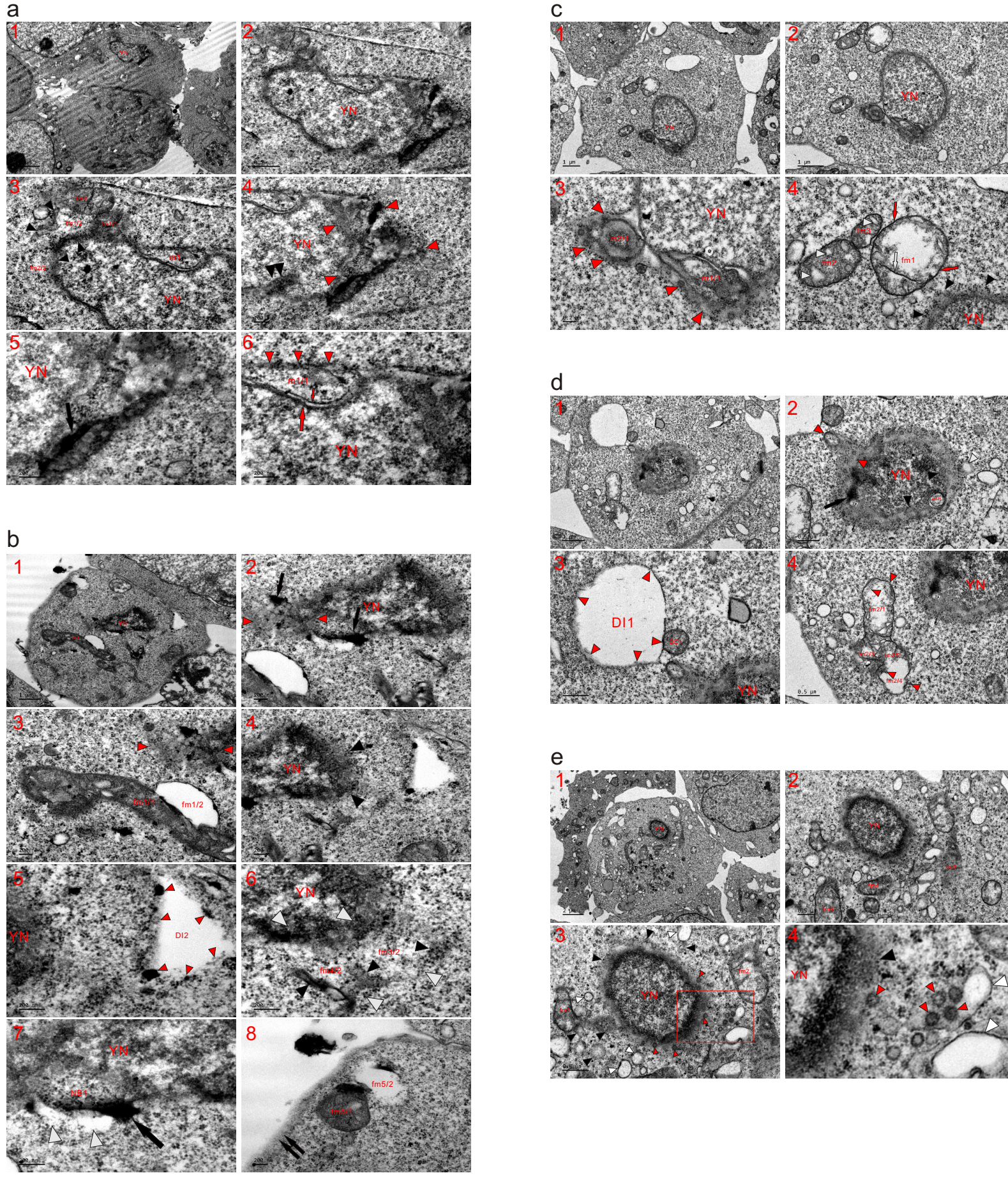

**Extended data Fig. 11. The mitochondria continuously fragmented into dense particles to enlarge a young nucleus.** TEM was performed on five K562 cells at the 2 (**a** and **b**), 6 (**c** and **d**) and 8 (**e**) time points, and the micrographs revealed that mitochondria fragmented into dense particles to promote the growth of a small nucleus or a YN. (**a**) Almost all mitochondria that surrounded the YN had been heavily or completely fragmented, and dense fm1/1 was directly dispersed into the nucleus; the external aggregation of dense particles (black arrowheads) diluted the organelles (such as fm2/1 and fm2/2), and condensed fm3 diffused into fm1 (fm1/1 and fm1/2) as well as the surroundings, providing particles for nuclear enlargement. Mitochondrial assembly of the particles occurred at the nuclear edge (opposite red arrowheads) and within the YN (double black arrowheads). Black arrow: an ink mass formed by the mitochondrial compaction of dense particles and further assembled to provide particles for nuclear growth (**a1-a5**). (**a6**) A mitochondrion assembled dense particles (large red arrow and arrowheads) to include itself into the YN, and through further congregation and linearization of dense particles, the internal aggregate (m1/1) formed a dark thread (small red arrow) at the edge. (**b**) A dense mitochondrion directly dispersed into the YN for nuclear development (opposite red arrowheads), and mitochondrial aggregations of dense particles appeared at the nuclear edge (black arrows and arrowheads). Their assembly formed an electron-dense and elongated fm1/1, leading to the formation of a lucent fm1/2 (**b1-b4**). (**b5**) External mitochondrial congregation to the fringe (red arrowheads) formed a large electron-transparent body (DI2). (**b6**) At the edge of the YN, the mitochondrial aggregation of dense particles for nuclear expansion was observed (fm3/2 and fm4/2; black and opposite white arrowheads). (**b7** and **b8**) The mitochondrial assembly of dense particles formed a nuclear body (NB1) and an ink mass (black arrow) at the edge of the YN, concurrently dispersing the lucent part of the organelle (white arrowheads). Adjacent to the plasma membrane (double black arrows), the assembly of particles separated the organelle into dense fm5/1 and transparent fm5/2. (**c**) At the edge of the YN, mitochondria externally assembled dense particles to incorporate the internal aggregates (m1/1 and m2/1) in the nucleus (**c1-c3**). (**c4**) Mitochondrial aggregation of particles into YN (black arrowheads) for its growth and their external assembly transiently blackened LMM of fm1 (red arrows); external and internal (white arrow) aggregation of dense particles caused a large part or matrix of fm1 to become electron-lucent, appearing as cristolysis. Adjacent to fm1, particle congregation led to the formation of lucent spots (white arrowheads) in condensed mitochondria (fm2 and fm3). (**d**) Mitochondria assembled dense particles to disperse into YN (opposite red arrowheads), and incomplete mitochondria-to-nucleus transition or mitochondrial aggregation of the particles occurred within the nucleus (m1/1, black arrow and arrowheads) as well as at the nuclear edge (white arrowhead) (**d1** and **d2**). (**d3**) A large mitochondrion or multiple mitochondria assembled dense particles at the peripheries (DE1 and red arrowheads) to form a large electron-lucent body (DI1). (**d4**) Near YN, the aggregation of dense particles separated fm2 (fm2/1-fm2/4) and diluted the organelle (fm2/1 and fm2/4), which then dispersed into dense particles (red arrowheads). (**e**) Mitochondria that surrounded YN were heavily or completely fragmented (black arrowheads), and virus-like granules (VLGs) (red arrowheads) developed from the dispersed mitochondria before becoming dense particles. The organelles (such as fm1-fm5) that were somewhat distant from the nucleus assembled to become particles; both SDBs (black arrowheads) and SDIBs (white arrowheads) formed based on mitochondrial fragmentation (**e1-e4**).

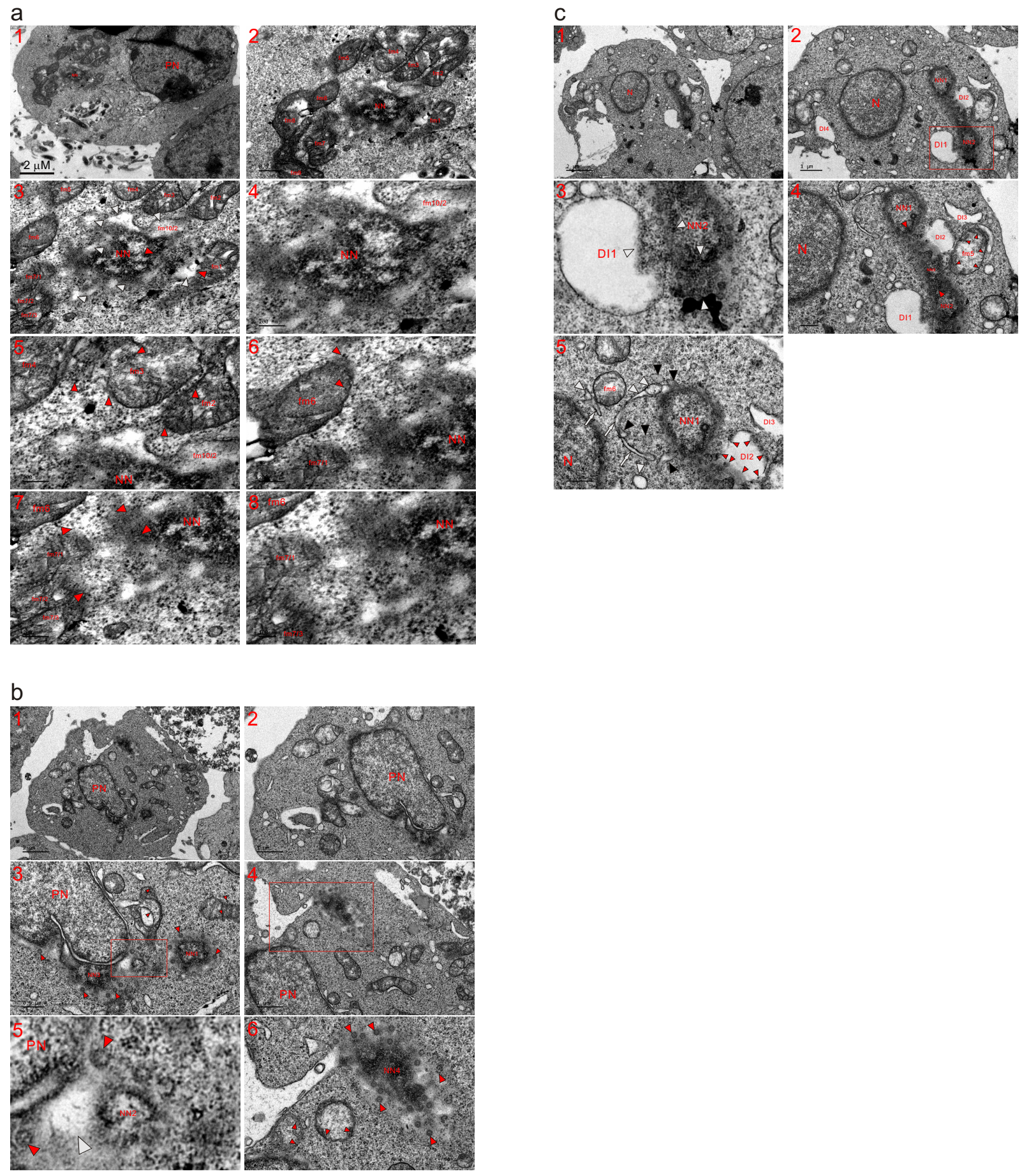

**Extended data Fig. 12. Formation of a nascent nucleus in a cell that already contained a PN or a large nucleus.** TEM was performed on three K562 cells at the 2 (**a**), 8 (**b**) and 12 (**c**) h time points, and the micrographs demonstrated that a nascent nucleus formed in the cell that already possessed a large nucleus. (**a**) In addition to the PN, a nascent nucleus (NN) was constructed and enlarged among less fragmented mitochondria (fm1-fm9), and part of fm1 dispersed into the NN (opposite red arrowheads). Within the NN and at its edge, mitochondrial assembly of dense particles led to the formation of electron-lucent or dilute structures (white arrowheads), and although the organelle was heavily fragmented, the mitochondrial origin of fm10/2 was recognizable. Through further assembly of the particles, condensed mitochondria were dispersed to continuously provide dense particles (red arrowheads) for growth of the NN (**a1-a6**). (**a7** and **a8**) Following the mitochondrial aggregation of dense particles into the nucleus, areas between the nucleus and the condensed mitochondria became relatively dilute and lacked dense particles. The dark organelles dispersed (fm7/1-fm7/3) or were in the process of dispersing (fm6) to supply dense particles for the continued growth of the NN. (**b**) In addition to a large nucleus (PN), four nascent nuclei (NN1-NN4) were formed by the mitochondrial assembly of dense particles, whose aggregation transiently formed VLGs (large red arrowheads) at the nuclear edges. MN2 looked like a mitochondrion that was assembling dense particles, whose aggregation created an electron-lucent structure between MN2 and PN. MN3 achieved fusion with PN or formed directly at the edge of PN. In the cytoplasm, the mitochondrial assembly of dense particles (small red arrowheads) for fragmentation was observed (**b1-b6**). (**b5** and **b6**) High-magnification images of the inset in (**b3**) and (**b4**), respectively. (**c**) In addition to a large nucleus (N), two small nascent nuclei (NN1 and NN2) were enlarged and connected by the mitochondrial aggregation of dense particles, whose external assembly to the peripheries caused mitochondria to become electron-lucent bodies (DI1-DI4). At the edge of NN2, a mitochondrion assembled particles into the nucleus, leading to the formation of the lucent DI1. Mitochondria assembled dense particles for nuclear development (NN3 and opposite red arrowheads), concomitantly linking NN1 and NN2. Adjacent to DI2, whose formation occurred due to the external aggregation of dense particles (small red arrowheads), fm5 assembled the particles to NN3 as well as the surroundings (small red arrowheads). At the nuclear edges and between the nuclei (N and NN1), partially (such as fm2) and heavily or completely (white arrows, black and white arrowheads) fragmented mitochondria were becoming particles, whose assembly enlarged both nuclei and ultimately let them join together.

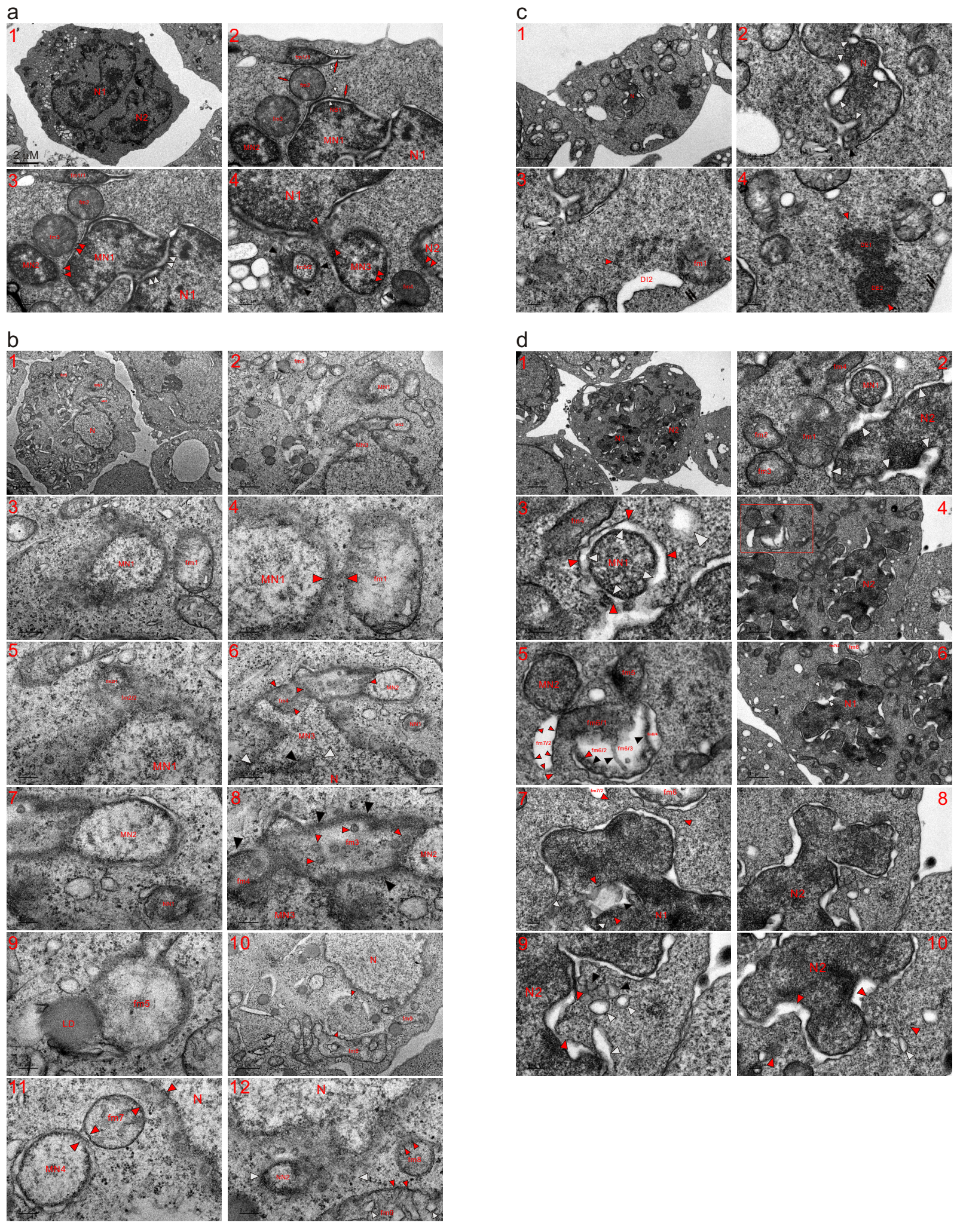

**Extended data Fig. 13. A single mitochondrion developed into a nuclear appearance or nucleus-like structure.** TEM was performed on four K562 cells at the 8 (**a** and **b**) and 12 (**c** and **d**) h time points, and micrographs showed that a single mitochondrion could develop a nuclear appearance. (**a**) In addition to two large nuclei (N1 and N2), three micronuclei appeared in the cell (MN1-MN3). Internally, the mitochondrial assembly of dense particles caused the organelle to adopt a nuclear appearance (fm1/1) and formed a dark strand (red arrow), leading to the formation of lucent structures (white arrowheads), and similar aggregation occurred at the edge of MN1 (NB1, red arrow and white arrowhead). MN2, which displayed a mitochondrial size and shape, fused with MN1 (double red arrowheads). A condensed mitochondrion (fm2) without cristae blackened its LMM (red arrow) via the further assembly of dense particles, whose aggregation linked fm1/1 with MN1 via fm2 and let dense fm3 lose the LMM to fuse with the micronucleus (double red arrowheads). Both fm2 and fm3 seemed to develop the appearance of MN2 by the reassembly or further assembly of particles, which intermixed with lucent lumens or intervals to complete the merging between micronuclei (double white arrowheads) (**a1-a3**). (**a4**) MN3 fused with N1 by assembling dense particles of condensed fm4 (opposite red arrowheads), and the dispersion of condensed fm5 (fm5/2 and black arrowheads) developed into a nuclear appearance with the help of dense particles from neighbouring mitochondria. (**b**) In addition to a large nucleus (N), which lacked tubule-NE, micronuclei (MN1-MN4) and nascent nuclei (NN1 and NN2) existed in these cells. fm1 displayed a similar density to that of MN1 and was fused with the micronucleus (opposite red arrowheads); part of a mitochondrion (fm2/1) developed a nuclear appearance with the dispersion of its other part (fm2/2) into MN2 (**b1-b5**). (**b6-b8**) Between the large nucleus and MN3, incomplete mitochondria-to-nucleus transition was demonstrated (black and white arrowheads). MN2 displayed a mitochondrial shape and looked like a mitochondrion that was assembling dense particles. The mitochondrial aggregation of dense particles (opposite red arrowheads) was observed at the edge of MN3 as well as between micronuclei (MN2 and MN3); fm3 exhibited both external (black arrowheads) and internal aggregations of dense particles, whose internal assembly formed VLGs (small red arrowheads) in the organelle. fm4 aggregated the particles (black arrowhead) to include itself in MN3 and was completely fragmented for nuclear development. Adjacent to MN2, a mitochondrion assembled dense particles to look like a nascent nucleus (NN1). (**b9**) One mitochondrion (fm5), which lacked cristae and was membraneless, fused with a lipid droplet (LD) to take in dense particles and tended to develop into a micronucleus. (**b10-b12**) Adjacent to the large nucleus, mitochondria fragmented to form a group (opposite red arrowheads), neighbouring which a large mitochondrion (fm6) appeared, and fm6 exhibited the internal and external aggregations of dense particles. Electron-opaque fm7 was dispersed into both nuclei (opposite red arrowheads) to join MN4 with the large nucleus. NN2 looked like a mitochondrion that was assembling dense particles (opposite white arrowheads); electron-opaque fm8 aggregated dense particles (small red arrowheads) into the nucleus (N) as well as the surroundings, and fm9 underwent internal (small white arrowheads) and external (small red arrowheads) aggregation of the particles, which entered NN2 and the large nucleus. (**c**) The nucleus (N) was being built by mitochondria jointly assembling dense particles, whose aggregation was observed within the nucleus and at its edge (black and white arrowheads). Adjacent to the plasma membrane (double black arrows), the mitochondrial assembly of dense particles was observed (fm1, DI2 and opposite red arrowheads) (**c1-c3**). (**c4**) Next to the plasma membrane (double black arrows), mitochondria assembled the particles to form dense bodies (DE1, DE2 and opposite red arrowheads), which appeared as a group of chromatins and tended for nuclear development via further assembly inside dense particles (**c1-c4**). (**d**) The same cell shown in **Figure 3c.** The large nuclei (N1 and N2) appeared to be constructed by continuous nuclear transition followed by the fusion of mitochondria or a mitochondrion. MN1 looked like the internal assembly of dense particles of a mitochondrion, and between the internal (MN1) and external (red arrowheads) aggregation of the particles, a lunar halo structure (white arrowheads) formed, which also existed at the edges of both N1 and N2 (white arrowheads). Condensed mitochondria (fm1-fm4) were dispersed into dense particles, whose external assembly diffused into an SDIB (large white arrowhead) in the cytoplasm (**d1-d3**). (**d4** and **d5**) Compared to MN1, both MN2 and fm5 were less advanced regarding nuclear development; MN2 tended to develop a similar appearance to MN1 via the further assembly of dense particles, and fm5 appeared as a nascent nucleus. Near MN2 and fm5, mitochondria exhibited the aggregation of dense particles (fm6/1 and fm6/2-fm6/4; fm7/2), and their internal assembly (black arrowheads) further divided the electron-lucent part of fm6 (fm6/2-fm6/4). External aggregation (small red arrowheads) caused fm7/2 to become electron-lucent and formed an electron-opaque aggregate between the organelles (large red arrowheads). (**d6** and **d7**) Mitochondria (such as fm6 and fm7) assembled dense particles into N1 (opposite red arrowheads), and incomplete mitochondria-to-nucleus transition or mitochondrial aggregation of dense particles was observed within the nucleus (opposite red and black arrowheads) and at the nuclear edge (opposite white arrowheads). (**d8-d10**) At the edge of N2, SDBs (black arrowheads) and SDIBs (white arrowheads) formed based on mitochondrial fragmentation, and mitochondria-derived dense particles tended for nuclear development (opposite red arrowheads).

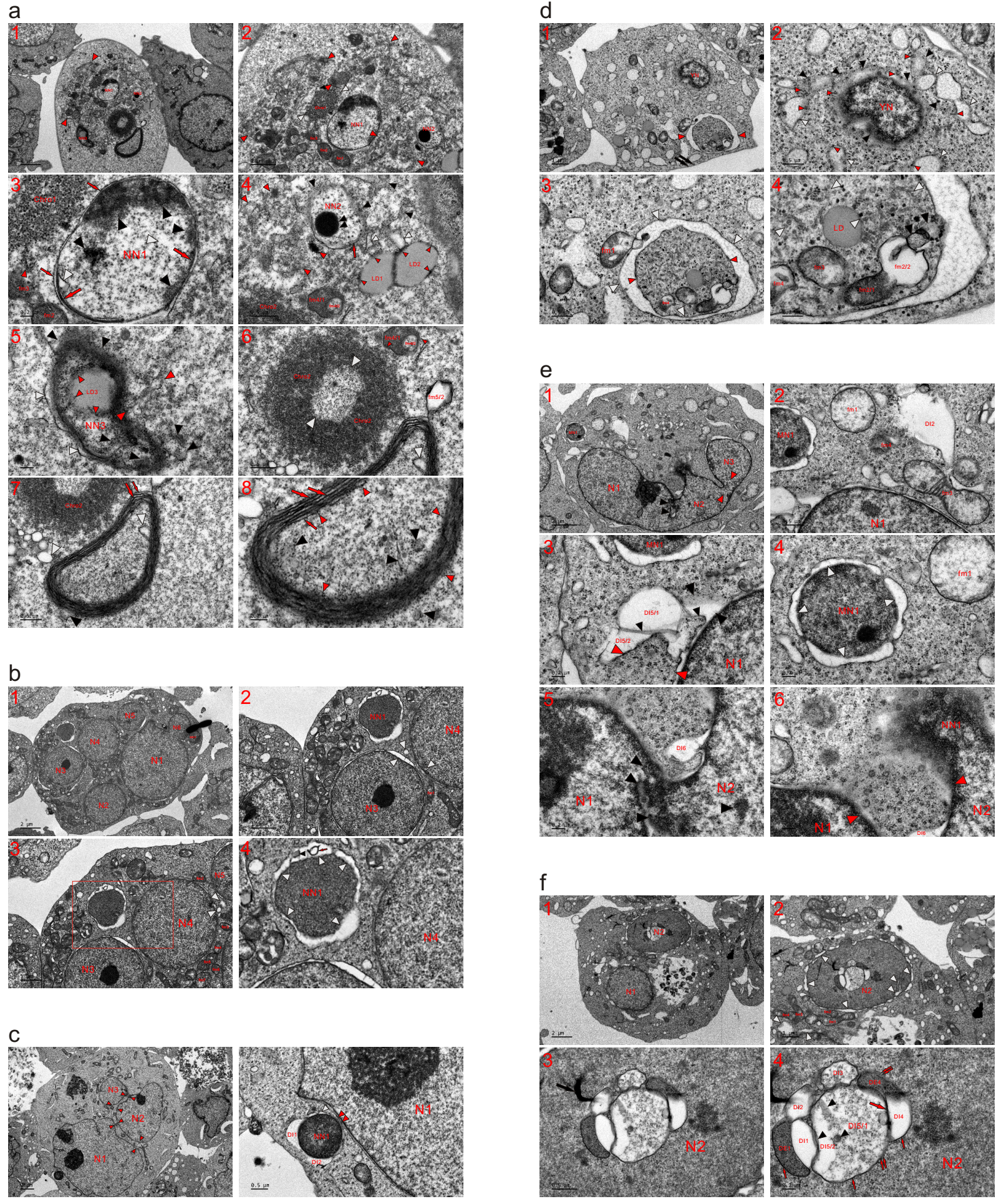

**Extended data Fig. 14. Mitochondria assembled dense particles together to develop into a small nucleus (or an MN).** TEM was performed on six K562 cells at the 4 (a), 6 (**b**), 8 (**c-e**) and 12 (**f**) h time points, and the micrographs showed that mitochondria assembled dense particles together for nuclear development. (**a**) Three bodies with nuclear appearance were labelled as the nascent nuclei (NN1-NN3). Near NN1, the mitochondrial aggregation of dense particles formed a group of chromatins (Chro1 and opposite white arrowheads). In the cytoplasm, joint mitochondrial assembly of dense particles is indicated (opposite red arrowheads). Adjacent to NN1, condensed mitochondria (such as fm1-fm3) were dispersed into the particles (**a1** and **a2**). (**a3**) Within NN1 and at its edge, the mitochondrial assembly of dense particles was observed (large and small red arrows; black arrowheads), leading to the formation of electron-lucent structures or spots (white arrowheads), and condensed fm3 diffused into the chromatin group (Chro1) in the form of dense particles (red arrowhead). (**a4**) NN2 was diluted by the aggregation of the particles (black arrowheads and red arrow), whose internal assembly formed a Nu-like body (double black arrowheads). Near NN2, mitochondria aggregated dense particles to fragment (black, white and opposite red arrowheads). Not far from the nascent nucleus, two lipid droplets (LD1 and LD2) were assembled to become particles (small red arrowheads). (**a5**) The mitochondrial assembly of dense particles (black, white and opposite red arrowheads) displayed the appearance of a nascent nucleus (NN3), within which LD3 was turning into particles (small red arrowheads). (**a6-a8**) Mitochondria-derived dense particles assembled to form a group of chromatin (Chro2), leading to the formation of a central area (opposite white arrowheads) in Chro2. The aggregation of the particles separated fm5 (dense fm5/1 and dilute fm5/2), which was dispersed into Chro2 as well as the surroundings in the form of dense particles (small red arrowheads). Mitochondria assembled the particles to form dark strands (large red arrows) or threads (small red arrows), which combined to display an onion-like structure. Within the structure and at its edge, traces of mitochondrial fragmentation were exhibited (such as fm5/2, black and white arrowheads). Through further aggregation, each dark strand or thread dispersed to become dense particles (small red arrowheads), whose reassembly could initiate nuclear formation. (**b**) Six relatively large nuclei (N1-N6) appeared in the cell. N1 achieved fusion with N6, which merged with MN1. Between the nuclei, mitochondria fragmented (such as fm1-fm7 and white arrowheads) into dense particles for nuclear development, leading to the joining together of these individually formed nuclei. Several mitochondria externally assembled dense particles to construct an NN, leading to the formation of a lunar halo structure (white arrowheads) around the nascent nucleus; within the lunar halo, internal aggregation of dense particles appeared (red arrow and small black arrowhead) (**b1-b4**). (**c**) Separately formed nuclei (N1-N3) were joined together by the mitochondrial assembly of dense particles (opposite red arrowheads), whose aggregation formed NN1 and fused the nascent nucleus with N1 (double red arrowheads), concurrently leading to the formation of electron-lucent structures or bodies (DI1, DI2 and white arrowheads) (**c1** and **c2**). (**d**) In addition to a young nucleus (YN), a group of fragmented mitochondria (opposite red arrowheads) appeared adjacent to the plasma membrane (double black arrows). Around the YN, mitochondrial aggregations of dense particles were observed (black arrowheads), and somewhat distant from the YN, either SDBs or SDIBs were becoming particles (small red arrowheads) (**d1** and **d2**). (**d3** and **d4**) Around an electron-opaque aggregate (opposite red arrowheads) of fragmented mitochondria, the congregation of dense particles from the connected mitochondria formed a lunar halo structure (white arrowheads), which was fused with condensed fm1. Within the aggregate, which tended to develop a nuclear appearance, partially (such as fm2/1, fm2/2, fm3 and fm4) or completely (black and opposite white arrowheads) fragmented mitochondria and a lipid droplet (LD) were present. (**e**) Partial nuclear fusion (opposite red arrowheads) and incomplete mitochondria-to-nucleus transition (black arrowheads) were observed. The mitochondrial aggregation of dense particles diluted fm1, formed a large electron-transparent body (DI2), diffused fm3 into N1 and heavily fragmented fm4 into particles. At the edge of N1, the mitochondrial assembly of dense particles was demonstrated (black and opposite red arrowheads), concurrently leading to the formation of electron-lucent bodies or structures (DI5 and white arrowhead), and their internal aggregation (small black arrowhead) further divided DI5 (DI5/1 and DI5/2) (**e1-e3**). (**e4**) An aggregate of mitochondria-derived dense particles developed into a micronucleus (MN1), around which electron-transparent structures connected to form a lunar halo (white arrowheads). (**e5** and **e6**) Between nuclei and within N2, incomplete mitochondria-to-nucleus transition or mitochondrial aggregation of dense particles was observed (black arrowheads), and their external assembly created electron-transparent DI6. Immediately next to DI6, mitochondria-derived particles reassembled for nuclear development (opposite red arrowheads). At the edge of N2, dense particles assembled into the appearance of a nascent nucleus (NN1), whose formation consequently enlarged N2. (**f**) Two nuclei were under construction, and partially or completely fragmented mitochondria existed between them (such as fm1-fm4 and white arrowheads). Within N2 and at its edge, the mitochondrial aggregation of dense particles led to the formation of electron-lucent structures (white arrowheads) (**f1** and **f2**). (**f3** and **f4**) Within N2, aggregations of dense particles appeared (DE1 and DE4; black and red arrowheads; black arrowheads), concurrently leading to the formation of electron-transparent or electron-dilute bodies (DI1-DI5), one of which was divided (DI5/1 and lucent DI5/2) by the internal assembly of dense particles. Their external aggregation and linearization blackened the LMMs (small red arrows), which dispersed through further assembly (double small red arrows); these aggregates of dense particles eventually became particles for nuclear transition.

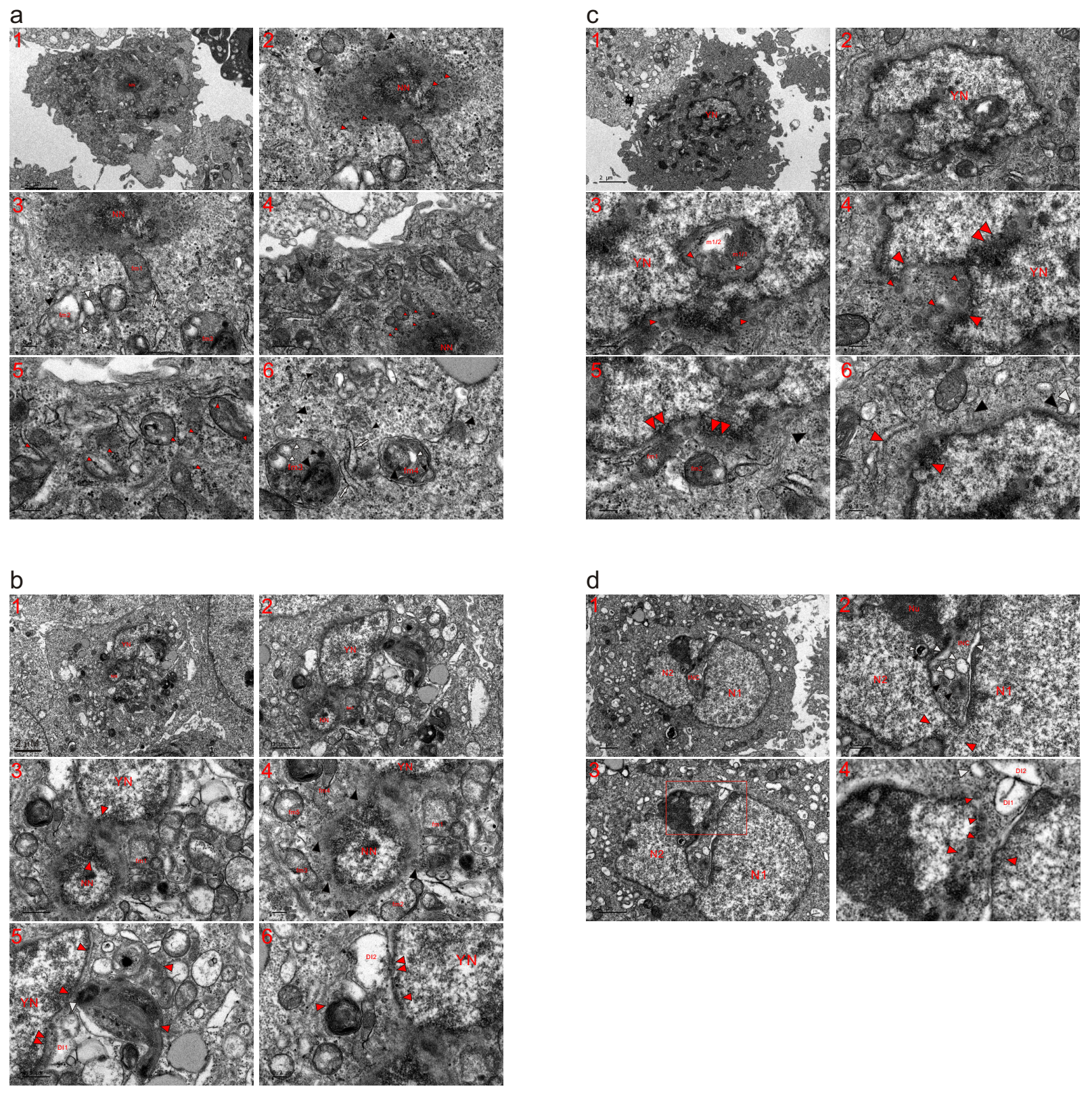

**Extended data Fig. 15. Mitochondria fragmented into dense particles to initiate nuclear formation and construct nuclei in HepG2 cells.** TEM was performed on four HepG2 cells at the 3 h time point, and the micrographs revealed that mitochondria fragmented into dense particles to initiate nuclear formation and promote nuclear growth. (**a**) A nascent nucleus (NN) was constructed by the mitochondrial assembly of dense particles, and part of fm1 dispersed into the NN. At the nuclear edge, VLGs (red arrowheads) were derived from complete mitochondrial fragmentation. Immediately next to the NN, SDBs (black arrowheads) diffused into it in the form of dense particles. Based on the heavy fragmentation of fm2, an SDB (black arrowhead) and SDIBs (white arrowheads) were derived. At the edge of fm1, an ERL (white arrow) was formed by particle assembly (**a1-a3**). (**a4**) At the edge of NN, mitochondria became dense particles (small red arrowheads) to enlarge the nucleus. (**a5** and **a6**) In the cytoplasm, mitochondria assembled dense particles to disperse into particles (small red arrowheads) whose internal aggregations (white, large and small black arrowheads) caused the organelles to transiently appear as MVBs (fm3 and fm4). Based on mitochondrial fragmentation, the aggregation of dense particles formed small and large SDBs as well as ERLs (white arrows) in the cytoplasm (small and large arrowheads). (**b**) In addition to a young nucleus (YN), a nascent nucleus (NN) appeared and fused with the YN by the mitochondrial assembly of dense particles (opposite red arrowheads). Partially (such as fm1-fm5) or completely (black arrowheads) fragmented mitochondria surrounded the NN and promoted its growth (**b1-b4**). (**b5** and **b6**) At the edge of YN, mitochondria assembled the particles into the nucleus as well as the surroundings. The external aggregation of dense particles caused the organelles to fuse with YN (double red arrowheads), leading to the formation of electron-lucent or electron-dilute structures (DI1, DI2 and white arrowhead). (**c**) The nucleus was constructed by mitochondrial fragmentation into dense particles, and incomplete mitochondria-to-nucleus transition was exhibited within the nucleus (dense m1/1, lucent m1/2 and double red arrowheads) and at its edge (opposite red arrowheads). Based on mitochondrial fragmentation, VLGs were derived (small red arrowheads) (**c1-c4**). (**c5** and **c6**) Part of the organelles dispersed into the YN (fm1, fm2 and double red arrowheads), and heavily or completely fragmented mitochondria assembled dense particles into the nucleus (opposite red, black and white arrowheads) to promote nuclear growth. (**d**) Individually formed nuclei (N1 and N2) accomplished partial nuclear fusion (opposite red arrowhead), which consequently partitioned the cytoplasm to form an opened INC. At the edge of the Nu, incomplete mitochondrion-to-nucleus transition was observed (small black and white arrowheads). Within the INC, mitochondrial aggregation of dense particles was demonstrated (black and white arrowheads), and at its opening, the assembly of the particles caused mitochondria to become electron-transparent bodies (DI1, DI2 and white arrowhead) and sealed the INC (opposite and small red arrowheads) (**d1-d4**). (**d4**) A high-magnification image of the inset in (**d3**).

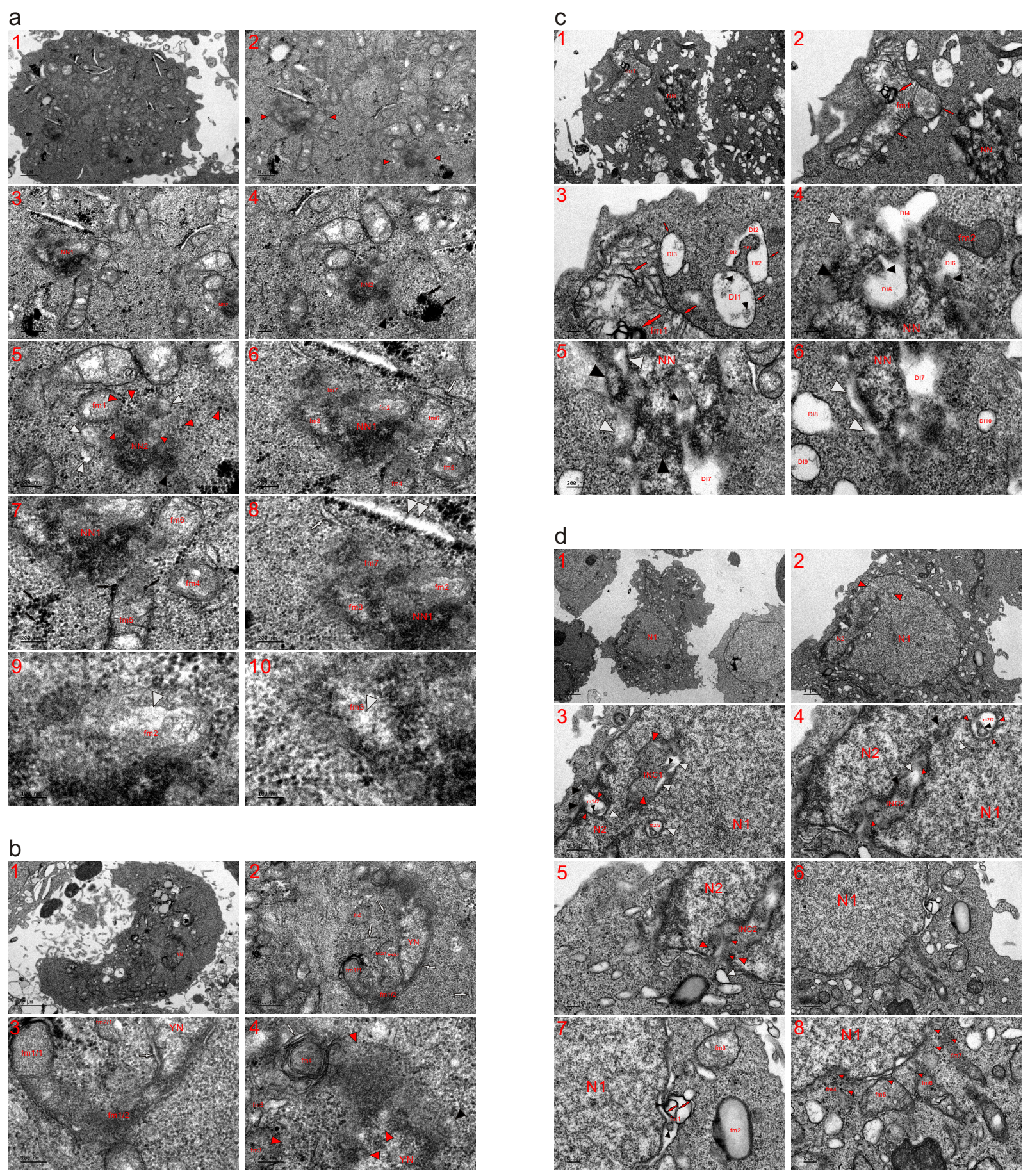

**Extended data Fig. 16. Mitochondria fragmented to initiate nuclear formation and build nuclei in HeLa cells.** TEM was performed on four HeLa cells at the 3 (**a** and **b**) and 6 (**c** and **d**) h time points, and the micrographs showed that mitochondria participated in nuclear initiation and building through the assembly of dense particles to fragment. (**a**) Two nascent nuclei (NN1, NN2 and opposite red arrowheads) appeared in the cells. NN2 looked like a dispersed dense mitochondrion and was surrounded by fragmented mitochondria (such as fm1; black, white and small red arrowheads). Large red arrowheads: dense particles, which were derived from mitochondrial fission; black arrows: ink masses, which were formed by the mitochondrial aggregation of dense particles (**a1-a5**). (**a6-a10**) Although heavily fragmented, the mitochondrial origins of the organelles were recognizable (fm2 and fm3), and they dispersed into each other to form NN1 (fm2, fm3 and fm7), which was enlarged and expanded by partially fragmented mitochondria (such as fm4-fm6). Assembly of the particles formed transparent spots (white arrowheads) in fm2 and fm3. Adjacent to NN1, the mitochondrial aggregation of dense particles formed ERLs (white arrowheads) and an elongated electron-lucent structure (double white arrowheads). Black arrows: glycogen-like granules formed by the mitochondrial assembly of dense particles. (**b**) A young nucleus was present in the cell, and mitochondria assembled dense particles to disperse into the nucleus (fm1/2 and fm2/2). At the nuclear edge, partially or completely fragmented mitochondria were observed (fm3-fm5, opposite red and black arrowheads). White arrows: ERLs, which were derived from mitochondrial aggregation of the particles. (**c**) A nascent nucleus (NN) was being built by mitochondria assembling dense particles. Near the nucleus, there appeared to be a large mitochondrion, which was larger than that of the NN and dispersed via the internal (large and small red arrows) and external (small red arrows) aggregation of particles whose congregations (DE2, red arrows and black arrowheads) caused the mitochondria to become electron-transparent bodies (DI1-DI3) (**c1-c3**). (**c4-c6**) Within the NN and at its edge, the mitochondrial assembly of dense particles was observed (large and small black arrowheads), and their aggregations led to the formation of electron-lucent bodies or structures (DI4-DI7 and white arrowheads). SDIBs were derived from mitochondrial fragmentation, and some of them looked like vesicles (DI9 and DI10), which ultimately diffused into the cytoplasm via the external aggregation of dense particles (such as DI8). (**d**) Nuclear fusion between nuclei (N1 and N2) partitioned the cytoplasm to form an INC, which was further divided into the closed INC1 and open INC2 following mitochondria-to-nucleus transition in the middle of nuclei (opposite white arrowheads). Part of a mitochondrion (lucent m1/2) was included in N2 as it assembled dense particles (large black and small red arrowheads) for nuclear development, and internal aggregation of the particles was observed in m1/2 (small black arrowhead). Part of the organelle became the nuclear-localized organelle (m2/2) in N1 along with the nuclear transition of itself and its neighbouring counterparts, and the internal (black arrowheads) and external (small red arrowheads) aggregation of dense particles caused m2/2 to be electron-lucent. By further assembling dense particles, the aggregate (small black arrowhead) dispersed within m2/2, which consequently disappeared (large black and white arrowheads). Within INC1, the mitochondrial assembly of dense particles (opposite red and small black arrowheads) for nuclear development led to the formation of electron-transparent intervals (white arrowheads). In INC2, mitochondria-derived dense particles reassembled for nuclear transition (black, white and small red arrowheads) and the sealing of the INC (opposite and small red arrowheads, and white arrowhead) (**d1-d5**). (**d6-d8**) At the edge of N1, the aggregation of dense particles caused a mitochondrion to become electron transparent, and the internal assembly of dense particles (red arrows and black arrowheads) further divided the organelle. The aggregation of dense particles caused fm2 to look like a lipid droplet and caused fm3 to disperse; partially (such as fm5-fm7) or completely (fm4) fragmented mitochondria diffused into N1 as well as the surroundings in the form of dense particles (small red arrowheads).

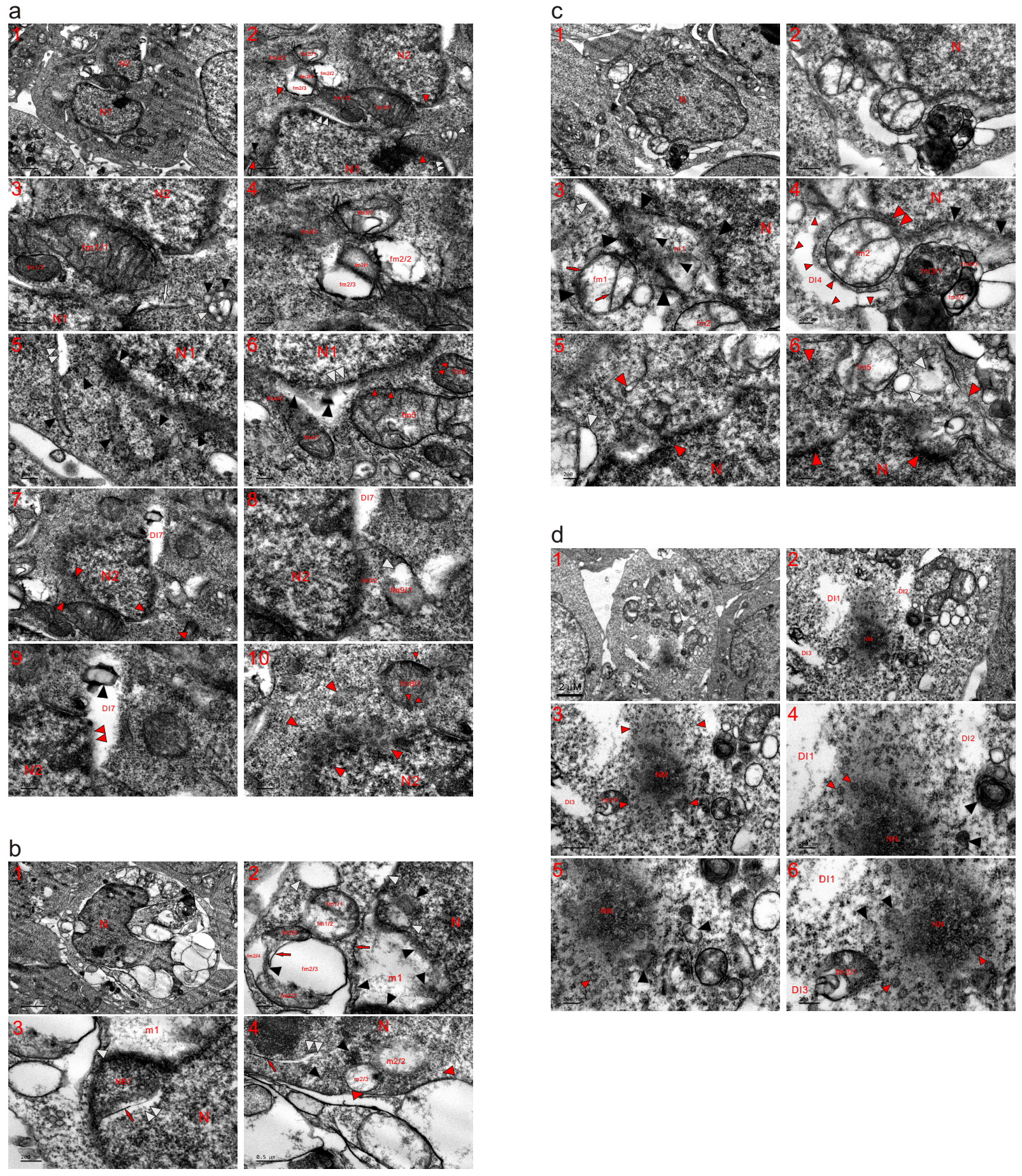

**Extended data Fig. 17. Mitochondria fragmented to initiate nuclear formation and construct nuclei in HEK293T cells.** TEM was performed on four HEK293T cells at the 3 (**a-c**) and 6 (**d**) h time points, and the micrographs revealed that mitochondria fragmented into dense particles to initiate nuclear formation as well as to build a nucleus. (**a**) Partially (fm1 and fm2) or completely (opposite red arrowheads) fragmented mitochondria were present between two individually constructed nuclei (N1 and N2) and at the edge of N1, and traces of mitochondrial fragmentation were observed (white arrow, black and white arrowheads). The mitochondrial assembly of dense particles separated fm1 (fm1/1-fm1/3), caused a large part of fm2 to become electron transparent (fm2/2 and fm2/3) and dispersed part of fm3 into dense particles (fm3/2), and their aggregations created electron-lucent structures at the edge of N1 (double white arrowheads). Mitochondrial aggregations of dense particles into N1 were observed (fm4/2 and black arrowheads), and electron-opaque mitochondria (fm5 and fm6) were dispersed into the nucleus as well as the surroundings in the form of dense particles (small red arrowheads) (**a1-a6**). (**a7-a10**) At the edge of N2, a mitochondrion became electron-lucent (DI7) and fused with the nucleus (double red arrowheads) as it assembled dense particles to N2 as well as the surroundings; in DI7, internal aggregation of the particles was observed (black arrowhead). Part of fm8 dispersed, and the remaining counterpart (fm8/1) was being assembled into dense particles (small red arrowheads). Completely fragmented mitochondria diffused into N2 (opposite red arrowheads) to promote its growth in the form of dense particles, whose aggregation caused fm9/2 to become part of the nucleus, and the internal congregation of the particles formed lucent spots in fm9/1. (**b**) A nucleus (N) was enlarged by the mitochondrial assembly dense particles, whose aggregation caused the nucleus to be surrounded by electron-transparent bodies. Electron-lucent m1 was incorporated in the nuclear edge as it externally assembled the particles to its periphery (red arrow and black arrowheads). Adjacent to m1 and within the nucleus, mitochondrial aggregation of dense particles was observed (black and white arrowheads). Near m1, the mitochondrial assembly of dense particles formed an electron-opaque nuclear body (NB1) and dark thread, concurrently forming a lucent structure (double white arrowheads). At the nuclear edge and in the cytoplasm, their congregation created transparent bodies or structures (white arrowheads) and separated mitochondria (fm1/1 and fm1/2; fm2/1-fm2/4), while their internal aggregations (black arrowhead and red arrow) further divided the electron-transparent part of fm2 (fm2/3 and fm2/4) (**b1-b3**). (**b4**) Within the nucleus, mitochondrial aggregations of the particles (red arrow, opposite red and black arrowheads) caused the mitochondria to become electron-transparent bodies or structures (m2/2, m2/3 and double white arrowheads), which looked like a vesicle (m2/3). (**c**) Mitochondria fragmented into dense particles to promote the growth of a nucleus (N), and a mitochondrion (m1) externally assembled particles to its periphery (large black arrowheads) while becoming incorporated into the nucleus; in m1, the internal aggregation of dense particles was observed (small black arrowheads). At the nuclear edge, the mitochondrial assembly of dense particles to the nucleus as well as the surroundings formed an electron-lucent structure (white arrowheads), and their congregations (red arrows and black arrowhead) caused fm1 to become transparent (**c1-c3**). (**c4**) A mitochondrion (fm2) aggregated the particles into the nucleus, leading to its fusion with the nucleus (double red arrowheads), and completely fragmented mitochondria appeared at the edge of the nucleus (black arrowheads). The internal aggregations of dense particles caused fm3 to look like an MVB (fm3/1-fm3/3), and their external assembly (small red arrowheads) formed lucent DI4. (**c5** and **c6**) At the edge of the nucleus, partially (such as fm5) and heavily or completely (opposite red and white arrowheads) fragmented mitochondria became dense particles for nuclear development. (**d**) In this cell, a nascent nucleus (NN) appeared, and mitochondrial aggregation of dense particles (opposite red arrowheads) expanded its size, leading to the formation of large electron-lucent structures. Heavily or completely fragmented mitochondria neighboured the nucleus (fm3/1, black and white arrowheads), and based on the dispersed mitochondria, VLGs (small red arrowheads) appeared and disappeared as they enlarged the NN.

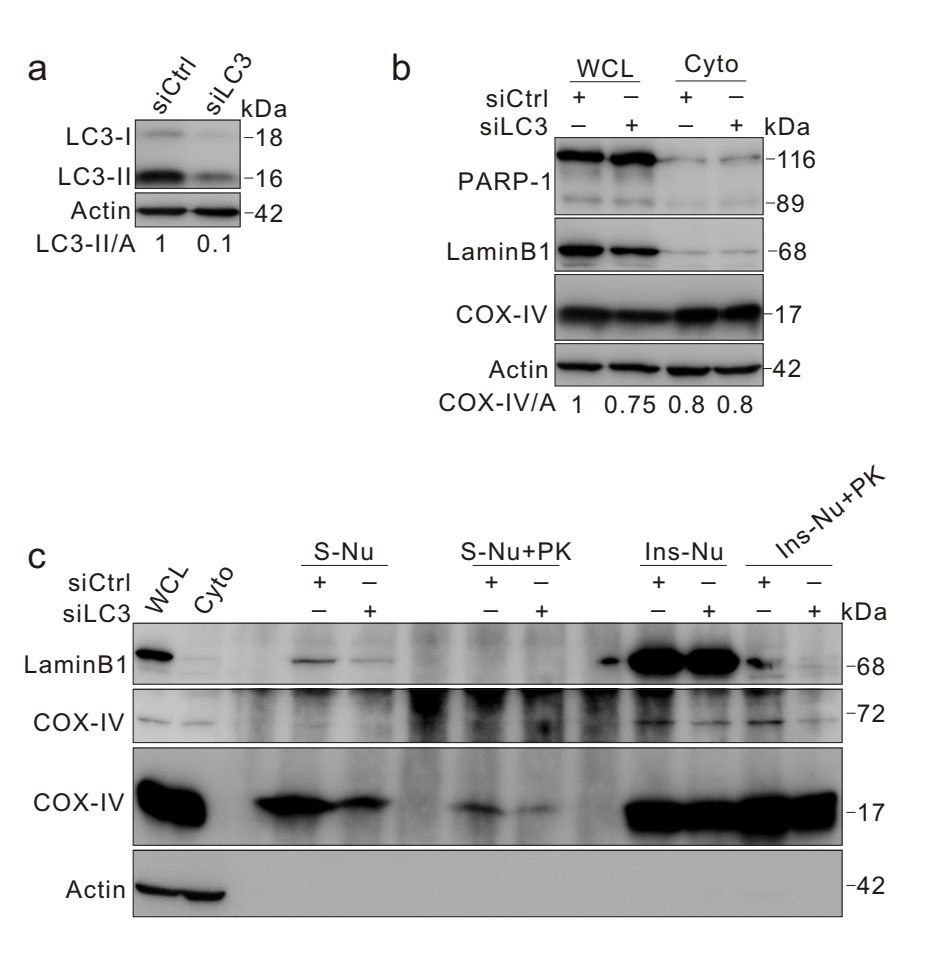

**Extended data Fig. 18. Knockdown of LC3 markedly decreased the level of COX-IV in the soluble nuclear fraction.** K562 cells were transfected with either small interfering RNA (siRNA) against LC3 (siLC3) or with a control (siCtrl) for 48 h, and thereafter, cells were gathered and cultured with new medium for 2 h before subcellular fractionation; nuclear fractions were treated with or without proteinase K (PK, 200 ng/ml) for 30 min at room temperature, and whole cell lysate (**a**) and subcellular fractions were immunoblotted with the indicated antibodies. S-Nu: soluble nuclear fraction; Ins-Nu: insoluble nuclear fraction; WCL: whole cell lysate; Cyto: cytoplasmic fraction.

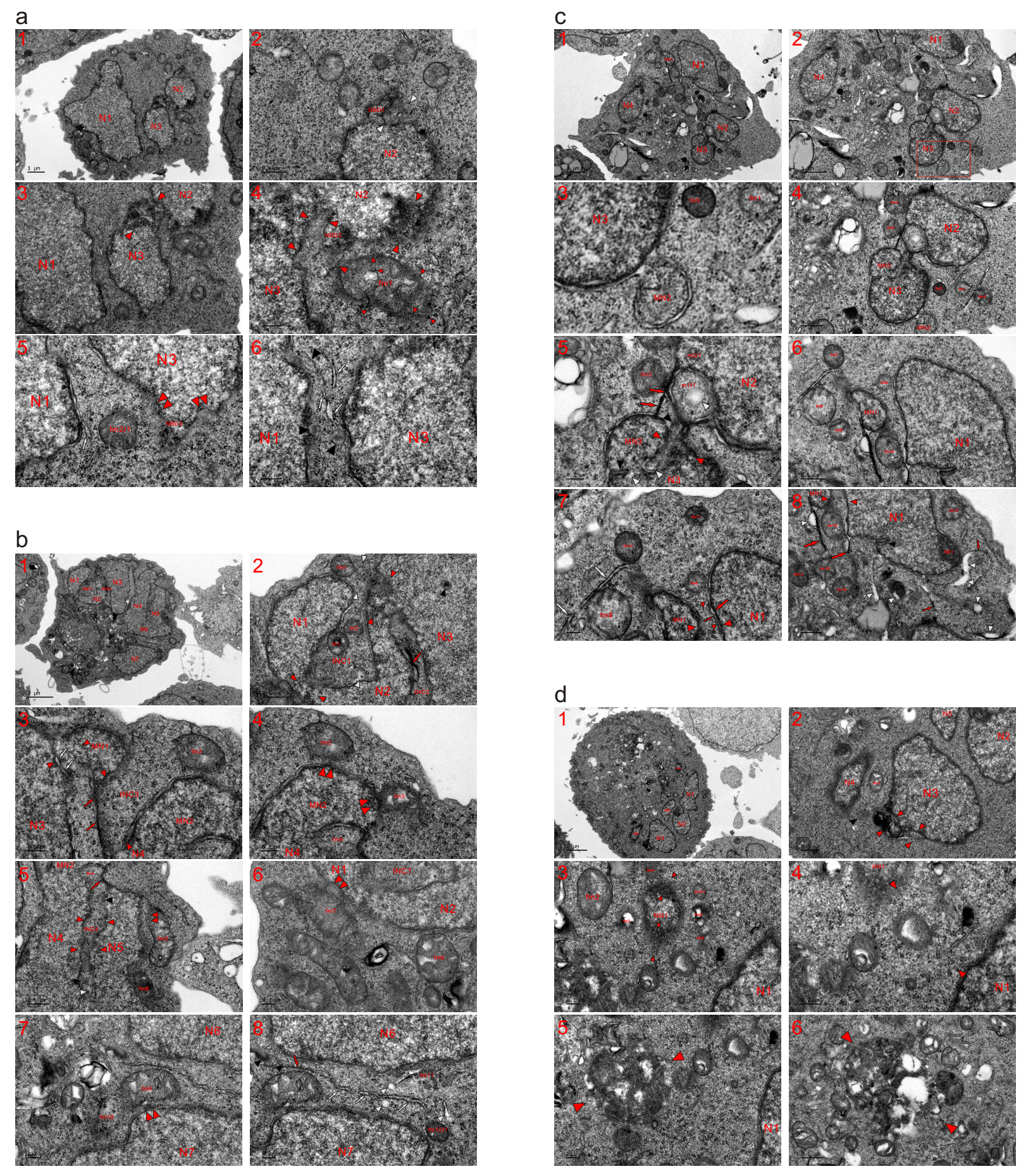

**Extended data Fig. 19. Knockdown of LC3 delayed nuclear fusion of individually formed nuclei in a single cell.** TEM was performed on four K562 cells with (**c** and **d**) or without (**a** and **b**) depletion of LC3 at the 2 h time point, and the micrographs revealed that, compared to the control (**a** and **b**), LC3 loss delayed and obstructed the nuclear fusion of individually formed nuclei in a single cell. (**a**) Three individually formed nuclei (N1-N3) appeared in this control cell, and partial nuclear fusion was achieved between N2 and N3 by the mitochondrial assembly of dense particles for nuclear development (opposite red arrowheads). At the edge of N2, the mitochondrial assembly of dense particles formed a nuclear bud (NBD1), whose formation caused the incomplete nuclear conversion of a mitochondrion to look like an INC (opposite white arrowheads) (**a1-a3**). (**a4**) Between nuclei and at the nuclear edge, mitochondrial aggregation of dense particles for nuclear development was observed (NBD2 and opposite red arrowheads). Adjacent to the nuclei (N1 and N2), a less fragmented and condensed mitochondrion (fm1) was being dispersed into dense particles (small red arrowheads) to enlarge both N1 and N2. (**a5** and **a6**) Between N1 and N3, most mitochondria had completely fragmented into dense particles; traces of mitochondrial fragmentations were recognized (white arrows and black arrowheads), and the remaining part of dense fm2 (fm2/1) diffused into the particles. At the edge of N3, mitochondria assembled dense particles to expand the nuclear size (NBD3 and double red arrowheads). (**b**) In addition to N7, most separately constructed nuclei (N1-N6) had accomplished nuclear fusion in this control cell, and N6 and N7 were joined together by partially or completely fragmented mitochondria (**b1**). (**b2**) Mitochondria assembled dense particles to achieve nuclear transition, leading to partial nuclear fusions (opposite red arrowheads), which in turn partitioned the cytoplasm to form intranuclear inclusions (INC1 and INC2). Within the INCs, partially (m1 and m2) and heavily (red arrow and white arrowhead) fragmented mitochondria were observed, and most of the organelles had completely fragmented into particles. The mitochondrial assembly of dense particles for nuclear development (fm1 and opposite white arrowheads) at the opening of the INC sealed INC1. In N3, incomplete mitochondrion-to-nucleus transition was observed (black arrowhead). (**b3** and **b4**) At the edges of the large nuclei, the mitochondrial assembly of dense particles formed micronuclei (MN1 and MN2) and enlarged both nuclei (N3 and MN1), merging the micronuclei with these large nuclei (MN1 and N3; MN2 and N4) (opposite red arrowheads); between N3 and MN1, incomplete nuclear conversion of a mitochondrion was observed (white arrows). The linkage of MN1 with N4 (red arrows) by the congregation of dense particles compartmentalized the cytoplasm to form INC3, in which mitochondria completely fragmented into dense particles for nuclear development (opposite red arrowheads). At the edge of MN2, less fragmented mitochondria (fm2 and fm3) dispersed into it as well as the surroundings in the form of dense particles. With the nuclear transition of fm4 and its neighbouring counterparts, fragmented fm4 was included between MN2 and N4. (**b5** and **b6**) Mitochondrial aggregation of particles linked MN2 with N5 while simultaneously closing INC4, within which mitochondria-derived dense particles assembled for nuclear development (opposite red, black and white arrowheads) to make the whole INC disappear. At the nuclear edges, less fragmented mitochondria assembled the particles to partially (fm5 and fm7) or totally (fm6) fuse with nuclei (double red arrowheads), and fm8 aggregated dense particles to diffuse into the N2. (**b7** and **b8**) fm9 was dispersed into N7 as well as the surroundings, and the mitochondrial origin of some completely fragmented mitochondria (fm10 and fm11) was recognized at the nuclear edges. Between N6 and N7, although most of the organelles had dispersed into dense particles for nuclear development, traces of mitochondrial fragmentation were observed (fm12/1; red and white arrows; black and white arrowheads). (**c**) Compared to the nuclei in control cells, nuclei (N1-N4) in this LC3-depleted cell were widely separated, accompanied by an increase in free or incompletely attached micronuclei, some of which looked like a single mitochondrion that was assembling dense particles for nuclear transition (fm1, MN1 and MN2). In the cytoplasm, it was rare to find orthodox mitochondria (fm2-fm16), which lacked either or both cristae and the limiting membrane (LMM). fm8 was evolving into an MN; between the nuclei (N2 and MN3; N3 and MN3), mitochondria were assembling dense particles (m1/1 and white arrowheads; m2/1 and fm3; red arrows, and opposite red, black and white arrowheads) to enlarge the nuclei, concomitantly accomplishing total nuclear merging. MN1 joined with N1 (opposite red arrowheads) by assembling dense particles from itself and N1 with the help of partially (fm6 and fm10) as well as completely (large and small red arrows; small red arrowheads) fragmented mitochondria. Mitochondrial aggregations of the particles led to the formation of dark strands (large red arrows) or threads (small red arrows), ERLs (white arrows) and electron-lucent structures (white arrowheads). Within N1, mitochondrial aggregations of dense particles were observed (NB1 and black arrowhead). NB: nuclear body (**C1-C8**). (**d**) In this LC3-deprived cell, nuclear fusions between these individually formed nuclei (N1-N5) were obstructed or delayed, and the mitochondrial assembly of dense particles for nuclear expansion were observed at the edge of N3 (opposite red arrowheads). Between nuclei (N3 and N4), most of the mitochondria had fragmented into particles, accompanied by the less fragmented fm1 and a heavily fragmented mitochondrion (black arrowhead) (**d1** and **d2**). (**d3** and **d4**) Adjacent to N1, there appeared a nascent nucleus (NN1), which showed a similar size to fm2, surrounded by fragmented mitochondria (such as fm3-fm6 and fm7/1) and enlarged by the assembly of mitochondria-derived dense particles at the edge (opposite red arrowheads). Between NN1 and N3, fragmented mitochondria dispersed into each other to form a circle (opposite red arrowheads), whose formation accelerated the joining of the two nuclei. (**d5** and **d6**) In the cytoplasm, mitochondria jointly assembled dense particles to form two groups (opposite red arrowheads), whose formation promoted the development of the enclosed organelles into an MN.

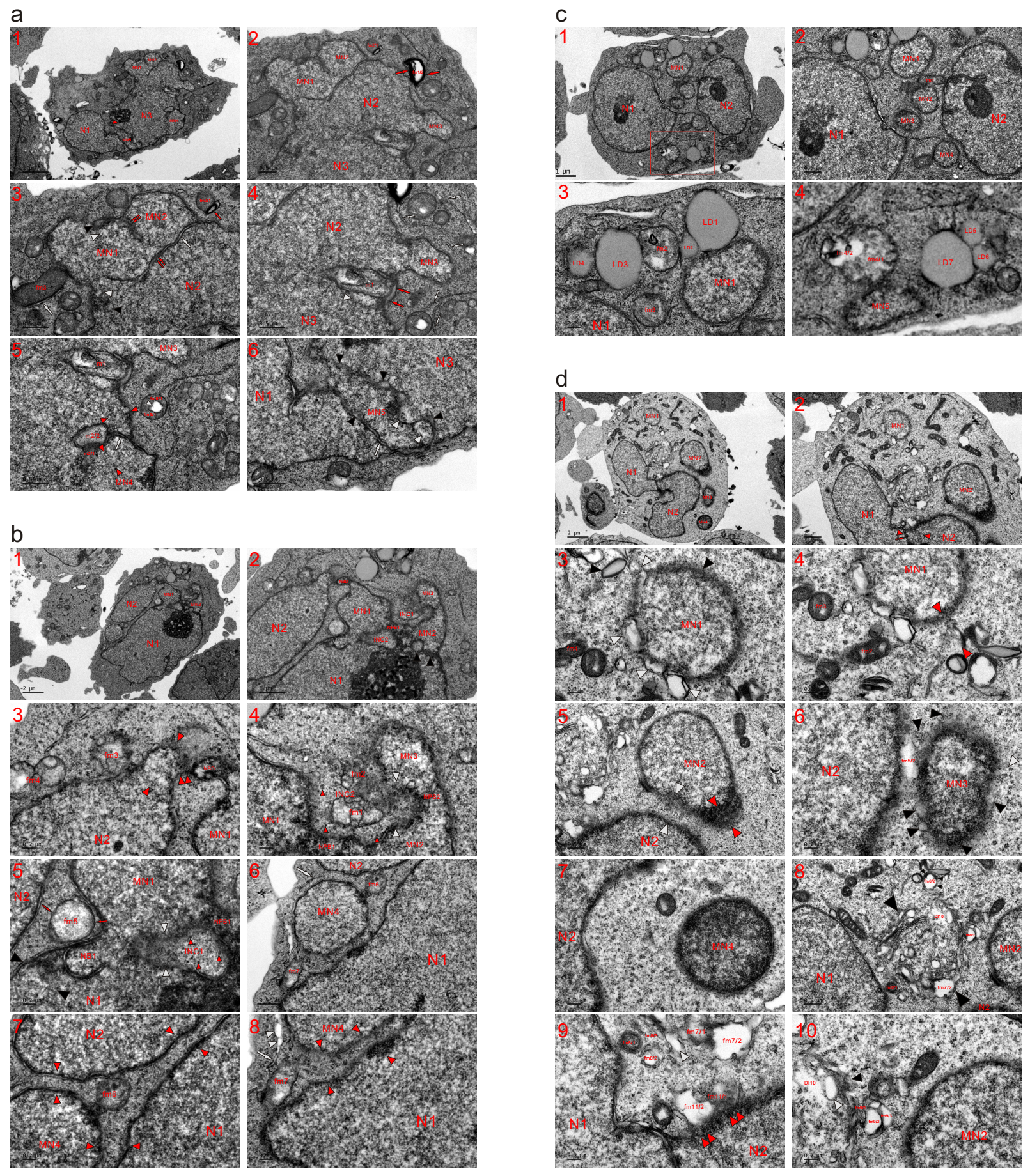

**Extended data Fig. 20. Knockdown of LC3 increased free micronuclei by promoting mitochondria-to-micronucleus transition.** TEM was performed on four K562 cells with (**c** and **d**) or without (**a** and **b**) the transfection of siRNA against LC3 at the 2 h time point, and the micrographs demonstrated that LC3 loss led to the formation of free micronuclei. (**a**) In addition to three large nuclei in this control cell, five attached micronuclei appeared (MN1-MN5), and incomplete mitochondria-to-nucleus transition or mitochondrial aggregations of dense particles were observed between nuclei (large white and double small red arrows) as well as at the nuclear edges (black and white arrowheads). The assembly of the particles heavily fragmented m1, which was concurrently incorporated into the nucleus (red arrow) between N2 and N3; immediately next to m1, an SDIB (white arrowhead) was disappearing via the assembly of dense particles. Mitochondrial congregations of dense particles formed ink strands (large red arrows), causing part of a mitochondrion to become either electron-transparent (fm1/2) or condensed (fm2/1). At the edge of condensed fm3, particle assembly formed an ERL (white arrow). Based on mitochondrial fragmentation, a number of ERLs appeared in the cytoplasm (white arrows) (**a1-a4**). (**a5**) Having achieved nuclear transition of its neighbouring counterparts (opposite red arrowheads and white arrow) caused a mitochondrion to become nuclearly localized, and the mitochondrion itself assembled the particles (m2/1 and m2/2) for nuclear development. At the nuclear edge, fm5 aggregated dense particles (fm5/1) to create a vesicle structure in its centre (fm5/2). (**a6**) Mitochondria assembled the particles to form MN5, which in turn joined the separately constructed N1 and N3, and incomplete mitochondria-to-nucleus transition was observed between nuclei (white arrow, black and white arrowheads). (**b**) The same cell shown in **Figure 4a** (**a1** and **a2**). Individually formed large nuclei (N1 and N2) were joined by mitochondria-to-nucleus transition, leading to both the formation and attachment of micronuclei (MN1-MN3). The mitochondrial assembly of dense particles to form NPB1 linked two micronuclei (MN1 and MN2) and simultaneously partitioned the cytoplasm to form the closed INC1 and open INC2. In both INCs, the incomplete nuclear conversion of mitochondria was observed (such as fm1 and fm2; opposite white and small red arrowheads). Mitochondrial aggregation of the particles occurred between N1 and MN2, which connected with MN3 through NPB2. An NBD was being evolved into the NPB by the mitochondrial assembly of dense particles (double red arrowheads), leading to partial fusion of N2 and MN1, which consequently acted as a linkage between two large nuclei (N1 and N2). fm3 developed a nuclear appearance (opposite red arrowheads), fm4 assembled the particles into N2 as well as the surroundings, and the external aggregation of dense particles at the periphery diluted fm5. The mitochondrial assembly of dense particles joined two nuclei (N1 and MN1) and formed a nuclear body (NB1), near which a fragmented mitochondrion had almost completed nuclear transition (opposite black arrowheads) (**b1-b5**). (**b6-b8**) Less fragmented mitochondria dispersed into nuclei (N1, N2 and MN4) to link the separately constructed MN4 with both N1 and N2. Between the nuclei, mitochondria completely fragmented into dense particles for nuclear development to complete nuclear merging (opposite red arrowheads). At the edge of MN4, ERLs (white arrows) and electron-lucent bodies (white arrowheads) temporarily formed during mitochondrial assembly of the particles to enlarge the micronucleus. (**c**) Same cell shown in **Figure 4a** (**a3** and **a4**). In addition to two large nuclei in this LC3-silenced cell, five free micronuclei (MN1-MN5) appeared, and electron-opaque fm1 dispersed into N1 and two micronuclei (MN1 and MN2). A whole mitochondrion (fm3) or part of other mitochondria (fm2 and fm4/1) had developed a nuclear appearance; lipid droplets (LD1-LD7) dispersed into nuclei (MN1 and MN5) and/or into each other (**c1-c4**). (**c4**) A high-magnification image of the inset in (**c1**). (**d**) Two large nuclei had recently accomplished nuclear fusion by the mitochondrial aggregation of dense particles (red arrow and opposite red arrowheads). In addition to the large nuclei, four micronuclei (MN1-MN4) existed in this LC3-deprived cell (**d1** and **d2**). (**d3** and **d4**) At the edge of MN1, the mitochondrial assembly of dense particles (opposite red, black and white arrowheads) enlarged the micronucleus, and somewhat distant from MN1, condensed mitochondria (fm1-fm4 and black arrowhead) were dispersed to continuously provide particles for nuclear growth. (**d5**) Mitochondrial aggregation of dense particles at the edge of MN2 (opposite red arrowheads), which was joined with N2 by completely fragmented mitochondria (opposite white arrowheads). (**d6**) MN3 was being expanded by the mitochondrial congregation of particles (black and white arrowheads) whose aggregation caused part of fm5 to become an electron-lucent body (fm5/2) between nuclei (N2 and MN3). After mitochondria assembled dense particles to enlarge both nuclei, MN3 eventually merged with N2 to become an attached micronucleus. (**d7**) Compared to the other three micronuclei, MN4 was denser and less advanced in nuclear development. (**d8-d10**) Among nuclei (N1, N2 and MN2), mitochondria jointly assembled dense particles to form a Golgi complex-like structure (opposite black arrowheads), and the external aggregation to the periphery (small black arrowheads) caused part of a mitochondrion to be electron-lucent (fm6/2). At the edge of the Golgi complex as well as within it, the assembly of dense particles separated mitochondria (such as fm7/1 and fm7/2; fm8/1-fm8/2; fm9/1-fm9/3), concomitantly forming electron-transparent bodies or structures (fm7/2, fm8/2, fm9/2, fm9/3, DI10 and white arrowheads). At the edge of N2, both dense fm11/1 and lucent fm11/1 assembled particles into the nucleus (double red arrowheads).

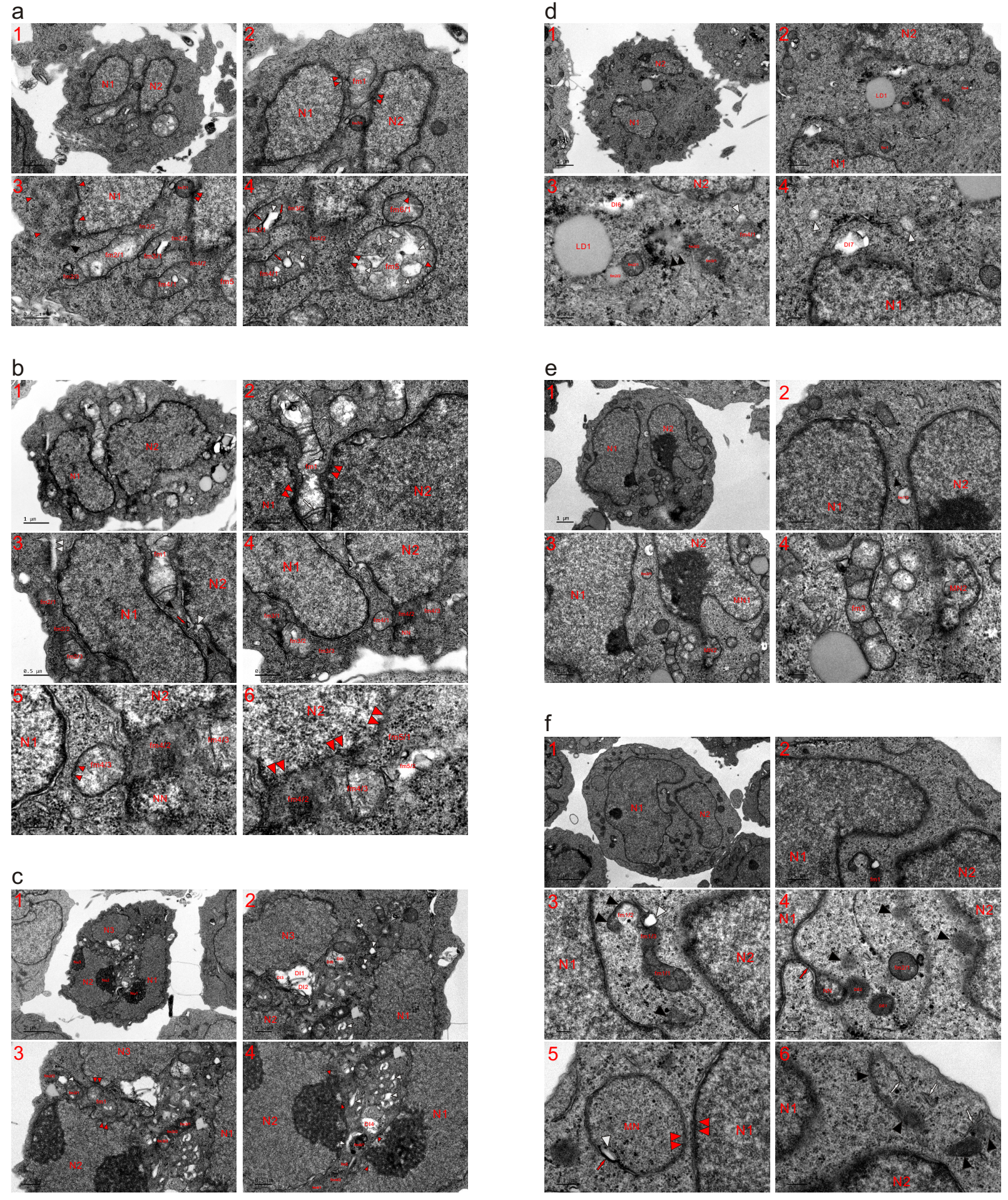

**Extended data Fig. 21. Knockdown of LC3 altered the pattern of mitochondria-to-nucleus transition.** TEM was performed on six K562 cells at the 2 h time point with (**d-f**) or without (**a-c**) knockdown of LC3, and the micrographs showed that LC3 deprivation altered the mitochondria-to-nucleus pattern by enhancing mitochondrial fragmentation. Compared to the mitochondria of control cells, mitochondria in cells with LC3 knockdown usually had decreased capability to form electron-lucent structures. (**a**) The same cell shown in **Figure 4b** (**b1** and **b2**). Having achieved nuclear transition of its neighbouring counterparts caused fm1 to localize between individually constructed nuclei (N1 and N2) and to disperse into the nuclei (double red arrowheads), while fm1 itself developed a nuclear appearance (**a1** and **a2**). (**a3** and **a4**) At the nuclear edges, partially (such as fm2-fm4) and completely (opposite red and black arrowheads) fragmented mitochondria diffused into the nuclei (fm2/2-fm4/2), promoting their growths; both fm5 and fm6/1 became particles (red arrowheads) whose internal aggregations (red arrows) created electron-lucent spots or structures (white arrowheads) in mitochondria. Between the nuclei, condensed fm7/1 assembled particles to fuse with N2 (double red arrowheads). (**b**) The nuclear conversion of mitochondria enlarged both N1 and N2, causing part of fm1 to become embedded between the nuclei; the external assembly of dense particles caused the mitochondria to fuse with either nucleus (double red arrowheads), and the internal aggregation formed an ERL (white arrow) within fm1 (**b1** and **b2**). (**b3** and **b4**) Adjacent to fm1 and between the nuclei, traces of mitochondrial fragmentation are displayed (red arrow, black and white arrowheads). At the edge of N1, the mitochondrial aggregation of dense particles created an electron-transparent rod-shaped structure (double white arrowheads) and had already dispersed (fm2/1, fm2/2, fm3/1 and fm3/3) or was dispersing (fm2/3 and fm3/2) the organelles for nuclear development to enlarge the nucleus. (**b5** and **b6**) Mitochondrial assembly of dense particles formed a nascent nucleus (NN), which merged with N2 by the dispersion of fm4 (fm4/2), and the less fragmented fm4/1 aggregated the particles into N1 (small red arrowheads). At the edge of N2, mitochondria (fm4/2, fm4/3 and fm5/1) fragmented into the nucleus (double red arrowheads) in the form of dense particles for nuclear expansion. (**c**) Three separately constructed nuclei (N1-N3) were joined by the mitochondrial assembly of dense particles for nuclear development, and mitochondrial aggregations of the particles led to the formation of electron-transparent or electron-dilute structures (DI1-DI6 and white arrowheads). fm1 aggregated dense particles to disperse into both N2 and N3, and their assembly separated either fm2 (fm2/1, fm2/2 and white arrowhead) or fm3 (fm3/1-fm3/3). Between N1 and N2, heavily fragmented mitochondria (fm4/1 and fm4/2; fm5; fm6/1 and white arrowhead) linked two nuclei; at the edges of nucleoli (Nu1 and Nu2), the mitochondrial assembly of particles (opposite red arrowheads) was observed (**c1-c4**). (**d**) The same cell shown in **Figure 4b** (**b3** and **b4**). Two nuclei were widely separated in this LC3-depleted cell, and almost all mitochondria between the nuclei were completely fragmented, with few less fragmented ones (fm1-fm4) either condensed or dispersed into dense particles (**d1** and **d2**). (**d3** and **d4**) The aggregation of dense particles separated fm2 into dense fm2/1 and fm2/2, which displayed the morphology of a lipid droplet (LD). Condensed mitochondria (fm3/1, fm3/2 and fm4/2) tended to directly disperse into the particles, whose assembly was likely to form an LD (double black arrowheads); these SDIBs were usually dilute but not electron-transparent (white arrowheads). At the nuclear edges, the external aggregations of dense particles at the peripheries allowed parts of the organelles to become electron lucent (DI6 and DI7). LD1: a large lipid droplet. (**e**) No large mitochondria were present between nuclei (N1 and N2), most of the organelles had completely fragmented, and a small number of less fragmented mitochondria were observed (fm1/2, fm2/1 and black arrowhead). At the edge of N2, an attached micronucleus (MN1) appeared, while MN2 was forming and looked like a mitochondrion assembling dense particles, whose aggregations attached MN2 to N2 and divided fm3 into small electron-opaque bodies. (**f**) In contrast to the mitochondria of the cell in (**c**), the aggregation of dense particles rarely caused the formation of electron-transparent structures following mitochondrial condensation (fm1/1, fm1/2, fm2/1, DE1 and DE2 and black arrowheads) between nuclei (N1 and N2). Through further assembly, the condensed organelles ultimately became particles, whose external aggregations (black arrowheads) caused parts of fm1 to become dilute (fm1/2) or lucent (white arrowhead). A mitochondrion assembled dense particles to look like a nascent nucleus (NN), which linked with N1 via a dark strand (red arrow) and attached to DE2 (**f1-f4**). (**f5**) At the edge of N1, mitochondria-derived particles developed the appearance of a micronucleus (MN), which was at the initial stage of nuclear formation, contained traces of mitochondrial fragmentation (red arrow and white arrowheads) and fused with N1 via the further assembly of dense particles (double red arrowheads), which existed at the edge of either MN or N1. (**f6**) The aggregations of dense particles dispersed mitochondria along with forming SDBs or dense structures (black arrowheads) and ERLs (white arrows), both of which eventually were becoming particles.

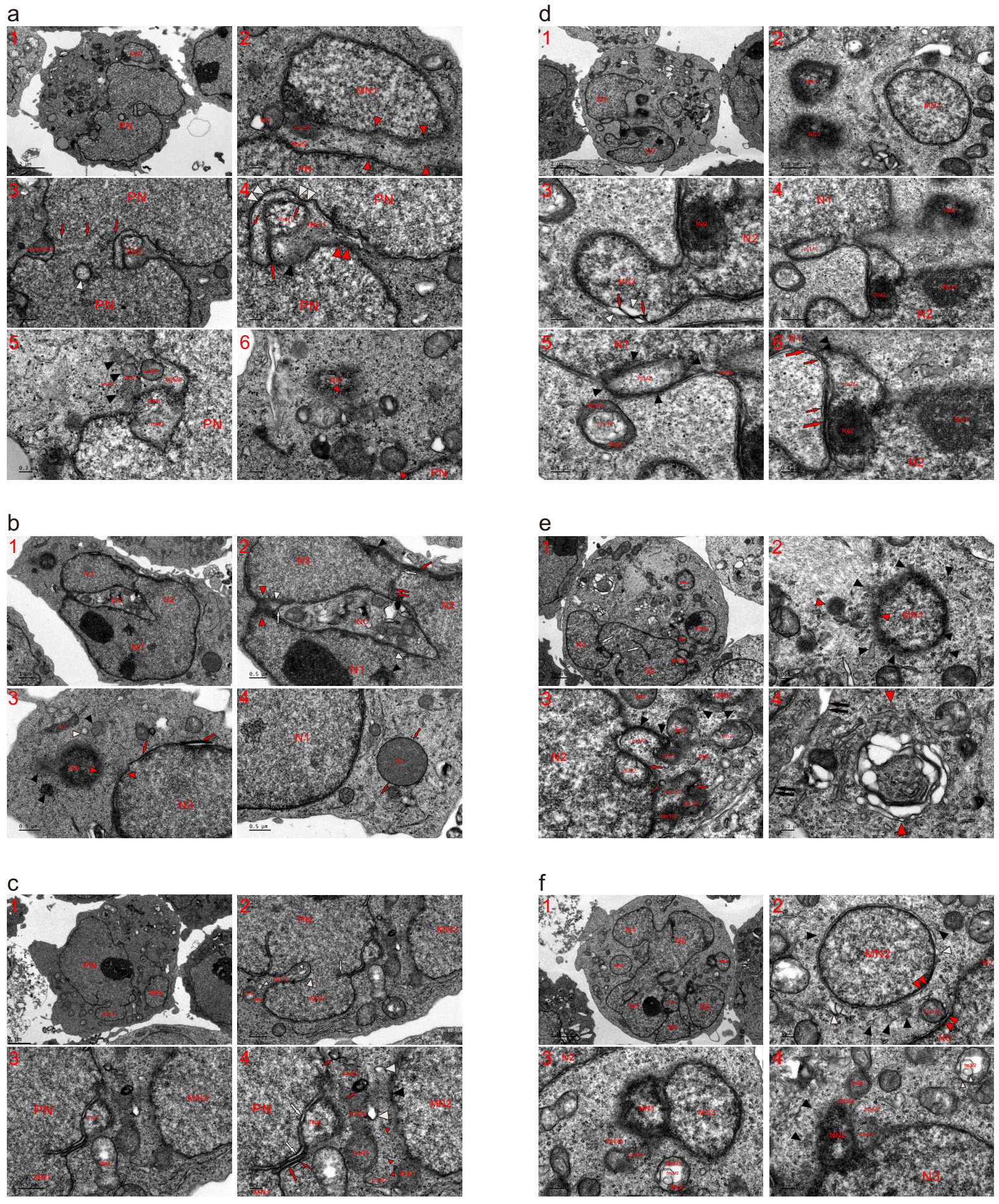

**Extended data Fig. 22. LC3 loss promoted mitochondria-to-micronucleus transition.** TEM was performed on three control K562 cells (**a-c**) and three LC3-silenced cells (**d-f**) at the 2 h time point, and the micrographs demonstrated that LC3 silencing promoted the formation of nascent nuclei and hindered the attachment of a small nucleus (or an MN) to the large nucleus. (**a**) The same cell shown in **Figure 4c** (**c1** and **c2**). MN1 was attached to PN by less fragmented (fm1, fm2/1 and fm2/2) and completely fragmented (opposite red arrowheads) mitochondria, and the aggregation of dense particles separated fm2 into dense fm2/1, which was being dispersed into particles and dilute fm2/2, which was already dispersed. (**a1** and **a2**). (**a3** and **a4**) The PN was formed from two individually formed nuclei, as shown by traces of nuclear fusion within the nucleus (three red arrows side by side); the combination of partial nuclear fusion and incomplete mitochondria-to-nucleus transition caused the partitioned cytoplasm to look like small intranuclear inclusions (INC1 and fm3/INC2) at the nuclear edge. Within PN, a mitochondrion assembled dense particles to form a dilute nuclear body (white arrowhead). INC1 was being closed by the mitochondrial assembly of dense particles for nuclear development (double red arrowheads), and within the INC, mitochondrial aggregation of the particles was observed (large and small red arrows; black arrowhead), causing part of a mitochondrion to exhibit a nuclear appearance (fm4/1). The external assembly of dense particles at the peripheries of fm4/1 and the PN caused the mitochondrion to fuse with the nucleus (double white arrowheads). (**a5**) Assembling dense particles of fm5 and its neighbouring counterparts (fm6 and fm7) formed an NPB, and their aggregations (NPB, fm5/1 and fm7/1) diluted parts of mitochondria (fm5/2, fm6/2 and fm7/2). (**a6**) Between PN and NN, condensed mitochondria were dispersed into dense particles for nuclear development (opposite red arrowheads) to enlarge both nuclei. (**b**) A combination of individually formed nuclei partitioned the cytoplasm to form a closed INC. Between N2 and N3, vestiges of nuclear fusion or traces of the mitochondrial assembly of dense particles were observed (double red and white arrowheads). At the nuclear edge and within N1, particle aggregation was observed (red arrows, black and white arrowheads); the INC was being sealed by the mitochondrial assembly of dense particles for nuclear development (white arrow, opposite red white arrowheads). Within the INC, mitochondria assembled particles to become partially, heavily or completely fragmented. At the edge of N3, mitochondria-derived particles assembled into the appearance of a nascent nucleus (NN), which was surrounded by fragmented organelles (fm1, black and white arrowheads). Between N3 and NN, traces mitochondrial fragmentation were observed (opposite red arrowheads). Adjacent to N1, there appeared a dense and globular body (B1), which seemed to be formed by the aggregation of mitochondria-derived particles or derived from a single condensed mitochondrion; through further assembly of the inside dense particles, the body tended to develop the appearance of an NN and transiently blackened the LMM (red arrows) (**b1-b4**). (**c**) Incomplete mitochondria-to-nucleus transition between the nuclei was observed (white arrow and arrowhead), and partial nuclear fusion attached MN1 to PN, concurrently partitioning the cytoplasm to form INC1. Within the INC and at its opening, traces of mitochondrial fragmentations were observed (fm1/1, fm1/2, small black and white arrowheads) (**c1** and **c2**). (**c3** and **c4**) MN2 was attached to the PN by mitochondria, which were partially (fm2, fm3 and fm4/1), heavily (fm4/2, fm4/3, fm5/1 and fm5/2; red arrows, black and white arrowheads) or completely (small red arrowheads) fragmented. Both fm2 and fm3 displayed a similar nuclear appearance, and the aggregation of dense particles caused fm2 to be incorporated into the PN and to look like an attached small MN. Mitochondrial assembly of the particles transiently formed dark strands (large red arrow), threads (small red arrow) and nuclear tubules (white arrows), and all these structures disappeared with the further aggregation of dense particles contained in the strand and thread. (**d**) The same cell shown in **Figure 4c** (**c3** and **c4**)**.** Two large nuclei (N1 and N2) were being joined by the mitochondrial aggregation of dense particles, and MN1 was widely separated from the large nuclei. Adjacent to N1, mitochondria-derived particles assembled to form nascent nuclei (NN1 and NN2) (**d1** and **d2**). (**d3**) MN2 was been separately constructed and attached to N2 following the mitochondrial assembly of dense particles to enlarge both nuclei (N2 and MN2), and at the edge of MN2, the mitochondrial assembly of dense particles (red arrows) formed electron-lucent structures (white arrowheads). (**d4-d6**) At the edge of N1, external mitochondrial assembly of the particles at the periphery (black arrowheads) included m1 itself in the nucleus, and part of the mitochondrion (m1/2) had adopted a nuclear appearance. In the cytoplasm and adjacent to m1, similar aggregation (fm1/1) caused the central part of fm1 to become dilute (fm1/2); within N2, mitochondria-derived particles assembled to form or enlarge nucleoli (Nu1 and Nu2). Unlike m1/2, m2/2 already exhibited a nuclear appearance and functioned as a part of the linkage between two large nuclei. Between the nuclei, mitochondria assembled dense particles to form an NPB (double black arrowheads), dark strands (large red arrows) and threads (small red arrows). (**e**) Nuclear tubules (white arrows) appeared in N2 and between two large nuclei, representing the mitochondrial aggregation of dense particles or incomplete mitochondria-to-nucleus transition; MN1 was far away from the large nuclei, and MN2 was adjacent to N2 (**e1**). (**e2**) Mitochondria surrounding MN1 were heavily or completely fragmented (opposite red and black arrowheads) to enlarge the micronucleus. (**e3**) At the edge of N2, the external aggregation of dense particles (m1/1, red arrow and black arrowheads) was incorporating part of the mitochondrion (m1/2) in N2, while m2/2 had already became part of the nucleus, and the dense counterpart of m2 had achieved nuclear transition and disappeared in N2. The assembly of dense particle particles separated fm1 (fm1/1-fm1/3, small and large red arrows) and caused the mitochondrion to fuse with N2 for nuclear development. At the edge of MN2, fragmented mitochondria dispersed into it in the form of dense particles for nuclear growth (black arrowheads). Along with the mitochondria-to-nucleus transition (such as m1 and fm2-fm5) to expand the nuclear sizes of both N2 and MN2, two individually constructed nuclei ultimately joined together. (**e4**) Near the plasma membrane (double black arrows), the mitochondrial assembly of dense particles together formed a group whose formation tended to initiate nuclear formation (opposite red arrowheads). (**f**) The combination of separately constructed nuclei (N1-N5 and MN1) led to nuclear atypia or irregularity in this LC3-silenced cell (**f1**). (**f2**) Heavily fragmented mitochondria appeared at the nuclear edges (black and white arrowheads), and less fragmented fm1 was dispersed into either the large nucleus (dense fm1/1 and double red arrowheads) or MN2 (double red arrowheads). (**f3** and **f4**) Mitochondria assembled dense particles to simultaneously enlarge MN3 and form a nascent nucleus (NN1), which consequently appeared to be fused with the micronucleus. At the edge of NN1, mitochondrial aggregation of the particles was observed (fm2/1), concurrently causing the dilute counterpart (fm2/2) to become part of the cytoplasm. At the edge of MN3, the aggregation of dense particles (fm3/1) diluted part of fm3 (white arrowheads), causing its central area to become lucent (fm3/2). At the edge of N2, mitochondria (fm4/1, fm5/2 and black arrowheads) assembled particles to form a nascent nucleus (NN2) or an NBD, which looked like a mitochondrion assembling dense particles.

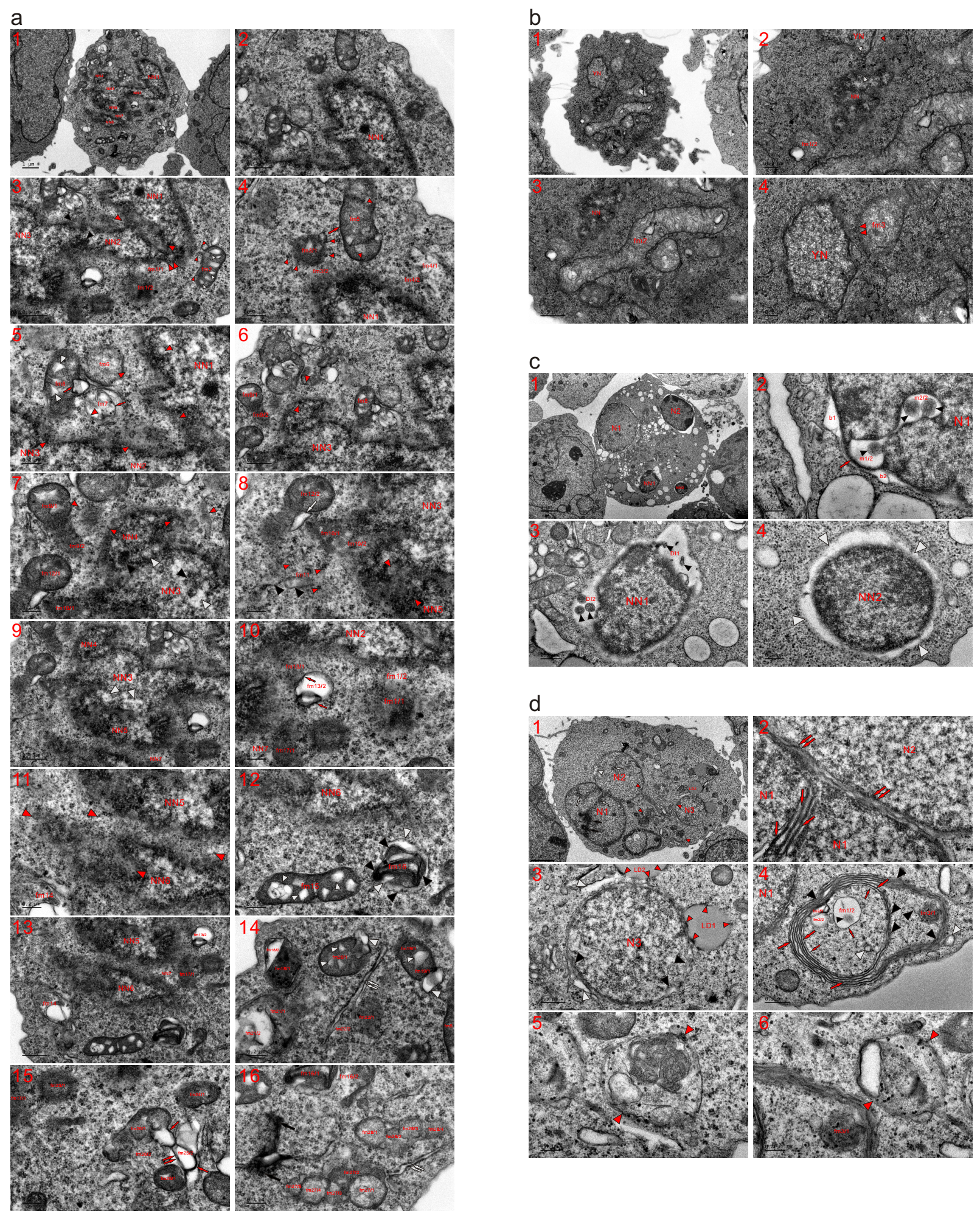

**Extended data Fig. 23. Knockdown of LC3 led to separate nuclear initiations and formations.** TEM was performed on two control K562 cells (**a** and **b**) and two LC3-depleted cells (**c** and **d**) at the 2 h time point, and the micrographs revealed that, in contrast to the control cells, in which nascent nuclei were connected by fragmented mitochondria, nuclear initiations and formations occurred individually in LC3-depraved cells. (**a**) The same cell shown in **Figure 4d** (**d1-d4**). Seven nascent nuclei (NN1-NN7) were enlarged, and NN7 was at a very early stage of nuclear initiation (NN7). Upon the mitochondria-to-nucleus transition, the nuclei became connected (**a1** and **a2**). (**a3**) The mitochondrial assembly of dense particles achieved partial fusion of NN1 and NN2 (double red arrowheads), accompanied by most mitochondria that were completely fragmented, with no nuclear transition between the nuclei (opposite red arrowheads). At the edge of NN2, fm1 assembled particles to disperse itself (fm1/1 and fm1/2). Between NN2 and NN3, mitochondria-derived dense particles promoted nuclear development (black arrowheads); their aggregation created electron-transparent or electron-dilute spots (small white arrowheads) in condensed fm2, which diffused into NN1 as well as the surroundings in the form of dense particles (small red arrowheads). (**a4**) Parts of the mitochondria dispersed into NN1 (fm3/2 and fm4/1), and less fragmented fm3/1 was becoming particles (small red arrowheads), whose aggregation transiently blackened the LMM of the organelle (red arrow) and caused fm5 to become dense particles (small red arrowheads). (**a5**) To enlarge the nascent nuclei (NN1-NN3), mitochondria (opposite red arrowheads) had to completely fragment into particles and consequently lost their morphology; mitochondrial origins of both fm6 and fm7 were recognized, and parts of the organelles (fm7, fm8 and opposite red arrowheads) dispersed into the nuclei (NN2 and NN3). The aggregation of dense particles intermediately formed dark threads (red arrows) and electron-lucent spots in fm8 (small white arrowheads). (**a6** and **a7**) Completely fragmented mitochondria appeared at the edge of NN4 (fm9/2 and opposite red arrowheads), and incomplete mitochondria-to-nucleus transition was observed (black and white arrowheads) between nuclei (NN3 and NN4) as well as within NN3. (**a8-a10**) Mitochondrial assembly of dense particles occurred between NN3 and NN5 (white arrowheads), and compared to NN3, NN5 was less advanced regarding nuclear development, with half of the nascent nucleus looking like mitochondria assembling dense particles (opposite red arrowheads). Part of fm10 (fm10/2) was completely fragmented into the nuclei, and the mitochondrial origin of fm10/1 was recognizable, while most of fm11 (black arrowheads) had dispersed into dense particles (small red arrowheads). fm12/1 diffused into the particles, whose aggregation in fm12/2 formed an ERL at the edge of the mitochondrion (white arrow). Along with the dispersion of dense fm13/1, electron-lucent fm13/2 and internal assembly of the particles (red arrows) appeared in the cytoplasm. (**a11** and **a12**) At the edge of NN6 and between nuclei (NN5 and NN6), mitochondria fragmented into dense particles for nuclear development (opposite red arrowheads), and fm14 was heavily fragmented and assembled particles into NN6. The further aggregation of dense particles in condensed fm15 formed electron-transparent spots (small white arrowheads) and caused fm16 to disperse (black and white arrowheads). (**a13**) The aggregation of the particles (fm17/1) caused the dilute part of a mitochondrion to look like a nascent nucleus (NN7), whose formation linked NN5 and NN6. (**a14**) The aggregation of dense particles separated the organelles into dense (fm18/1-fm21/1) and lucent or dilute (fm18/2, fm21/2 and large white arrowheads) parts, and their further assembly created transparent or dilute spots (small white arrowheads) in dense areas of the mitochondria (fm19/1 and fm20/1). Both dilute and dense parts of fm22 became particles, and mitochondria-derived dense particles formed a tubule structure (double white arrows). (**a15**) Condensed mitochondria (fm23/1-fm26/1) were dispersed to become particles whose internal aggregation formed dark threads (red arrows), and both dark threads and dilute fm26/2 diffused (double red arrows). (**a16**) Mitochondria assembled dense particles into ink masses, which were dispersed, and their assembly led the organelles to transiently form small mitochondria (fm27/1-fm27/5 and fm28/1-fm28/4) before diffusing into particles. (**b**) Adjacent to YN, a nascent nucleus (NN) was formed by the mitochondrial assembly of dense particles, and almost all mitochondria (fm1/2 and opposite red arrowheads) surrounding the NN had fragmented into particles, which were promoting nuclear development to link two nuclei (opposite white arrowheads). Near the NN, there appeared an elongated mitochondrion (fm2), which was being dispersed via the internal and external aggregation of dense particles. Next to the YN, fm3 developed a similar nuclear density and diffused to fuse with the nucleus (double red arrowheads) via external particle assembly (**b1-b4**). (**c**) In this LC3-depleted cell, four nuclei (large N1 and N2; small NN1 and NN2) were individually constructed. The external aggregation of dense particles to the periphery (the external assembly) caused part of the mitochondria to become nuclear bubbles (b1 and b2), and the external assembly (red arrow) incorporated an electron-transparent part of a mitochondrion into N1 (m1/2). Along with its nuclear transition and its neighbouring counterparts, m2/2 became a nuclear-localized mitochondrion, and the internal aggregation of dense particles was displayed in both m1/2 and m2/2 (small black arrowheads) (**c1** and **c2**). (**c3**) At the edge of NN1, external mitochondrial aggregation of dense particles to NN1 as well as the surroundings led to the formation of electron-lucent structures at the nuclear edge (DI1 and DI2), and within the DIs, the internal assembly of the particles was observed (small black arrowheads). (**c4**) The aggregation of dense particles formed a lunar halo structure at the edge of NN2 (white arrowheads). (**d**) Three nuclei (N1-N3) were separately constructed in the cell, and in both N1 and N2, incomplete mitochondria-to-nucleus transition was observed (white and black arrows, and white arrowheads) (**d1**). (**d2**) Between N1 and N2, mitochondria-derived dense particles assembled for nuclear development to merge two nuclei (double red arrows). Within N1, the dispersion of dark strands was observed (red arrows). (**d3**) At the edge of N3, incomplete mitochondria-to-nucleus transition was observed (black arrowheads), and the mitochondria assembled dense particles into the nucleus and the surroundings to become electron-transparent (white arrowheads). LD1 aggregated the particles to fuse with N3, and both lipid droplets (LD1 and LD2) assembled dense particles into the nucleus as well as the surroundings (small red arrowheads). (**d4**) Adjacent to N1, mitochondrial aggregations of dense particles (fm3/1 and black arrowheads) appeared whose assembly formed dark strands (large red arrows) or threads (small red arrows) and caused some of the organelles to become electron-lucent (fm1/2, fm2/2, fm2/3 and white arrowheads). fm1/2 exhibited the external (small red arrow) and internal (small black arrowhead) assembly of dense particles to disperse, and both fm2/2 and fm2/3 eventually diffused into the particles. Dark strands combined into the onion-like structure, and the mitochondrial aggregation of the particles into circles benefited mitochondria-derived dense particles for nuclear development. (**d5** and **d6**) In the cytoplasm, mitochondria assembled dense particles to form groups (opposite red arrowheads and **d1**), promoting the nuclear transition of the inside fragmented mitochondria to form small nuclei or micronuclei.
